# 4D structural biology: quantitative dynamics in the eukaryotic RNA exosome complex

**DOI:** 10.1101/2024.01.28.577622

**Authors:** Jobst Liebau, Daniela Lazzaretti, Torben Fürtges, Anna Bichler, Michael Pilsl, Till Rudack, Remco Sprangers

## Abstract

Molecular machines play pivotal roles in all biological processes. Most structural methods, however, are unable to directly probe molecular motions. Here, we demonstrate that dedicated NMR experiments can provide quantitative insights into functionally important dynamic regions in very large asymmetric protein complexes. We establish this for the 410 kDa eukaryotic RNA exosome complex that contains ten distinct protein chains. Methyl-group and fluorine NMR experiments reveal site-specific interactions among subunits and with an RNA substrate. Furthermore, we extract quantitative insights into conformational changes within the complex in response to substrate and subunit binding for regions that are invisible in static cryo-EM and crystal structures. In particular, we identified a flexible plug region that can block an aberrant route of RNA towards the active site. Based on molecular dynamics simulations and NMR data we provide a model that shows how the flexible plug is structured in the open and closed conformations. Our work thus demonstrates that a combination of state-of-the-art structural biology methods can provide quantitative insights into large molecular machines that go significantly beyond the well-resolved and static images of biomolecular complexes, thereby adding the time domain into structural biology.

## Main Text

Protein dynamics are tightly coupled with function (*1–3*). Nuclear magnetic resonance (NMR) methods are particularly well suited to study dynamic processes in solution, at quasi atomic resolution and on a wide range of timescales (*4–6*). Recent advances in sample preparation combined with NMR pulse-sequence and hardware design have made complexes over 100 kDa accessible to detailed solution NMR studies (*7–11*). This thus opens ample opportunities where NMR methods can complement static structural information obtained by e.g. single particle cryo-electron microscopy (cryo-EM) or *in silico* tools (*12*, *13*).

Here, we study the eukaryotic RNA exosome, an essential 3′–5′ ribonuclease complex (**Fig. 1A**). In the cytoplasm, the exosome is involved in the canonical turnover of mRNA and in mRNA quality control; in the nucleus the complex degrades and processes a wide variety of RNA substrates (*14*, *15*). The exosome is a modular molecular machine that consists of an inert, nonameric core (Exo9; 300 kDa). This core contains an essential central channel that is formed by six distinct RNase PH-like subunits (Rrp41, Rrp45, Rrp43, Rrp46, Mtr3, Rrp42) and a substrate entrance pore that is formed by three cap subunits (Csl4, Rrp4, Rrp40) that contain RNA binding domains (*16*). Rrp41 and Rrp45 recruit the catalytic subunit Rrp44 (Dis3 in humans) (*17*) to assemble the catalytically active decameric complex (Exo10; 410 kDa). Within Rrp44, the RNB domain harbors processive exonucleolytic activity, while the PIN domain can hydrolyse RNA in an endonucleolytic manner (*18*, *19*) During catalysis, the 3′ end of a single-stranded RNA substrate is recruited by the cap subunits, threaded through the channel (*20*) and is finally presented to Rrp44 (**Fig. 1A**). Several compartment-specific co-factors associate with the complex to convey substrate specificity (*18*, *21*). Additionally, the exosome can recruit Rrp6, a second catalytic subunit that harbors distributive exonucleolytic activity (*21*, *22*). Mutations in the exosome complex have been linked to multiple human diseases, underscoring its central functional importance (*23*). In the past, static structures of the human (*18*, *24*, *25*) and yeast (*22*, *26–32*) exosome complexes have been reported that reveal its subunit organization and RNA interactions.

**Figure 1:**
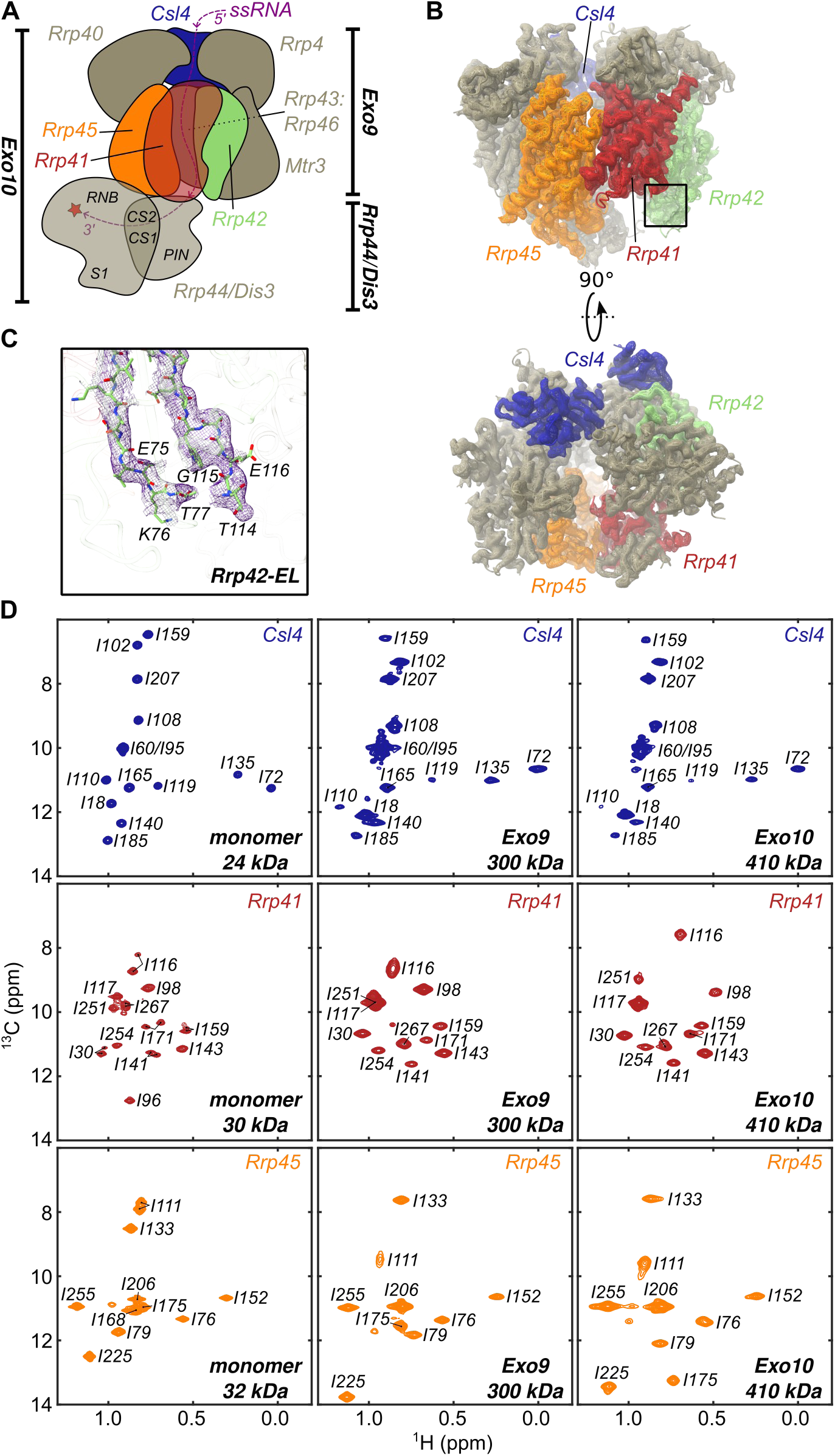
Structure of ctExo9 and assignment of NMR spectra. (**A**) Schematic depiction of Exo10. Throughout the manuscript the Csl4, Rrp41, Rrp42 and Rrp45 subunits are colored blue, red, green and orange, respectively. The individual Rrp44 domains (PIN; pilT N-terminal, CS: cold-shock, RNB: RNA binding and S1) are labeled and the exonucleolytic site is highlighted with a star. The path of the RNA towards the active site is indicated with a purple line. (**B**) Side view (top) and top view (bottom) of the ctExo9 cryo-EM density map. (**C**) Zoom around the boxed region in panel **B**, top, that displays the cryo-EM density around the invisible section of the extended loop in Rrp42 (Rrp42 -EL). (**D**) Ile-δ1 region of methyl-TROSY spectra for Csl4, Rrp41 and Rrp45 in the monomeric form (column 1) and when reconstituted into Exo9 (column 2) or Exo10 (column 3). Resonance assignments are indicated.

### Static structures of the ctExo9 complex

Here, we complement this structural information and determined the structure of Exo9 from the eukaryotic thermophile *Chaetomium thermophilum* (ctExo9) by X-ray crystallography to 3.8 Å resolution (**fig. S1B, C, table S1A**) and by cryo-EM to 3.2 Å resolution (**Fig. 1B**, **fig. S1A, C, fig. S2, table S1B**). The general architecture of ctExo9 is identical to that of yeast and human exosome complexes (*18*, *22*, *24–31*). Despite this wealth of structural information, several disordered regions are invisible in these structures. In particular, an extended loop region in Rrp42 (Rrp42-EL) and the entry loop in Rrp41 are largely unresolved, while the shorter exit loop of Rrp41 is only partially visible (**Fig. 1C, fig. S1A, D, E**). A priori, these “invisible” regions cannot be considered functionally unimportant, as disordered regions are often directly involved in biological function (*3*, *33*, *34*).

### NMR assignments in the exosome complex

To obtain insights complementary to the static structures, we turned to methyl-based NMR spectroscopic methods (*10*). Such approaches have been successfully applied to large, highly symmetric protein assemblies with molecular weights of up to 1 MDa (*35–38*) and to single-chain proteins of up to 100 kDa (*39*). In that light, the eukaryotic exosome complex is significantly more challenging to study, as it contains ten distinct protein chains with a total molecular weight of almost half a megadalton. To render the exosome complex visible to NMR spectroscopy, we employed a labeling scheme, in which one subunit at a time was labeled with NMR-active Ile-δ1[^13^CH_3_] and Met-ε1[^13^CH_3_] methyl groups in an otherwise fully deuterated background (‘IM-labeling’) (**fig. S3**). Methyl resonance assignments were obtained by exploiting a divide-and-conquer strategy, where we first assigned resonances in the monomeric subunits Csl4, Rrp41 and Rrp45 (**Fig. 1D**, **fig. S4**). These assignments were then transferred to the Exo9 and Exo10 complexes, assisted by numerous point mutants (**table S6C**, exemplified in **fig. S5**). The assignment completion of the Ile-δ1 resonances was close to 90% (**table S2**) providing a set of NMR probes that can report on interactions and dynamics and that are well distributed over the complex.

### Interactions between Exo9 and Rrp44

Based on chemical shift perturbations (CSPs) site-specific insights into intermolecular interactions can be obtained. Chemical shifts of a number of resonances in the ring subunits Rrp41 and Rrp45 differ significantly between the Exo9 and Exo10 complexes, whereas resonances in Csl4 were unaffected by the addition of Rrp44 (**Fig. 1D**, **fig. S6**). These observations are in agreement with existing structural information for yeast and human exosomes that show that Rrp44 is recruited to the Exo9 complex by Rrp41 and Rrp45 (*18*, *24*, *27–30*). Our data thus reveal that interactions between ctExo9 and ctRrp44 are conserved and that methyl-TROSY methods can be exploited to identify interaction interfaces in large asymmetric eukaryotic assemblies.

The methyl-TROSY methods that we deployed are “blind” in regions that are devoid of Ile or Met residues. To also investigate such regions, we turned to ^19^F NMR methods that were recently shown to be excellent tools to study interactions and dynamics on a broad range of timescales (*40–43*), even for larger complexes (*44–47*). First, we employed amber codon suppression to introduce a 4-trifluoromethyl-L-phenylalanine (tfmF) into Rrp41 at position D113 (Rrp41^D113tfmF^). Based on our structures, this position is located next to a partially structured loop (‘exit loop’) that faces the Rrp44 interaction interface of Exo9 and that lines the exit site of the RNA channel (**fig. S1A, D**). Upon addition of Rrp44 the resonance of Rrp41^D113tfmF^ shifts, demonstrating its spatial proximity to Rrp44 (**fig. S7A**). Second, we introduced a tfmF label at position Q86 in Rrp41 (Rrp41^Q86tfmF^) that is located in an extended loop close to the cap subunits and not visible in any of the structures (‘entry loop’, **fig. S1A, D**). This resonance is not affected by the addition of Rrp44, in agreement with a remote location of the entry loop from Rrp44 (**fig. S7B**). To probe if the “invisible” entry loop approaches the entry site of the RNA channel, we assembled an exosome complex in which the cap subunit Csl4 was labeled with a paramagnetic 2,2,6,6-tetramethylpiperidine-1-oxyl (TEMPO) spin-label at position E130 (Csl4^C122S, E130C-TEMPO^, **fig. S1C**). Rrp41^Q86tfmF^ proved to be too remote to be affected by the Csl4 spin-label. However, Rrp41^G71tfmF^, for which the flourine label is located in the center of the entry loop, displayed fluorine paramagnetic relaxation enhancements (PREs, Γ) that are a direct reporter of the distance between the Csl4 spin-label and Rrp41^G71tfmF^. The spin-label gives rise to enhanced R_1_ and R_2_ relaxation rates, establishing that the Rrp41 entry loop is located at the entry site of the RNA channel (**fig. S8, table S3A**). To probe if the entry (Rrp41^Q86tfmF^) and exit (Rrp41^D113tfmF^) loops undergo motions on the micro-to millisecond timescale we measured fluorine CPMG relaxation dispersion experiments (**fig. S9**) that reveal no signs of chemical exchange, indicating that both loops move on a fast (≪ms) timescale in solution.

### RNA threads through the exosome channel

To investigate interactions between the exosome and RNA in solution, we first assessed methyl CSPs in the subunit-specific IM-labeled Exo9 and Exo10 complexes. These data reveal that Csl4, Rrp41 and Rrp45 all interact with the substrate (**Fig. 2A**). In Csl4, CSPs are most pronounced in the S1 domain (residues 98-178) indicating direct interactions with the linear RNA substrate (**Fig. 2C, D**). This is expected since S1 domains have been implicated in RNA binding (*48*, *49*). In Rrp41 and Rrp45, resonances of residues that line the channel are affected by RNA (**Fig. 2A, C**), consistent with RNA being threaded through the channel (**Fig. 2D**). Additionally, ^19^F NMR data confirm the involvement of both the Rrp41 entry and exit loops in RNA interactions (**Fig. 2B**, **fig. S8C**). Moreover, for the entry loop, we observe a reduction of the PRE effect that is caused by Csl4^C122S, E130C-TEMPO^ upon addition of RNA (**fig. S8A, B**). This indicates that the dynamic entry loop is displaced away from Csl4 when RNA enters the exosome barrel (**fig. S8D**). Based on previous RNA interaction studies and alignments of the ctRrp41 and ctRrp45 sequences with corresponding archaeal, yeast and human sequences (**fig. S10**) we establish that the RNA coordination via positively charged residues inside the channel is conserved among those species (*17*, *50*).

**Figure 2:**
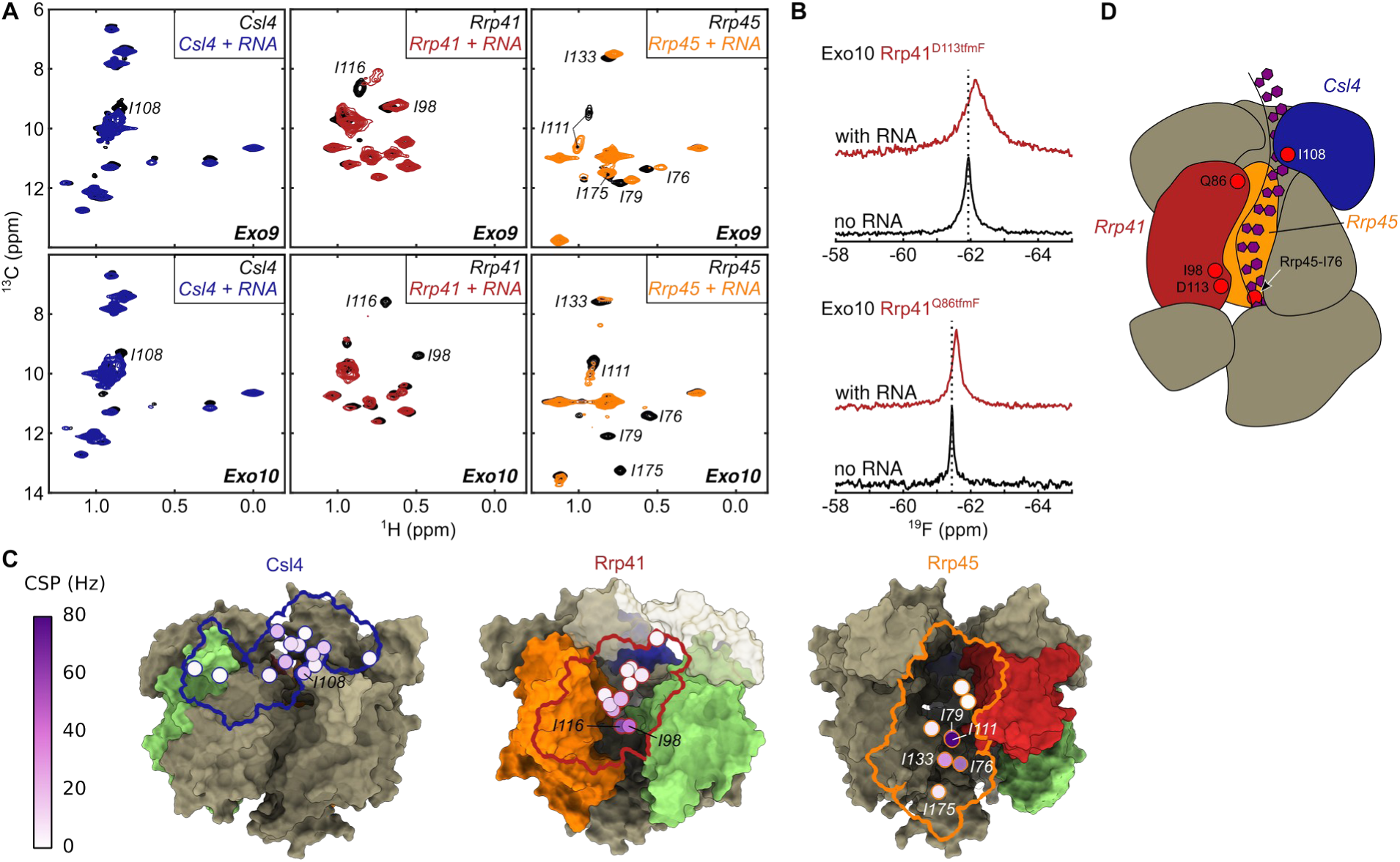
RNA interaction in the exosome. (**A**) Ile-δ1 region of methyl-TROSY spectra in the absence (black) and presence of RNA for Csl4 (blue), Rrp41 (red) and Rrp45 (orange) reconstituted into Exo9 (top) and Exo10 (bottom). (**B**) 1D ^19^F spectrum of Rrp41^D113tfmF^ (top; exit loop) and Rrp41^Q86tfmF^ (bottom; entry loop) reconstituted into Exo10 with (red) and without (black) RNA. The dashed line indicates the center of the resonance for Exo10. Both loops are affected by RNA interactions (**C**) RNA-induced CSPs of Ile-δ1 for Exo9 plotted onto the cryo-EM structure with the same coloring scheme as in panel **A**. Rrp42 is in green. For clarity only the outline of the subunit that is NMR active is shown. (**D**) Schematic depiction of Exo10 showing a tentative RNA path through the exosome channel. The coloring scheme is as in panel **A**. Rrp42 is omitted for clarity.

### RNA displaces a channel exit loop

Next, we investigated the dynamics and function of an extended loop region in the Rrp42 subunit, Rrp42-EL (**fig. S11A**), that is unresolved and thus invisible in both the X-ray and cryo-EM structure (**Fig. 1C**). To obtain insights into the location of Rrp42-EL in the exosome complex, we engineered a double mutant, Rrp42^C59S, A106C^, in which the single wild-type Cys residue is replaced by a Ser residue (C59S) and a new Cys residue is incorporated into Rrp42-EL (A106C). Subsequently, we attached a TEMPO spin-label to Rrp42-EL (Rrp42^C59S, A106C-TEMPO^) and reconstituted this subunit together with IM-labeled Csl4, Rrp41 or Rrp45 into Exo9 and Exo10 complexes. Csl4 resonances are not affected by the spin-label (**Fig. 3A, E**), indicating that Rrp42-EL does not approach the cap subunit Csl4. In contrast, a number of Rrp41 and Rrp45 resonances display substantial PRE effects (I_para_/I_dia_ < 1) in Exo9 and Exo10 (**Fig. 4B**, **C, F, G****, fig. S12**). The affected residues face the exit site of the Exo9 channel, and PREs are stronger in the Exo10 complex than in the Exo9 complex indicating that the conformation of invisible Rrp42-EL is affected by Rrp44. In the presence of RNA substrate, PRE effects in Rrp45 are obliterated (**Fig. 4D**, **fig. S12**), indicating that RNA displaces the loop away from Rrp45. Based on that, we conclude that Rrp42-EL adopts two conformations: one, in which it is proximal to the channel and which is stabilized by Rrp44 (‘closed’) and another, in which it is distant from the channel and which is preferred in the presence of RNA (‘open’).

**Figure 3:**
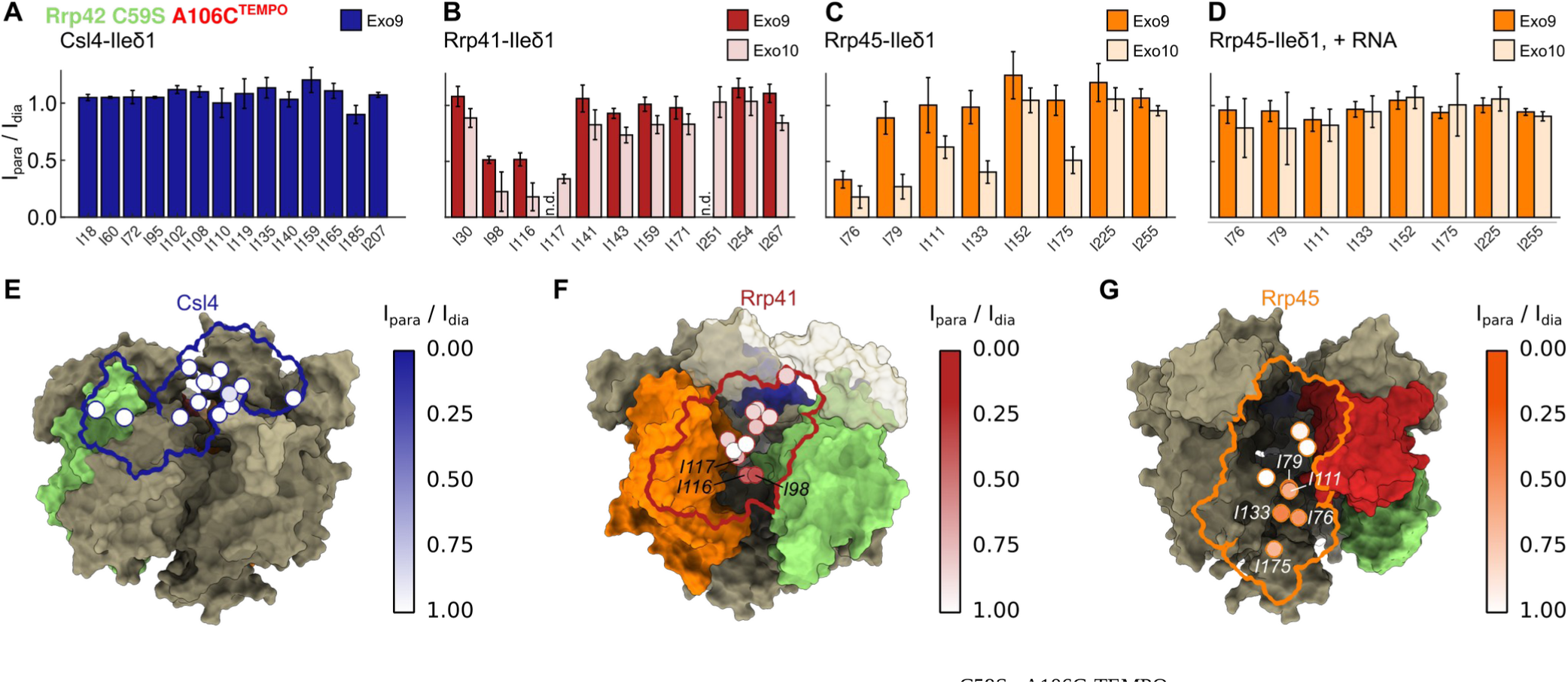
Localization of Rrp42-EL. PRE effects of Rrp42^C59S, A106C-TEMPO^ on Ile-δ1 of (**A**) Csl4 in Exo9, (**B**) Rrp41 in Exo9 (red) and Exo10 (light red), (**C**) Rrp45 in Exo9 (orange) and Exo10 (light orange) without RNA and (**D**) Rrp45 in Exo9 (orange) and Exo10 (light orange) with RNA. n.d.: value not determined due to signal overlap. Ile-δ1 PREs of (**E**) Csl4 (in Exo9), (**F**) Rrp41 (in Exo10) and (**G**) Rrp45 (in Exo10) plotted onto the cryo-EM structure. Csl4 is shown in blue, Rrp41 in red, Rrp45 in orange and Rrp42 in green. Note, that Rrp42-EL is not visible in the structure (see Fig. 1C). For clarity only the outline of the subunit that is NMR active is shown.

**Figure 4:**
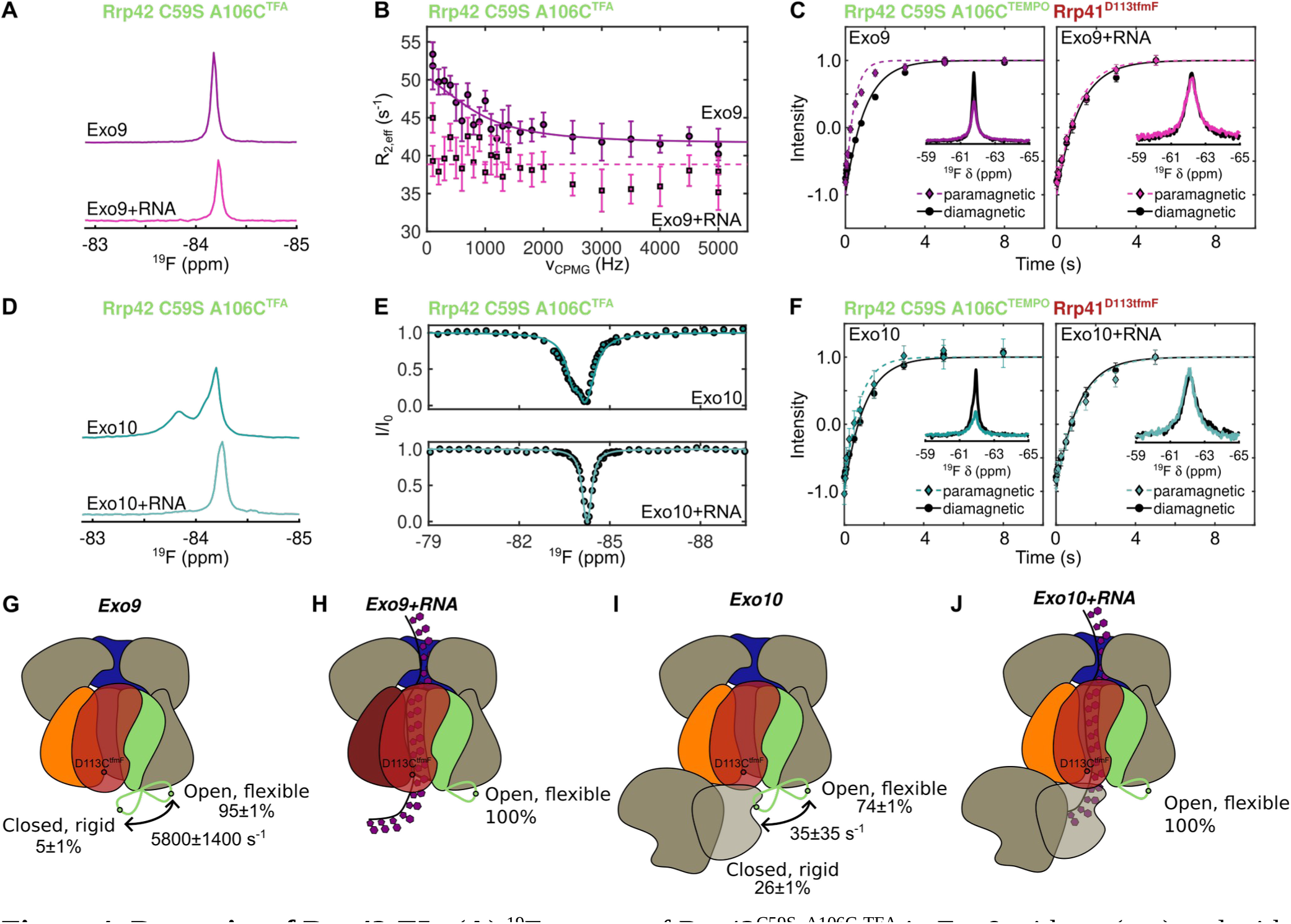
Dynamics of Rrp42-EL. (**A**) ^19^F spectra of Rrp42^C59S, A106C-TFA^ in Exo9 without (top) and with (bottom) RNA and (**B**) corresponding RD profiles. (**C**) Inversion recovery experiments (curves) and ^19^F 1D spectra (insets) for Rrp41^D113tfmF^ in Exo9 Rrp42^C59S, A106C-TEMPO^ without (left) and with (right) RNA for paramagnetic (in color) and diamagnetic (in black) samples. (**D**) ^19^F spectra of Rrp42^C59S, A106C-TFA^ in Exo10 without (top) and with (bottom) RNA and (**E**) corresponding CEST profiles, acquired for B_1_ = 25 Hz. (**F**) Inversion recovery experiments (curves) and ^19^F 1D spectra (insets) for Rrp41^D113tfmF^ in Exo10 Rrp42^C59S, A106C-TEMPO^ without (left) and with (right) RNA for paramagnetic (in color) and diamagnetic (in black) samples. (**G-J**) Schematic depiction of Rrp42-EL dynamics in Exo9 (**G**) without and (**H**) with RNA and in Exo10 (**I**) without and (**J**) with RNA. Rrp41 is in red, Rrp45 in orange, Csl4 in blue and Rrp42 in green. The green loop depicts Rrp42-EL, the green dot A106C^TFA/TEMPO^ and the red dot Rrp41^D113tfmF^. Fits for panels **C** and **F** were obtained using **Eq. S1**. Fits for panels **B** and **E** were obtained using the model described in materials and methods.

### Rrp44 and RNA modulate Rrp42-EL dynamics

To obtain direct insights into the dynamics of the invisible Rrp42-EL, we labeled A106C with 3-Bromo-1,1,1-trifluoro-acetone (BTFA) to form Rrp42^C59S, A106C-TFA^ (**fig. S13**), which retains ribonucleolytic activity of the Exo10 complex (**fig. S14**). Within the Exo9 complex, Rrp42^C59S, A106C-TFA^ displays one fluorine resonance (**Fig. 4A**); however, CPMG relaxation dispersion (**Fig. 4B**, **fig. S15B**) and chemical exchange saturation transfer (CEST) (**fig. S15A**) measurements reveal the presence of a second, minor conformation. Addition of single-stranded RNA results in a minor shift of the fluorine resonance frequency (**Fig. 4A**) and restricts the motions of the loop considerably as evidenced by an attenuated relaxation dispersion profile and a symmetric CEST dip (**Fig. 4B**, **fig. S15C, D**). To obtain additional information on the localization and dynamics of this invisible loop, we attached a spin-label to position A106C (Rrp42^C59S, A106C-TEMPO^) and determined its PRE effect on a ^19^F resonance in Rrp41^D113tfmF^ (**Fig. 4C**, **table S3B**). In the absence of RNA, sizable PRE effects are visible for R_1_ and R_2_ relaxation rates, indicating that Rrp42-EL and the Rrp41 exit loop come within less than ∼10 Å from each other. This is in agreement with the complementary methyl-TROSY data (**Fig. 3B**), where spin-labeled Rrp42-EL caused PRE effects close to the exit loop of Rrp41 (e.g. I98 and I116). Upon addition of substrate, the fluorine PRE effects are abolished (**Fig. 4C, H**), which implies that the Rrp41 exit loop and the invisible Rrp42-EL move apart.

We next turned to the Exo10 complex, in which the fluorine label in Rrp42-EL displays a second downfield-shifted resonance implying the formation of a second, long-lived conformation (**Fig. 4D**). This second conformation is induced by the C-terminal RNB-S1 domains of Rrp44, as Exo10 complexes that only contain the Rrp44 PIN domain or the PIN domain plus the two cold-shock (CS) domains fail to stabilize the second conformation (**fig. S16**). Structurally, a direct interaction between Rrp42-EL and the Rrp44-RNB-S1 domains is unlikely based on known structures of the human and *S. cerevisiae* Exo10 complexes (**fig. S17**). It is, however, plausible that the CS domains are brought into close spatial proximity of Rrp42-EL when Rrp44-RNB-S1 interacts with Exo9 as also indicated by a model of ctExo10 (see below).

The dynamics of Rrp42-EL in the Exo10 complex can be directly probed using CEST (**Fig. 4E**, **fig. S15E**) and longitudinal exchange (EXSY) (**fig. S15F**) experiments. Since EXSY experiments exclusively probe motions on the slow NMR timescale, we conclude that the dynamics of Rrp42-EL are significantly slowed down by Rrp44. ^19^F PRE effects show that Rrp42-EL in the Exo10 complex is still in close proximity to the Rrp41 exit loop (**Fig. 4F**), which corroborates the methyl-TROSY data (**Fig. 3B**). Upon addition of RNA to Exo10, the downshifted ^19^F resonance and PRE effects disappear and there are no indications for motions in the CEST profiles (**Fig. 4D-F**, **J, fig. S15G, H**) implying that Rrp42-EL is fully in the open state. Rrp42-EL likely adopts multiple inter-converting conformations in the open state as evidenced by a weak relaxation dispersion profile of Exo10 Rrp42^C59S, A106C-TFA^ in the presence of RNA (**fig. S15H**) and a shoulder of the open-state resonance in the absence of RNA (**Fig. 4D**, **fig. S15F, inset**).

### Quantification of Rrp42-EL dynamics

Interestingly, we observe that upon addition of Rrp44 to the apo Exo9 complex Γ _2_ rates are enhanced (spectra in **Fig. 4C, F**; indicating that Rrp42-EL moved towards Rrp41 upon formation of the Exo10 complex) while Γ_1_ rates are diminished (inversion recovery plots in **Fig 4C**, **F, table S3B**; indicating that Rrp42-EL moved away from Rrp41 upon formation of the Exo10 complex). This apparent contradiction can be explained by the differential dependence of Γ_1_ and Γ_2_ on fast timescale motions. As noted before (*51*, *52*) decreased order parameters (S^2^) and increased internal motions (τ_i_) can result in enhanced Γ_1_ rates, whereas Γ_2_ rates are largely unaffected by motions that are faster than the rotational correlation time (**Eq. S9, fig. S19**). Our data thus imply that the invisible Rrp42-EL is more rigid in the closed conformation, that is more prominently populated in the Exo10 complex, than in the ensemble of open conformations, which is mainly populated in the Exo9 complex.

To quantitatively assess the dynamics of the open-closed equilibrium and to obtain insights into order parameters of Rrp42-EL in the two states, we globally fitted a two-site exchange model to the dynamics experiments (CEST, RD, EXSY) of apo Exo9 and Exo10, and to ^19^F PRE experiments in the Exo9 and Exo10 complexes in the absence and presence of RNA (**fig. S15**, **fig. S18**, **fig. S20, table S4**). We assumed that the chemical shifts, local correlation times and order parameters of the open and closed states are the same in the Exo9 and Exo10 complexes. The analysis revealed that in Exo9, the invisible Rrp42-EL adopts the closed (open) conformation to 5 ±1 % (95 ±1 %) and that the open to closed transition takes place at a rate (k_ex_=k_open→closed_+k_closed→open_) of 5800 ±1400 s^-1^ (**Fig. 4G**). In the Exo10 complex the population of the closed conformation is significantly higher (26 ±1 %), whereas the exchange rate is reduced to 35 ±35 s^-1^ (**Fig. 4I**). The order parameter (S^2^) of the open conformation (∼0.1) is significantly lower than of the closed conformation (∼0.7), revealing that the loop in the open state is highly flexible, whereas it is stably fixed to the rest of the exosome complex in the closed state. In the presence of RNA, the open conformation in the Exo9 and Exo10 complex is occupied to 100% (**Fig. 4C, F, H, J**), indicating that Rrp42-EL is fully displaced by substrate RNA.

### Structural insights into Rrp42-EL

The Rrp42 extended loop is invisible in the static cryo-EM and crystal structures (**Fig. 1**). To obtain further structural and dynamic insights of Rrp42-EL in the closed and open conformation, we exploited molecular dynamics (MD) simulations of the Exo9 complex in aqueous solution. To initiate the MD simulations of the open state all missing loops of the here obtained cryo-EM structure were modeled using the ColabFold implementation of AlphaFold2 (*13*). The closed state was obtained by interactively modeling Rrp42-EL into the unoccupied cavity near Rrp41, a location that agrees with our NMR data (**Fig. 3**, **4**). Our MD simulations reflect that the open and the closed conformations are energetically stable (**Fig 5A**, **fig. S21**). The nanosecond timescale mobility of Rrp42-EL in the closed state is clearly reduced compared to the open state, as monitored by the C^α^ root mean square deviation (RMSD) of Rrp42-EL (**Fig. 5B**). This reduced flexibility within the MD simulations is fully consistent with the NMR order parameter (S^2^) analysis (**Fig. 4**, **fig. S18)**. Moreover, in the open state Rrp42-EL is remote to Rrp41 forming very few inter-subunit contacts restricted solely to Mtr3 (**Fig. 5C**). In contrast, in the closed state Rrp42-EL forms numerous contacts with Mtr3, Rrp43 and Rrp45 (**Fig. 5C**), in agreement with the PRE experiments (**Fig. 3**). In the closed conformation, Rrp42-A106 remains in close distance to Rrp41-D113, whereas the distance is much longer in the open state (**fig. S22, SI movie 1**), which agrees qualitatively with the ^19^F PRE data (**Fig. 4**, **fig. S18**). Interestingly, AlphaFold-predicted α-helical elements (residues K82-A94 and A106-N112) of Rrp42-EL begin to partially unfold during the simulation, indicating flexibility within the secondary structure (**fig. S23**). A comparison of the representative MD simulation structures complemented with Rrp44 from *S. cerevisiae* with the human Exo10 complex suggest that Rrp44 impacts the transition of Rrp42-EL between the open and closed conformation (**fig. S24**). The observed impact of Rrp44 agrees with what is expected from exosome structures of other organisms (**fig. S17**) and with our NMR experiments (**Fig. 4**, **fig. S16**). Finally, an analysis of the RNA channel within the representative MD simulation structures illustrates that the closed conformation of Rrp42-EL blocks the exit site of the RNA channel in the exosome core, whereas the RNA exit channel is unobstructed in the open state (**fig. S25**). Also in that regard, the MD and NMR data are fully consistent and explain that the closed state of Rrp42-EL is not observed in the presence of an RNA substrate (**Fig. 4**, **fig. S18**).

**Figure 5:**
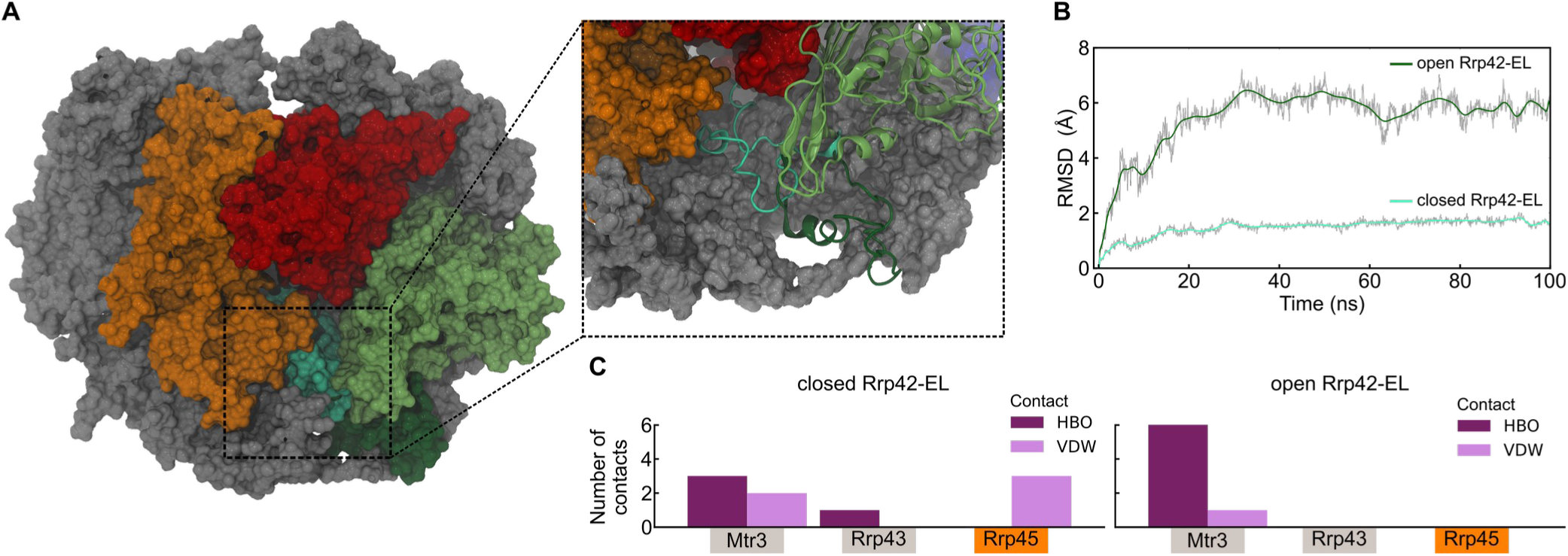
MD simulations of Rrp42-EL closed and open state. (**A**) Overlay of the representative structures from MD simulations of the completed Exo9 complex comparing the closed and open state of Rrp42-EL. The enlargement illustrates the distinct open (dark green) and closed (cyan) conformation of Rrp42-EL. (**B**) Structural dynamics of Rrp42-EL in the closed (dark green) and open (cyan) state revealed by the RMSD of the C^α^ carbons of Rrp42-EL within the MD simulation. The RMSD is smoothed by a Bézier curve. (**C**) Interaction network analysis between the Exo9 subunits and Rrp42-EL in the closed (left) and open (right) conformation. Shown are the number of hydrogen bonds (HBO, violet) and van-der-Waals contacts (VDW, pink) between Rrp42-EL and the indicated subunits.

### Rrp42-EL blocks an aberrant RNA path

The observation that Rrp42-EL can dynamically interact with the channel exit of the exosome raises the question of whether this is functionally relevant. To address this, we studied the activity of the Exo10 complex in the presence of full-length Rrp42 and with a version, in which Rrp42-EL was deleted (Rrp42^Δ93-125^). The RNA degradation rate in Exo10 is unaffected by the truncation of Rrp42 (**fig. S26**), which means that the loop displacement by substrate RNA comes at a low energetic cost. In the canonical substrate route, RNA threads through the Exo9 channel; it has, however, been shown that RNA can employ alternative paths to the Rrp44 active site that bypass the Exo9 channel (*32*, *53*). To assess if RNA can access Rrp44 via a direct path, we introduced an extension into a channel-lining loop of Rrp45, termed Rrp45-L, that has previously been shown to block the exosome channel in *S. cerevisiae* (*16*, *53*) (**fig. S27**). We observe that Rrp45-L reduces the activity of the exosome considerably, to ∼4% of wild-type activity (**fig. S26D**), confirming that the “through-channel” path is the major route that the RNA substrate employs. Next, we combined channel-blocked Exo10 Rrp45-L with the Rrp42^Δ93-125^ mutant, in which Rrp42-EL is deleted. Interestingly, we find that the activity in this exosome complex is partially recovered (**Fig. 6A**, **fig. S26**). Based on that we conclude that Rrp42-EL functions as a barrier that blocks an aberrant direct access path to the Rrp44 active site (**Fig. 6B**). Rrp42 in *C. thermophilum* thus contains a previously unidentified flexible “one-way-plug” that readily allows for passage of substrate RNA via the through-channel route but that prevents access to the Rrp44 active site via an aberrant direct path. The latter would result in the potentially detrimental ability of the exosome to degrade substrates that are not selected for processing or degradation by accessory factors, which interact with the cap subunits of the exosome complex.

**Figure 6:**
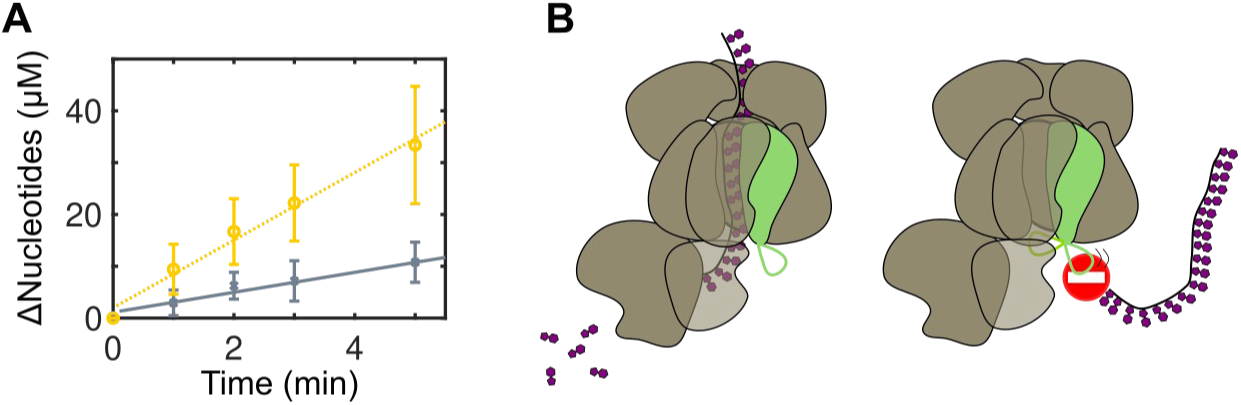
Rrp42-EL blocks an aberrant RNA access path. RNA activity assays for channel-blocked Exo10 Rrp45-L (gray crosses, solid line) and Exo10 Rrp45-L Rrp42^Δ93-125^ (yellow circles, dashed line). The lines are global linear fits to the linear activity regime. (**B**) Rrp42-EL allows on-path RNA to access Rrp44 for degradation (left) but blocks a direct access path towards Rrp44 (right).

### Conclusions

The “resolution revolution” in cryo-electron microscopy (*54*) and the remarkable performance of structure prediction algorithms have substantially increased insights into the relationship between protein structure and function. However, those methods provide static structural snapshots and often lack information on loop regions, protein dynamics and transient interactions, all of which may be crucial for protein function. Here, we demonstrate that dedicated NMR methods, such as CEST, EXSY, relaxation dispersion and PRE experiments and a combination of methyl-TROSY and ^19^F NMR, together with molecular dynamic simulations can complement static structural information, even in large, fully asymmetric eukaryotic molecular machines. In particular, we show that quantitative and functionally important information can be obtained on regions that are “invisible” in structures derived from cryo-EM and X-ray crystallography. Since large asymmetric or transiently formed complexes play a key role in virtually all aspects of molecular biology, we envision that the strategies to study large complexes by NMR and MD we laid out here will be of future importance to gain a deeper understanding of how protein structure, interactions and dynamics relate to function. We are convinced that the approach described here will spark further studies that facilitate the transition from 3D to 4D structural biology.

## Acknowledgments

We thank Iris Holdermann (MPI Tübingen) for support in the crystallization of the Exo9 complex, Janina Petters for help with the cloning, Nadine Stefan and Johanna Stöfl for excellent technical assistance, Jan Overbeck and Philip Wurm for support in conducting NMR experiments and David Stelzig for assistance with the RNA activity assays. All present and past group members are acknowledged for critically discussing the results in the course of the project.

## Funding

This project is funded from the European Union’s Horizon 2020 research and innovation programme under the Marie Skłodowska-Curie grant agreement No. 89550 (to JL), by the German Research Foundation (Deutsche Forschungsgemeinschaft) under grant agreement No. SP 1324/3-1 and by European Research Council under the European Union’s Seventh Framework Programme (FP7/2007– 2013), ERC grant agreement No. 616052 (to RS).

## Authors contributions

Conceptualization: RS, JL, DL, TR

Methodology: RS, JL, DL, TR

Investigation: JL, DL, AB, MP, TF

Formal Analysis: RS, JL, DL, TR, TF

Visualization: JL, DL, TF

Funding acquisition: RS, JL, TR

Project administration: RS

Supervision: RS, TR

Writing – original draft: RS, JL, DL, TR, TF with consent of all the authors

Writing – review & editing: RS, JL, DL, TR, TF with consent of all the authors

## Competing interests

The authors declare no competing interests.

## Data and materials availability

All data are available in the manuscript or the supplementary materials. Atomic coordinates have been deposited in the Protein Data Bank (PDB) with accession code 8PEL (crystal structure) and 8R1O (cryo-EM structure). Cryo-EM maps have been deposited in the Electron Microscopy Data Bank (EMDB) with accession code EMD-18825. Raw and processed data will be made available upon reasonable request.

## Supplementary Materials

Materials and Methods

Figs. S1 to S27

Tables S1 to S9

Movie S1

## Notes

### Competing Interest Statement

The authors have declared no competing interest.

### Summary of Updates

Molecular dynamics data were added to substantiate key observations.

