## Supplementary Material for "4D structural biology: quantitative dynamics in the eukaryotic RNA exosome complex"

### Title:

### Table of contents

Materials and methods Page S2

Supplementary Tables S1-S9 Page S17

Supplementary Figures S1-S27 Page S37

Supplementary References Page S67

### Materials and Methods

#### Molecular cloning

Codon-optimized constructs of the ten subunits of the *Chaetomium thermophilum* exosome (see table S5) were obtained from GenScript, cloned into pETM-11 vectors and expressed with an N-terminal hexahistidine-tag, a tobacco edge virus (TEV) cleavage site and a kanamycin resistance cassette. For monomeric Rrp43 and Rrp42$^{\Delta 93\text{-}125}$, expression yields were low but yields could be improved by co-expression with Rrp46 or Mtr3, respectively, from bicistronic constructs, in which the downstream gene (*rrp46* or *mtr3*) did not code for a hexahistidine tag or a TEV cleavage site. Tight binding of Rrp43 to Rrp46 and of Rrp42$^{\Delta 93\text{-}125}$ to Mtr3 allowed for co-purification of the subunits. In order to reduce the number of purifications required, this approach could also be used for wtMtr3-Rrp42 and wtRrp41-Rrp45, where appropriate, even though the monomeric proteins provided sufficient yields.

For cryo-EM and X-ray crystallography, Exo9 was expressed from a polycistronic pETM-11 plasmid that contained genes coding for all 9 subunits in the order: *rrp40-csl4-rrp4-rrp46-rrp43-mtr3-rrp42-rrp41-rrp45*. Only Rrp40 was expressed with an N-terminal hexahistidine-tag and a TEV cleavage site.

Point mutations, inserts and deletions of the original constructs were obtained using site-directed mutagenesis. Primers are listed in tables S6A (assignment mutants) and S6B (other constructs). All constructs were sequenced to confirm that mutations were correctly incorporated and are listed in table S6C (assignment constructs) and S6D (other constructs).

#### Protein expression

Plasmid was transformed into BL21(DE3) CodonPlus-RIL cells (StrataGene) and grown overnight at 37°C in an LB pre-culture containing 50 µg/ml kanamycin and 34 µg/ml chloramphenicol. The content of the growth media depended on the labeling scheme, where adequate antibiotics were added in all cases:

1a. For methyl NMR-based experiments, non-labeled (NMR inactive) constructs were expressed in $D_2O$ based minimal medium, which contained ~95% $D_2O$ and ~5% $H_2O$ (referred to as r$D_2O$ M9 medium), supplemented with 0.5 g/L $^{14}NH_4Cl$ and 4 g/L $^{1}H^{12}C$ glucose.

1b. Ile-$\delta1[^{13}CH_3]$ and Met-$\varepsilon1[^{13}CH_3]$ labeled ('IM-labeled') proteins were expressed in 99.8% $D_2O$ based minimal medium (referred to as $D_2O$ M9), supplemented with 0.5 g/L $^{14}NH_4Cl$ and 2 g/L $^2H^{12}C$ glucose, except for monomer assignment mutants, which were expressed in $rD_2O$ M9 medium.

2a. For Csl4 backbone assignments, Csl4 was expressed in $H_2O$ based minimal medium supplemented with 0.5 g/L $^{15}NH_4Cl$ and 2 g/L $^1H^{13}C$ glucose.
2b. For Csl4 sidechain assignments, Csl4 was expressed in $D_2O$ based minimal medium supplemented with 0.5 g/L $^{15}NH_4Cl$ and 2 g/L $^2H^{13}C$ glucose.

3. Constructs used for cryo-EM, X-ray crystallography and $^{19}F$ NMR experiments were expressed in LB medium.

Cells were inoculated from the pre-culture into $H_2O$ M9 (1. and 2.) or LB (3.) medium to an $OD_{600}$ of 0.1 and grown at 37°C to an $OD_{600}$ of 0.6 – 0.8. For non-deuterated and LB media (2a. and 3.), cells were then induced with 1 mM isopropyl ß-D-1-thiogalactopyranoside (IPTG). For deuterated media (1. and 2b.), cells were spun down (15 min, 1,300 xg) and resuspended into $rD_2O$ M9 (1a.) or $D_2O$ M9 (1b. and 2b.) medium, inoculated into another pre-culture to an $OD_{600}$ of 0.1 and grown overnight at 37°C. On the next day, cells were diluted into fresh $rD_2O$ M9 (1a.) or $D_2O$ M9 medium (1b. and 2b.) to an $OD_{600}$ of ~0.2 and grown to an $OD_{600}$ of 0.6 – 0.8. At this point cells were either induced with 1 mM IPTG (1a.) or, if constructs were to be IM-labeled, 60 mg/L $^2H^{12}C$ (1b.) or $^2H^{13}C$ (2b.) ketobutyric acid-4-$^{13}CH_3$ and 100 mg/L L-methionine-(*methyl*-$^{13}C$) (1b. and 2b.) were added and the cells were further incubated for 1 hour at 37°C prior to induction with 1 mM IPTG. Proteins were over-expressed for ~18 hours at 20°C, harvested by centrifugation (20 min, 6,000 xg), after which cell pellets were stored at -20°C until purification.

**Incorporation of tfmF using amber codon suppression**

The non-natural amino acid 4-trifluoromethyl-L-phenylalanine (tfmF) was incorporated into Rrp41 using amber codon suppression. The *rrp41* gene including an N-terminal hexahistidine-tag and a TEV cleavage site was cloned into an ampicillin resistance pBAD vector by Gibson assembly (*1*). An amber stop codon (TAG) was introduced at position G71, Q86 or D113 of Rrp41 using site-directed mutagenesis with primers as given in table S6B. The resulting plasmid was co-transformed with a tetracycline resistance pDule plasmid encoding for the tfmF amino-acyl-tRNA synthetase (tfmF-A65V-S158A) and its cognate suppressor $tRNA_{CUA}$ (*2*) into Top10 cells (Thermo Fisher Scientific). The pDule-tfmF-A65V-S158A vector was a kind gift from Ryan Mehl (Addgene plasmid #85484; http://n2t.net/addgene:85484; RRID: Addgene_85484). An LB preculture containing 100 µg/ml ampicillin and 15 µg/ml tetracycline was grown overnight at 37°C. Cells were inoculated into LB containing appropriate antibiotics to an $OD_{600}$ of 0.1 and grown to an $OD_{600}$ of 0.4 at 37°C at which point 1 mM tfmF was added to the solution. Cells were further grown for 1 hour at 37°C, shifted to 20°C and induced with 1% L-arabinose. Proteins were over-expressed for ~18 hours at 20°C, harvested by centrifugation (20 min, 6,000 xg) after which the cell pellets were stored at -20°C until purification.

**Protein purification**

Cell pellets were resuspended in 50 mM sodium phosphate buffer, pH 7.4, 150 mM NaCl, 0.5 mM DTT, 10 mM imidazole (buffer A), except for $Rrp44^{D168N, D536N}$ where 500 mM NaCl was used. Next, 0.1% (v/v) triton X-100 and 1 mg/L lysozyme were added. Cells were subsequently lysed by sonication, cell debris was removed by centrifugation (30 min, 39,000 xg) and the filtered lysate (1.2 µm) was passed onto a gravity flow column filled with 4 ml Ni-NTA resin. The column was washed with 5 to 10 column volumes of buffer A and additionally washed with buffer A supplemented with 5 M NaCl for $Rrp44^{D168N, D536N}$ to remove RNA. Protein was eluted from the resin using buffer A supplemented with 300 mM imidazole. 0.5 mg TEV protease were added to the eluate that was dialysed overnight into 20 mM HEPES, pH 7.5, 150 mM NaCl, 1 mM DTT (buffer B). All TEV-cleaved constructs contained Gly-Ala residues N-terminal to the protein sequence. The dialysate was subsequently passed onto a second Ni-column to remove the purification tag and TEV protease. The concentrated flow-through was subjected to a size-exclusion chromatography (SEC) purification step using a HiLoad 16/600 Superdex 200 pg (for full-length Rrp44) or a HiLoad 16/600 Superdex 75 pg (all other constructs) column in 10 mM HEPES buffer pH 7.5, 200 mM NaCl, 1 mM DTT (buffer C). For protein constructs that were subsequently linked to a TEMPO spin-label or reconstituted with a TEMPO spin-labeled construct (see below) a buffer devoid of DTT was used. Purity was assessed by sodium dodecyl sulfate polyacrylamide gel electrophoresis (SDS-PAGE). Concentrations were determined by $OD_{280}$ measurements and extinction coefficients that were computed with ProtParam (*3*). Purity of $Rrp44^{D168N, D536N}$ was further assessed by the $OD_{260}/OD_{280}$ absorption ratio, where only samples with an $OD_{260}/OD_{280}$ ratio below 0.75 were used.

#### $^{19}$F-labeling with BTFA

wtRrp42, Rrp42$^{C59S}$ and Rrp42$^{C59S,\ A106C}$ were labeled with 3-Bromo-1,1,1-trifluoroacetone (BTFA) prior to SEC purification. To that end, 1 mM DTT was added to the concentrated protein (ca. ~200 µM, 2 ml) and the solution was incubated for 15 min at room temperature under gentle agitation. Subsequently, 20 mM BTFA was added after which the linking reaction proceeded for 1 hour at room temperature under gentle agitation. Excess label was removed in the final SEC purification step.

#### TEMPO-labeling

Rrp42$^{C59S,\ A106C}$ and Csl4$^{C122S,\ E130C}$ were labeled with 4-maleimido-2,2,6,6-tetramethylpiperidine-1-oxyl (4-maleimido-TEMPO) prior to SEC purification. 4-maleimido-TEMPO was added in 3x excess to the DTT-free sample and incubated for 1 hour at room temperature under gentle agitation. Excess label was removed in the final SEC purification step using DTT-free buffer C. No DTT was added to the sample when TEMPO spin-labeled protein was reconstituted into Exo9 or Exo10 (see below).

#### Exosome reconstitution

To reconstitute Exo9 or Exo10, the individually purified subunits (or heterodimers) were mixed in stoichiometric amounts and incubated for 30 min at room temperature under gentle agitation. Next, the reconstituted exosome complex was purified by SEC with a HiLoad 16/600 Superdex 200 pg into buffer C (DTT-free if TEMPO spin-labeled protein was present). Successful reconstitution was assessed by SDS-PAGE (fig. S3).

#### X-ray crystallography

Crystals of Exo9 (8 mg/ml) grew in 0.2 M ammonium sulphate, 0.1 M sodium acetate pH 5.5, 10% PEG MME 2000, at 4°C, by vapor diffusion. Diffraction data were collected at the PXII beamline of the Swiss Light Source (SLS, Villigen, Switzerland). The crystals diffracted to 3.8 Å resolution, belonged to space group $P2_12_12_1$ with cell dimensions of a = 100.525 Å, b = 148.383 Å, c = 195.065 Å, α = β = γ = 90°, and contained one molecule in the ASU. Data were processed and scaled with XDS (*4*). The structure was solved by molecular replacement, using an *in silico* model (generated from the structures of the *S. cerevisiae* and *H. sapiens* homologs using MODELLER and AlphaFold) as search model (*5*, *6*). Iterative cycles of model building and refinement were carried out in COOT, PHENIX and ISOLDE (*7–9*). Data collection and refinement statistics are given in table S1A. Atomic coordinates have been deposited in the Protein Data Bank (PDB) with accession code 8PEL.

**Single particle cryo-EM**

To increase the purity of the complex and to exchange to cryo-EM buffer, the Exo9 complex was subjected to a second SEC purification step using a Superdex 200 10/300 GL column in 20 mM sodium phosphate buffer pH 7.5, 100 mM NaCl (buffer D). The protein was diluted to a final concentration of ~2 µM. Quantifoil R1.2/1.3 Cu300 holey carbon grids were glow discharged twice for 100 s, at 15 mA and 0.39 mBar in an easyGlow system (PELCO). 3 µl of sample were applied to freshly glow discharged grids using a Vitrobot mark IV plunge freezer (ThermoFisher Scientific). After a 5 s incubation at 4°C and 100% humidity, samples were blotted for 5 s using blot force 12 and plunged into liquid ethane. 6579 micrograph movies were collected on a CryoArm200 cryo-electron microscope (JEOL) equipped with a K2 direct electron detector (Gatan), in-column energy filter operated with slit width of 20 eV, and cold-field emission gun (low-flash interval 4 hours). Data were recorded using SerialEM (*10*), in a 5 x 5 multi-hole pattern, in counting mode, with a total dose of 40 $e^-/Å^2$ fractionated over 40 frames, and defocus range from -0.6 to -2 µm.

The data processing pipeline is depicted in fig. S2. Data were processed using RELION 4.0 (*11*). Particles were picked using the Topaz wrapper (*12*) within RELION and subjected to multiple rounds of 2D and 3D classification to eliminate partially disassembled complexes. The final 3D reconstruction was obtained from 276,958 particles and refined to an overall resolution of 3.19 Å (FSC cut-off 0.143). Iterative cycles of model building and refinement, using the refined crystal structure as starting model, were carried out in COOT, PHENIX and ISOLDE (*7*–*9*). Data collection and refinement statistics are given in table S1B. Atomic coordinates and density maps have been deposited in the PDB and in the Electron Microscopy Data Bank (EMDB) with accession codes 8R1O and EMD-1882, respectively.

**RNA *in vitro* transcription and purification**

The sequences of the RNAs used in this study are listed in table S7. RNA was obtained by *in vitro* transcription using an in-house purified T7 polymerase containing a P266L mutation (*13*). 1 µM template DNA and 1 µM T7 promoter oligonucleotide were mixed with 4 mM nucleotides, 30 mM $MgCl_2$, 10% (v/v) DMSO, 50 mM Tris pH 8.0, 0.01 % (v/v) triton X-100, 1 mM spermidine, 5 mM DTT and 18 µg/ml T7 polymerase and incubated for 4 hours at 37°C. 50 mM EDTA at pH 8.0 was subsequently added to dissolve phosphates and the RNA was precipitated by adding 300 mM sodium acetate at pH 5.0 and 70 vol% isopropanol followed by incubation for at least 1 hour at -20°C.

RNA was purified by anion exchange chromatography using a preparative DNAPac 100 column (Dionex) at 60°C with a linear buffer gradient from 20 mM Tris pH 8.0, 5 M urea (buffer E) to 20 mM Tris pH 8.0, 5 M urea, 1 M NaCl (buffer F). Fractions containing the desired RNA were pooled and RNA was precipitated by adding 300 mM sodium acetate at pH 5.0 and 70 vol% isopropanol and incubation for at least 1 hour at -20°C. Next, the RNA was pelleted, the supernatant was discarded and the pellet was washed with ethanol, dried at 37°C for 3 hours and re-suspended in $H_2O$. The purity of the preparation was assessed by Urea-PAGE on a 16% acrylamide gel.

**NMR sample preparation**

Methyl-TROSY NMR experiments were performed in $D_2O$-based buffer C. The final sample, in $H_2O$-based buffer C, was first concentrated to 200 µl, diluted 75 times with $D_2O$-based buffer C and then concentrated again. For all other experiments, 10% of $D_2O$-based buffer C was added to the sample for frequency locking. 46mer RNA (see table S7) was added in 1.5 times excess to either Exo9 or Exo10 Rrp44$^{D168N,\ D536N}$, which is a mutant that inactivates both endo- and exonucleolytic activity of Rrp44. Protein concentrations varied depending on the construct: for exosome samples, concentrations were typically between 50 – 120 µM, while monomers or dimers could often be investigated at higher concentrations. For NMR experiments on the exosome complex, 200 µl sample was placed into an NMR tube with a diameter of 3 mm. For all other constructs 500 µl sample was placed into an NMR tube with a diameter of 5 mm. Constructs used in this study and experiments conducted on them are listed in table S6E.

**NMR spectrometers**

NMR experiments were conducted on Bruker 500, 600 and 800 MHz Avance Neo spectrometers (11.7 T, 14.1 T and 18.8 T magnetic field strength, respectively) equipped with triple resonance cryogenic TCI probeheads cooled with liquid helium (800 MHz) or liquid nitrogen (500 and 600 MHz). For the 500 and 600 MHz spectrometers, the $^{1}H$ channel was tuned to $^{19}F$ frequency (471 MHz and 565 MHz, respectively) for fluorine NMR experiments.

**Methyl NMR and backbone assignment experiments**

2D methyl-TROSY spectra were collected using the SOFAST-HMQC pulse sequence (*14*) with carbon acquisition times of 30 ms (Exo9 and larger) or 60 ms (exosome monomers) and an interscan delay of 0.5 s at 40°C.

To assign Csl4 Ile-δ1 resonances, standard backbone assignment experiments (HNCACB, HNCA, HNCOCACB, HNCO, HNCACO) were conducted for the monomer and the Ile-δ1 methyl groups were assigned by standard H(CCCO)NH and C(CCO)NH experiments, in which either the Ile-δ1 $^{1}H$ or $^{13}C$ chemical shift is correlated with $^{1}H^{N}$ and $^{15}N$ chemical shifts of the preceding residue. In addition, assignments were also obtained from Met and Ile point mutations (table S6C).

**$^{19}$F NMR experiments**

Experiments were acquired with an acquisitions time of 0.05 s, 1 – 1.5 s interscan delay at 25°C. Chemical exchange saturation transfer experiments (CEST) were conducted at $B_1$ field strengths as indicated in the figures (10 – 25 Hz) applied for $t_{CEST}$ = 400 ms with 67 frequency offsets ranging from -2450 Hz to +2450 Hz. The central frequency (0 Hz) was set on resonance with the most intense fluorine resonance. CEST intensities were referenced to the intensity determined for offsets at ±10,000 Hz. EXSY experiments were acquired using 12 mixing delays $t_{ZZ}$ (1, 2, 5, 10, 25, 50, 75, 100, 150, 200, 400, 600 ms) and an acquisition time in the indirect dimension of 14 ms. $R_1$ relaxation rates were determined employing an inversion recovery pulse sequence with at least 8 delays $t_{R1}$ ranging between 0.001 s and 8 s. $R_1$ relaxation rates were obtained by fitting:

$$I(t_{R1}) = I_{\infty}\left(1 - 2\exp(-R_1 t_{R1})\right) \quad \text{(Eq. S1)}$$

to experimental intensities $I(t_{R1})$ for varying delay times $t_{R1}$.

CEST and inversion recovery experiments were conducted on a 500 MHz spectrometer, EXSY experiments were conducted on a 600 MHz spectrometer.

Constant time Carr-Purcell-Meiboom-Gill (CPMG) relaxation dispersion (RD) experiments were conducted with a relaxation delay ($T_{CPMG}$) between 2 and 20 ms (see table S8) using at least 20 frequencies ($\nu_{CPMG}$) if $T_{CPMG} \geq 6$ ms and 10 frequencies if $T_{CPMG} < 6$ ms. The maximum frequency that was used was 5000 Hz and the minimum frequency depended on the length of $T_{CPMG}$ (see table S8). CPMG RD experiments were conducted on a 500 MHz spectrometer unless indicated otherwise.

**PRE experiments**

For paramagnetic relaxation enhancement (PRE) experiments, an initial methyl-TROSY or $^{19}F$ spectrum was acquired in the presence of non-reduced TEMPO spin-label providing intensities $I_{para}$. Then, 5 mM sodium ascorbate was added to reduce the spin-label and another spectrum was acquired to obtain intensities $I_{dia}$. Methyl PREs were calculated as

$$\Gamma_{^{13}CH_3} = \frac{I_{para}}{I_{dia}} \quad \text{(Eq. S2)}$$

Additionally, for PRE experiments of tfmF-labeled samples, an initial $T_1$ inversion recovery experiment was acquired providing $R_{1,para}$ (Eq. S1). After addition of 5 mM sodium ascorbate another $T_1$ inversion recovery experiment was acquired to obtain $R_{1,dia}$ (Eq. S1). The $R_1$-based PRE, $\Gamma_1$, was calculated as

$$\Gamma_1 = R_{1,para} - R_{1,dia} \quad \text{(Eq. S3)}$$

**Data analysis**

NMR data were processed using the NMRPipe/NMRDraw software suite (*15*). Methyl resonance intensities were obtained with NMRPipe while $^{19}F$ resonance integrals were obtained using an in-house Matlab script. Assignments of Csl4 were performed in Cara (*16*).

For model fitting, in-house Matlab scripts were employed. In all fitting routines, the target function

$$\chi^2 = \sum_{\exp} \sum_{i=1} \left( \frac{O_{\mathrm{exp,i}} - O_{\mathrm{calc,i}}}{\sigma_{\mathrm{exp,i}}} \right)^2 \quad \text{(Eq. S4)}$$

was minimized using the fminsearch routine in Matlab. In Eq. S4 $O_{exp,i}$ corresponds to an experimentally determined data point (*i*) in one of the recorded datasets *exp* {$^{19}F$ RD data of Exo9 in the absence of RNA at 471 and 565 MHz; CEST data in the absence of RNA at $B_1$ fields of 10 (Exo9 and Exo10), 15 (Exo10) and 25 Hz (Exo9 and Exo10); EXSY data of Exo10 in the absence of RNA, a 1D NMR spectrum of Exo10 in the absence of RNA; 1D NMR spectra Exo9 and Exo10 in the presence and absence of RNA and in the paramagnetic and diamagnetic states (to assess $\Gamma_2$); intensities in $R_1$ inversion recovery experiments for Exo9 and Exo10 in the presence and absence of RNA and in the paramagnetic and diamagnetic states (to assess $\Gamma_1$)}. $O_{calc,i}$ corresponds to a back-calculated value of an observable based on the model parameters, as described below. $\sigma_{exp,i}$ is an estimate of the measurement uncertainty for a data-point and is based on the noise level in the spectra or duplicate measurements.

A two-site exchange process between a ground state G (the open conformation) and an excited state

E (the closed conformation) as described by the equilibrium $G \leftrightarrows E$, with $k_{ex}=k_{\mathrm{EG}}+k_{\mathrm{GE}}$, $p_E=\frac{k_{\mathrm{GE}}}{k_{ex}}$, $p_G=\frac{k_{\mathrm{EG}}}{k_{ex}}$, and $p_E=1-p_G$ was fitted to the data.

To reduce the number of fitting parameters and thus over-fitting of the data, we assumed that Rrp42$^{\mathrm{C59S,\ A106C\text{-}TFA}}$ chemical shifts of the ground ($\omega_G$) and excited ($\omega_E$) states were the same for Exo9 and Exo10 complexes in the presence and absence of RNA. Furthermore, we assumed that the RNA bound complexes were 100% in the open conformation as demonstrated by PRE experiments.

The **CPMG relaxation dispersion** data was back-calculated numerically using the equations derived by Baldwin (*17*). These equations provide an analytical solution of a system undergoing two-site exchange and are not limited to a specific timescale of the motion. $R_{2,\mathrm{inf}}$ of the ground and excited states were assumed to be the same.

To fit the **CEST** data, the signal intensities (*I*) at offsets $\omega_{\mathrm{CEST}}$ were back-calculated according to:

$$I(\omega_{\mathrm{CEST}})=\vec{I}_{proj}*\exp(\boldsymbol{M}\,t_{\mathrm{CEST}})\vec{I}_0 \quad \text{(Eq. S5)}$$

where $t_{\mathrm{CEST}}$ is the time during which the weak $B_1$ field is applied. The equilibrium *z* magnetization is $\vec{I}_0=\left(E/2 \quad I_x^G \quad I_y^G \quad I_z^G \quad I_x^E \quad I_y^E \quad I_z^E\right)^T=\left(1/2 \quad 0 \quad 0 \quad p_G \quad 0 \quad 0 \quad p_E\right)^T$, where $E$ is the identity operator and $I_{\{x,y,z\}}^{\{G,E\}}$ are the components of the magnetization vector in *x*, *y*, or *z* direction for the G and E states, respectively. $\boldsymbol{M}$ is the evolution matrix (according to the Bloch-McConnell equations) (*18*):

$$\boldsymbol{M}=\begin{pmatrix} 0 & 0 & 0 & 0 & 0 & 0 & 0 \\ 0 & -R_2^G-k_{\mathrm{GE}} & -\omega_G & \omega_1 & k_{\mathrm{EG}} & 0 & 0 \\ 0 & \omega_G & -R_2^G-k_{\mathrm{GE}} & 0 & 0 & k_{\mathrm{EG}} & 0 \\ 2R_1^G p_G & -\omega_1 & 0 & -R_1^G-k_{\mathrm{GE}} & 0 & 0 & k_{\mathrm{EG}} \\ 0 & k_{\mathrm{GE}} & 0 & 0 & -R_2^E-k_{\mathrm{EG}} & -\omega_E & \omega_1 \\ 0 & 0 & k_{\mathrm{GE}} & 0 & \omega_E & -R_2^E-k_{\mathrm{EG}} & 0 \\ 2R_1^E p_E & 0 & 0 & k_{\mathrm{GE}} & -\omega_1 & 0 & -R_1^E-k_{\mathrm{EG}} \end{pmatrix}$$

where, $R_1^G$, $R_1^E$, $R_2^G$ and $R_2^E$ are the longitudinal and transverse relaxation rates of state G and E, respectively. $\omega_G=\omega_{\mathrm{CEST}}-\omega_G$ and $\omega_E=\omega_{\mathrm{CEST}}-\omega_E$ (in rad/s) denote the offsets between the CEST frequency and the chemical shifts of the ground and excited states, respectively. $\omega_1=\gamma_{\mathrm{F}}B_1$ is the weak $B_1$ field (in rad/s) applied from the y-direction and $\gamma_{\mathrm{F}}$ is the gyromagnetic ratio of fluorine. $\hat{I}_{proj}=\begin{pmatrix}0 & 0 & 0 & 1 & 0 & 0 & 1\end{pmatrix}$ is the vector that projects the magnetization onto $I_z^G+I_z^E$ , which results in observable magnetization.

To fit the **EXSY** data, the experimental signal intensities were back-calculated according to (*19*):

$$\begin{pmatrix} I_{\mathrm{GG}} & I_{\mathrm{EG}} \\ I_{\mathrm{GE}} & I_{\mathrm{EE}} \end{pmatrix} = S_{EXSY} * \exp\left(\begin{pmatrix} -k_{\mathrm{GE}}-R_1^G & k_{\mathrm{EG}} \\ k_{\mathrm{GE}} & -k_{\mathrm{EG}}-R_1^E \end{pmatrix} t_{ZZ}\right)\begin{pmatrix} p_G & 0 \\ 0 & p_E \end{pmatrix} \quad \text{(Eq. S6)}$$

where $S_{\mathrm{EXSY}}$ is a scaling factor of the experimental intensities, *I* are the intensities of either the auto peaks of the ground ($I_{GG}$) or excited ($I_{EE}$) state, or the intensities of the cross peaks between the ground and excited states ($I_{EG}$ or $I_{GE}$) and $t_{\mathrm{ZZ}}$ is the EXSY mixing time.

The **1D NMR** spectra were simulated based on

$$I(\omega)=S_{spectum} * \left|\Re\left(\Sigma\left(\boldsymbol{M}^{-1} * \vec{I}_0\right)\right)\right| \quad \text{(Eq. S7)}$$

where $S_{\mathrm{spectrum}}$ is a scaling factor for the intensity in a specific spectrum, $\omega$ is the offset, $\vec{I}_0=\begin{pmatrix} p_G & p_E \end{pmatrix}^T$ and

$$\boldsymbol{M}=\begin{pmatrix} -R_2^G+i(\omega_G-\omega)-k_{\mathrm{GE}} & k_{EG} \\ k_{\mathrm{GE}} & -R_2^E+i(\omega_E-\omega)-k_{\mathrm{EG}} \end{pmatrix}$$ in the absence of PREs or

$$\boldsymbol{M}=\begin{pmatrix} -R_2^G-\Gamma_2^G+i(\omega_G-\omega)-k_{\mathrm{GE}} & k_{EG} \\ k_{\mathrm{GE}} & -R_2^E-\Gamma_2^E+i(\omega_E-\omega)-k_{\mathrm{EG}} \end{pmatrix}$$ in the presence of PREs,

where $\Gamma_2^{\{G,E\}}$ are the $R_2$ based PRE effects in states G or E.

To fit the **$R_1$ PRE** rates, the experimental signal intensities (*I*) from inversion recovery experiments were back-calculated according to:

$$I(t_{relax})=\vec{I}_{proj} * \exp(\boldsymbol{M}\, t_{\mathrm{R1}})\,\vec{I}_0 \quad \text{(Eq. S8)}$$

where $t_{\mathrm{R1}}$ is the relaxation delay and $\vec{I}_0=\begin{pmatrix} 1/2 & p_G & p_E \end{pmatrix}^T$ . The evolution matrix $\boldsymbol{M}$ is

$$\boldsymbol{M}=\begin{pmatrix} 0 & 0 & 0 \\ 2\left(R_1^G+\Gamma_1^G\right)p_G & -R_1^G-\Gamma_1^G-k_{\mathrm{GE}} & k_{\mathrm{EG}} \\ 2\left(R_1^E+\Gamma_1^E\right)p_E & k_{\mathrm{GE}} & -R_1^E-\Gamma_1^E-k_{\mathrm{EG}} \end{pmatrix}$$

where the $\Gamma_1^{\{G,E\}}$ are the $R_1$ based PRE effects in states G or E and $\vec{I}_{proj}=\begin{pmatrix}0 & 1 & 1\end{pmatrix}$ is the projection vector that results in observable magnetization.

The determined $\Gamma_1^G$ , $\Gamma_1^E$ , $\Gamma_2^G$ and $\Gamma_2^E$ values were used in combination with the Solomon-

Bloembergen equations to extract order parameters $S^2$ of the ground and excited state according to (*20–22*)

$$\Gamma_1 = \frac{1}{r^6}\frac{2}{5}\left(\frac{\mu_0}{4\pi}\right)^2 \gamma_I^2 g^2 \mu_B^2 s(s+1) J(\omega_I)$$

$$\Gamma_2 = \frac{1}{r^6}\frac{1}{15}\left(\frac{\mu_0}{4\pi}\right)^2 \gamma_I^2 g^2 \mu_B^2 s(s+1)\left(4J(0)+3J(\omega_I)\right) \quad \text{(Eq. S9)}$$

where the spectral density $J$ is expressed as $J(\omega)=\frac{S^2\tau_c}{1+(\omega\tau_C)^2}+\frac{(1-S^2)\tau_t}{1+(\omega\tau_t)^2}$ , $\tau_c^{-1}=\tau_R^{-1}+\tau_s^{-1}$ and $\tau_t^{-1}=\tau_r^{-1}+\tau_s^{-1}+\tau_i^{-1}$ . In these equations $r$ is the distance between the spin-label and the probing nucleus (in the ground or excited state), $\mu_0$ is the permeability of vacuum ($1.257 \cdot 10^{-6}$ N $A^{-2}$), $\gamma_I$ is the gyromagnetic ratio of flourine ($251.815 \cdot 10^{6}$ rad $T^{-1}$ $s^{-1}$), $g$ is the Landé factor (-2.002), $\mu_B$ is the magnetic moment of the free electron ($-9.285 \cdot 10^{-24}$ J $T^{-1}$), $s$ is the electron spin quantum number (0.5), $\omega_I$ is the Larmor frequency of a flourine nucleus (471 MHz), $S^2$ is the squared order parameter of the ground or excited state and $\tau_i$ is the correlation time of the vector connecting the spin-label and the probing nucleus in the ground or excited state, $\tau_R$ is the rotational correlation time of the protein complex (assumed to be 100 ns for Exo9 and 140 ns for Exo10) and $\tau_s$ is the electron relaxation time (assumed to be 100 ns). The lower sensitivity of $\Gamma_2$ as compared to $\Gamma_1$ for fast internal motions follows from Eq. S9, in which the term *J(0)* dominates the value of $\Gamma_2$. *J(0)* in turn is dominated by motions of the entire complex ($\tau_R$) as long as $S^2$ is not too small. On the other hand, $\Gamma_1$ depends only on $J(\omega_I)$, which is sensitive to fast internal motions ($\tau_i$).

Uncertainties in the fitted model parameter (which are globally: $p_E$ in Exo9 and Exo10, $k_{ex}$ in Exo9 and Exo10; for Rrp42$^{C59S,\ A106C\text{-}TFA}$: the chemical shifts of the ground and excited states, $R_{2,inf}$ at 471 and 565 MHz, $R_1$ and $R_2$ in Exo9 and Exo10, scaling factors for EXSY spectra and for PRE spectra in Exo9 and Exo10 in the absence and presence of RNA; for Rrp41$^{D113tmF}$: the chemicals shifts of the ground and exited states in Exo9 and Exo10, $R_1$ and $R_2$ in Exo9 and Exo10 in the absence and presence of RNA, $\Gamma_1$ and $\Gamma_2$ for Exo9 and Exo10 for the ground and excited states and scaling factors for the PRE $R_1$ inversion recovery experiments; for the Solomon-Bloembergen equations: $S^2$, $\tau_i$ and $r$ for the ground and excited states) were obtained from Monte-Carlo simulations where 200 artificial datasets were created based on the measurement uncertainties. Subsequently, the model was fitted to these datasets. In that procedure the starting parameters for the fit were varied randomly by 5% to prevent model bias. Model parameters are reported as best fit value +/- standard deviations. The distributions of the fitting parameters, that are not necessarily Gaussian, are displayed in fig. S19.

**Chemical shift perturbations**

Chemical shift perturbations (CSPs) in $^{1}H$ ($\Delta\delta_H$) and $^{13}C$ ($\Delta\delta_C$) dimensions were combined to a global CSP ($\Delta\delta$) by:

$$\Delta\delta=\sqrt{\left(\left(\frac{\Delta\delta_C}{4}\right)^2+\Delta\delta_H^2\right)} \quad \text{(Eq. S10)}$$

**Molecular Modeling**

To generate a complete model of the Exo9 complex, we predicted the missing loops of the here obtained cryo-EM structure using AlphaFold2-based (*6*) ab intio structure prediction. Employing a local implementation of ColabFold 1.5.3 (*23*) we modeled five structures of the complete Exo9 complex, each of which was refined in 12 iterative refinement cycles. The models were ranked based on the Local Distance Difference Test (lDDT) (*24*) and the Template Modeling score (TM) (*25*). Subsequently the models were optimized with the original united atom AMBER force field (*26*). To complete the experimentally unresolved missing loops, we locally aligned the predicted structure to the cryo-EM structure and added the missing loops from the predicted structure to the experimental structure.

The resulting completed structure represents the open state of the Rrp42-EL (residues 77 to 117) assigned based on the NMR results (Fig. 4, fig. S18) as the distance between Asp113 of Rrp41 and Ala106 of Rrp42 is with 51.3 Å larger compared to the experimentally expected distance of 7 Å for the closed state (Table S4). To obtain a structure of the closed state Rrp42-EL was manually modeled into the unoccupied cavity near Rrp41 to achieve a short spin label distance comparable to the one observed in the $^{19}F$ PRE experiments (Fig. 4, fig. S18). During the manual modeling we iteratively repeated the process of interactively updating the coordinates of Rrp42-EL and subsequently energy optimized the loop using the MAXIMOBY program suit version 2023 (CHEOPS, Germany) with the original united atom AMBER force field (*26*). The resulting structure reflects the closed state with a distance of 14 Å between Asp113 of Rrp41 and Ala106 of Rrp42. The structures of both states were further refined to the cryo-EM density map with molecular dynamics flexible fitting (MDFF) (*27*, *28*). MDFF runs were set up with QwikMD (*29*) and performed with NAMD 2.13 (*30*) employing the CHARM36 force field (*31*). During an initial 800 step minimization phase, existing secondary structure elements (α-helix, β-sheet) as well as peptide isomerism (cis/trans) and center chirality were conserved. This was followed up by a 40 ns simulation phase at 300K in implicit solvent. The refined structures were energy optimized employing MAXIMOBY (CHEOPS, Germany).

**MD Simulations**

To adapt the structural models to aqueous environment and study the loop dynamics in solution at room temperature (293.15K) we ran MD simulations initiated by the two completed Exo9 models of the open and closed state obtained as described above. The structures were protonated based on the local pKa values of each residue calculated at a pH value of 7 following Nielsen and Vriend (*32*). Water molecules of the first and second solvation shell of the protein complex were set using a Vedani-like algorithm (*33*) implemented in MAXIMOBY. To prevent self-interactions of the protein due to periodic boundary conditions within the MD simulation, a cubic simulation box with the dimensions 17.97*17.97*17.97 nm was set and filled with water molecules, sodium and calcium ions, at physiological conditions using the solvation workflow implemented in GROMACS 2021 (*34*). Steric clashes between the hydrogen of the solvation shell and bulk water were locally resolved through energy optimization in MAXIMOBY (CHEOPS, Germany). Subsequently we performed MD simulations with GROMACS 2021 (*34*) utilizing the OPLS/all-atom force filed (*35*). First, the system was heated to room temperature (293.15K) in 1 ns with a step size of 1 fs within a nVT simulation, meaning the number of atoms (n), the volume (V), and the temperature (T) is constant while the pressure (p) is flexible. The temperature was kept constant using a V-rescale thermostat (*36*) with a coupling constant 0.1 ps. The heating was performed in two steps, over the first 100 ps the temperature was raised continuously from 0 K to 100 K, in the following 900 ps the temperature was raised continuously to 293.15 K. The heating procedure was followed by a 1 ns nVT simulation with a stepsize of 1 fs under the same conditions as for the heating but at 293.15 K. Next we performed a 10 ns npT run with a step size of 1 fs in which the number of atoms, pressure and temperature remained constant using a Berendsen barostat (*37*) (coupling constant 0.5 ps) and a V-rescale thermostat (*36*) (coupling constant 0.1 ps) while the volume was kept flexible. Following these equilibration steps we performed a 100 ns npT production run (step size 2 fs) for each of the two systems that was used for further analysis. In the production run, the temperature was controlled by a Nosé-Hoover thermostat (*38*, *39*) (coupling constant 0.5 ps) and a Parrinello-Rahman barostat (*40*) (coupling constant 2.5 ps) which perform better for equilibrated systems.

**Simulation Evaluation**

To evaluate the stability of the simulation we calculated the root mean square deviation (RMSD) of the $C^{\alpha}$ atoms of each snapshot of the simulation trajectory compared to the starting structure. Changes in the secondary structure were monitored using the defined secondary structure of proteins (DSSP) algorithm (*41*). Inter and intra protein subunit interactions were determined with the contact matrix algorithm in MAXIMOBY (CHEOPS, Germany) and the PyContact plugin (*42*) for VMD 1.9.4. VMD 1.9.4 and PyMOL 3.0 were used for visual inspection of protein structures and simulation trajectories.

To obtain a representative structures of the MD simulations, each 0.1 ns frame of the simulation was analyzed for the current conformation and contacts of each residue within the system, resulting in a contact matrix for every frame. Each contact and conformation were weighted based on their importance. A mean matrix across the second half of the simulation was calculated and scored against every frame of the simulation. The frame most closely resembling the weighted mean vector was defined as representative for the simulation.

The number of interactions in Fig. 5C was obtained by statistical analysis of the simulation based on the contact matrix algorithm implemented in the MAXIMOBY program package (CHEOPS, Gerrmany). Each contact with a percentage presence of over 40% across the second half of the simulation was deemed significant for the count.

**Activity assays**

*Urea-PAGE analysis*

5 µM 80mer RNA (see table S7) was mixed to 10 mM HEPES buffer pH 7.5, 200 mM NaCl and 5 mM $MgCl_2$ and incubated for 5 min at 40°C, after which a reference sample (0 min) in absence of the exosome was taken. Next, 1 µM exosome was rapidly mixed with the RNA-buffer solution and samples were taken at different time points (1, 2, 4, 8, 16, 32, 64, and 128 min) while the reaction proceeded at 40°C. For each sample, the reaction was stopped by rapidly mixing 2x Urea-PAGE loading dye containing 8 M Urea, 20 mM EDTA, 2 mM Tris-HCl pH 8 and 0.0001% (w/v) bromophenol blue in a 1:1 ratio. The activity was qualitatively assayed by Urea-PAGE on 16% acrylamide gels.

*HPLC analysis*

5 µM 80mer RNA (see table S7) was mixed to 10 mM HEPES buffer pH 7.5, 200 mM NaCl and

5 mM $MgCl_2$ and incubated for 5 min at 40°C, after which a reference sample (0 min) in absence of the exosome was taken. Next, 0.5 µM exosome was rapidly mixed with the RNA-buffer solution and samples were taken at different time points (1, 2, 3, 5, and 10 min) while the reaction proceeded at 40°C. For each sample, the reaction was stopped by rapidly mixing a three times volume excess of 8 M Urea and heating the sample to 95°C. Next, the nucleotide and RNA concentrations of each sample were determined by high-performance liquid chromatography (HPLC) using anion exchange on an analytical DNAPac PA100 column (Thermofisher) heated to 40°C. The samples were applied onto the column using buffer E supplemented with 100 mM NaCl and eluted using gradient steps of buffer F (described in table S9). Elution peaks of the nucleotides and the RNA were integrated and concentrations were obtained by comparing integrals to a calibration curve for GMP with known concentrations. The concentrations were scaled by the ratio of the extinction coefficient of the RNA (for the RNA elution peak) or an average extinction coefficient per nucleotide (for the nucleotide elution peak) divided by the extinction coefficient of GMP. To obtain the catalytic rate $k_{cat}$, a linear equation, for which the slope is $k_{cat}$, was simultaneously fitted to the linear regime (time points 0 – 3 min for wtExo10 and Exo10 Rrp42$^{\Delta 93-125}$, time points 0 – 10 min for Exo10 Rrp45-L and time points 0 – 5 min for Exo10 Rrp45-L Rrp42$^{\Delta 93-125}$) for the nucleotide and RNA data. The experiments were conducted for three distinct protein batches that were independently expressed and purified, with three technical repeats each.

**Sequence alignments**

The sequences of the subunits ctRrp41, ctRrp42 and ctRrp45 were aligned with homologous protein sequences from human, *S. cerevisiae* and the *S. solfataricus* using Clustal Omega (*43*).

**Table S1: X-ray crystallography and cryo-EM data collection and refinement statistics.** Statistics for the *C. thermophilum* core exosome (ctExo9) structure obtained by (**A**) X-ray crystallography (PDB ID: 8PEL) and (**B**) single particle cryo-EM (PDB ID: 8R1O, EMDB ID: EMD-18825). Statistics for the highest-resolution shell in the X-ray structure are shown in parentheses.

**A**

| **Parameter** | **Value** |
|---|---|
| **Data collection** | |
| Space group | P $2_12_12_1$ |
| Cell dimensions | |
| a, b, c (Å) | 100.525, 148.383, 195.065 |
| α = β = γ (°) | 90, 90, 90 |
| Wavelength (Å) | 0.9999 |
| Resolution range (Å) | 48.77 – 3.81 (3.946 – 3.81) |
| Total reflections | 393908 (37705) |
| Unique reflections | 29143 (2793) |
| Completeness (%) | 99.57 (97.14) |
| Redundancy | 13.5 (13.5) |
| I/σ(I) | 5.01 (1.19) |
| $CC_{1/2}$ | 0.984 (0.586) |
| $CC^*$ | 0.996 (0.86) |
| $R_{merge}$ | 0.5272 (2.127) |
| $R_{meas}$ | 0.5479 (2.211) |
| $R_{pim}$ | 0.1481 (0.5971) |
| **Refinement** | |
| Resolution range (Å) | 48.77 – 3.81 |
| No. reflections | 29114 (2790) |
| $R_{work}$ | 0.2427 (0.3486) |
| $R_{free}$ | 0.2923 (0.3830) |
| No. protein atoms (without H) | 17656 |
| Average B factor ($Å^2$) | 103.52 |

| | |
|---|---|
| R.m.s. deviations | |
| Bond lengths (Å) | 0.004 |
| Bond angles (°) | 0.690 |
| Ramachandran plot | |
| Favored (%) | 97.64 |
| Allowed (%) | 2.32 |
| Outliers (%) | 0.04 |
| Rotamer outliers (%) | 0.05 |
| Clashscore | 14 |
| MolProbity score | 1.29 |

**B**

| **Parameter** | **Value** |
|---|---|
| **Data collection and processing** | |
| Microscope | JEOL CryoArm |
| Voltage (kV) | 200 |
| Camera | Gatan K2 Summit |
| Magnification | 60000 |
| Pixel size (Å) | 0.772 |
| Total electron exposure ($e^-$/Å$^2$) | 40 |
| No. frames collected during exposure | 40 |
| Defocus range (µm) | 0.6 – 2.2 |
| Energy filter slit width (eV) | 20 |
| No. micrographs collected | 6579 |
| No. micrographs used | 6572 |
| No. extracted particles | 2448810 |
| **Reconstruction** | |
| No. refined particles | 1541277 |
| No. particles used in the final reconstruction | 276958 |
| Point-group | C1 |
| Resolution (FSC 0.143) (Å) | |
| Global (unmasked/masked) | 3.19/2.99 |

| | |
|---|---|
| Local | 2.907 – 3.707 |
| Map sharpening B factor (Å$^2$) | -114.7 |
| **Refinement** | |
| Resolution (Å) | 3.2 |
| No. protein atoms (without H) | 17918 |
| No. protein residues | 2335 |
| Average B factor (Å$^2$) | 125.57 |
| R.m.s. deviations | |
| Bond lengths (Å) | 0.006 |
| Bond angles (°) | 0.937 |
| Ramachandran plot | |
| Favored (%) | 97.60 |
| Allowed (%) | 2.40 |
| Outliers (%) | 0 |
| Rotamer outliers (%) | 0.56 |
| Clashscore | 4 |
| MolProbity score | 1.30 |
| $CC_{volume}$ | 0.830 |

**Table S2: Ile-δ1 and Met-ε1 assignments.** Ile-δ1 and Met-ε1 resonance assignments of (**A**) Csl4, (**B**) Rrp41 and (**C**) Rrp45 as monomer and when reconstituted in Exo9 and Exo10. Note that multiple resonances for one residue are arbitrarily assigned as 'A' or 'B', i. e. A or B assignments for different resonances do not necessarily report on the same conformation. n.d.: not determined/ not assigned (e.g. due to severe signal overlap or resonance broadening beyond detection).

**A**

| **Resonance** | **monomer** | | **in Exo9** | | **in Exo10** | |
|---|---|---|---|---|---|---|
| | **$^{1}$H (ppm)** | **$^{13}$C (ppm)** | **$^{1}$H (ppm)** | **$^{13}$C (ppm)** | **$^{1}$H (ppm)** | **$^{13}$C (ppm)** |
| Csl4-I18 | 0.981 | 11.734 | 1.023 | 12.117 | 1.022 | 12.111 |
| Csl4-I60[1] | 0.912 | 10.003 | 0.926 | 9.981 | 0.926 | 9.980 |
| Csl4-I72 | 0.038 | 11.255 | -0.002 | 10.656 | -0.002 | 10.661 |
| Csl4-I95[1] | 0.912 | 10.003 | 0.926 | 9.981 | 0.926 | 9.980 |
| Csl4-I102 | 0.828 | 6.792 | 0.810 | 7.330 | 0.812 | 7.326 |
| Csl4-I108 | 0.822 | 9.135 | 0.845 | 9.297 | 0.842 | 9.289 |
| Csl4-I110 | 1.012 | 11.000 | 1.171 | 11.837 | 1.162 | 11.830 |
| Csl4-I119 | 0.707 | 11.182 | 0.628 | 10.991 | 0.631 | 11.026 |
| Csl4-I135 | 0.236 | 10.833 | 0.275 | 11.010 | 0.273 | 10.984 |
| Csl4-I140 | 0.921 | 12.351 | 0.958 | 12.327 | 0.956 | 12.317 |
| Csl4-I159 | 0.765 | 6.465 | 0.903 | 6.590 | 0.898 | 6.634 |
| Csl4-I165 | 0.876 | 11.237 | 0.888 | 11.238 | 0.883 | 11.214 |
| Csl4-I185 | 1.003 | 12.883 | 1.078 | 12.729 | 1.078 | 12.730 |
| Csl4-I207 | 0.828 | 7.858 | 0.880 | 7.850 | 0.881 | 7.847 |
| Csl4-M1 | 2.161 | 14.535 | 2.203 | 14.454 | 2.206 | 14.452 |
| Csl4-M195 | 1.492 | 15.104 | n.d. | n.d. | n.d. | n.d. |

[1] Csl4-I60 and Csl4-I95 overlap.

**B**

| **Resonance** | | **monomer** | | **in Exo9** | | **in Exo10** | |
|---|---|---|---|---|---|---|---|
| | | **$^{1}$H (ppm)** | **$^{13}$C (ppm)** | **$^{1}$H (ppm)** | **$^{13}$C (ppm)** | **$^{1}$H (ppm)** | **$^{13}$C (ppm)** |
| Rrp41-I30 | A | 1.039 | 11.306 | 1.036 | 10.674 | 1.023 | 10.732 |
| | B | 1.022 | 11.116 | | | | |
| Rrp41-I96 | | 0.872 | 12.765 | n.d. | n.d. | n.d. | n.d. |

| | | | | | | | |
|---|---|---|---|---|---|---|---|
| Rrp41-I98 | | 0.765 | 9.270 | 0.676 | 9.295 | 0.484 | 9.404 |
| Rrp41-I116 | A | 0.855 | 8.743 | 0.858 | 8.650 | 0.695 | 7.570 |
| | B | 0.826 | 8.221 | | | | |
| Rrp41-I117[2] | | 0.947 | 9.522 | 0.961 | 9.749 | 0.939 | 9.712 |
| Rrp41-I141 | A | 0.750 | 11.258 | 0.748 | 11.628 | 0.734 | 11.591 |
| | B | 0.716 | 11.333 | | | | |
| Rrp41-I143 | | 0.560 | 11.137 | 0.554 | 11.288 | 0.549 | 11.286 |
| Rrp41-I159 | A | 0.540 | 10.599 | 0.576 | 10.438 | 0.567 | 10.425 |
| | B | 0.540 | 10.495 | | | | |
| Rrp41-I171 | A | 0.778 | 10.465 | 0.659 | 10.877 | 0.637 | 10.687 |
| | B | 0.690 | 10.345 | | | | |
| Rrp41-I251[2] | | 0.969 | 9.890 | 0.977 | 9.608 | 0.939 | 8.987 |
| Rrp41-I254 | | 0.948 | 11.027 | 0.942 | 11.202 | 0.899 | 11.099 |
| Rrp41-I267 | A | 0.897 | 9.825 | 0.793 | 11.005 | 0.782 | 11.073 |
| | B | 0.898 | 9.741 | | | | |
| Rrp41-M1 | | 2.209 | 14.537 | 2.206 | 14.578 | 2.194 | 14.524 |
| Rrp41-M44 | | 2.232 | 13.755 | n.d. | n.d. | n.d. | n.d. |
| Rrp41-M49 | | 1.949 | 14.518 | n.d. | n.d. | n.d. | n.d. |
| Rrp41-M119 | A | 2.122 | 14.784 | 2.119 | 14.578 | 2.116 | 14.571 |
| | B | 2.114 | 14.679 | | | | |
| Rrp41-M173 | | 1.991 | 14.258 | n.d. | n.d. | n.d. | n.d. |
| Rrp41-M240 | | 2.050 | 14.170 | n.d. | n.d. | n.d. | n.d. |
| Rrp41-M266 | A | 1.855 | 14.219 | n.d. | n.d. | n.d. | n.d. |
| | B | 1.832 | 14.162 | | | | |
| Rrp41-M281 | | 2.200 | 14.483 | n.d. | n.d. | n.d. | n.d. |

[2] Rrp41-I117 and Rrp41-I251 overlap for Exo9. Resonance positions were determined from point mutants.

**C**

| **Resonance** | | **monomer** | | **in Exo9** | | **in Exo10** | |
|---|---|---|---|---|---|---|---|
| | | **$^{1}$H (ppm)** | **$^{13}$C (ppm)** | **$^{1}$H (ppm)** | **$^{13}$C (ppm)** | **$^{1}$H (ppm)** | **$^{13}$C (ppm)** |
| Rrp45-I76 | | 0.558 | 11.335 | 0.565 | 11.366 | 0.557 | 11.467 |
| Rrp45-I79 | | 0.940 | 11.756 | 0.728 | 11.828 | 0.809 | 12.100 |
| Rrp45-I111 | A | 0.813 | 7.902 | 0.936 | 9.477 | 0.904 | 9.624 |
| | B | 0.805 | 7.705 | | | | |
| Rrp45-I133 | | 0.867 | 8.518 | 0.807 | 7.625 | 0.871 | 7.583 |

| | | | | | | | |
|---|---|---|---|---|---|---|---|
| Rrp45-I152 | | 0.302 | 10.677 | 0.240 | 10.640 | 0.239 | 10.626 |
| Rrp45-I168 | | 0.835 | 11.065 | n.d. | n.d. | n.d. | n.d. |
| Rrp45-I175 | | 0.790 | 10.966 | 0.809 | 11.557 | 0.734 | 13.242 |
| Rrp45-I206[3] | | 0.828 | 10.716 | 0.804 | 10.929 | 0.800 | 10.948 |
| Rrp45-I225 | A<br>B | 1.111 | 12.501 | 1.126 | 13.752 | 1.113 | 13.428 |
| Rrp45-I233 | | n.d. | n.d. | n.d. | n.d. | n.d. | n.d. |
| Rrp45-I236 | | n.d. | n.d. | n.d. | n.d. | n.d. | n.d. |
| Rrp45-I255 | | 1.182 | 10.954 | 1.120 | 10.983 | 1.122 | 10.933 |
| | A | 2.199 | 14.608 | | | | |
| Rrp45-M1 | B | 2.234 | 14.580 | n.d. | n.d. | n.d. | n.d. |
| | C | 2.171 | 14.766 | | | | |
| Rrp45-M86 | | 2.141 | 14.477 | n.d. | n.d. | n.d. | n.d. |
| Rrp45-M140 | | 2.063 | 14.258 | n.d. | n.d. | n.d. | n.d. |
| Rrp45-M227 | | 2.190 | 14.520 | n.d. | n.d. | n.d. | n.d. |

[3] Rrp45-I206 overlaps with an unassigned resonance in Exo9 and Exo10.

**Table S3: $\Gamma_1$ PRE experiments for the Rrp41 entry and exit loop.** (**A**) $R_1$ rates for paramagnetic (para) and diamagnetic (dia) Rrp41$^{G71tfmF}$ (entry loop) Csl4$^{C122S,\ E130C\text{-}TEMPO}$ in Exo9 and Exo9 with RNA as well as paramagnetic relaxation enhancements $\Gamma_1$ for these systems. (**B**) $R_1$ rates for paramagnetic (para) and diamagnetic (dia) Rrp41$^{D113tfmF}$ (exit loop) Rrp42$^{C59S,\ A106C\text{-}TEMPO}$ in Exo9, Exo9 with RNA, Exo10 and Exo10 with RNA as well as paramagnetic relaxation enhancements $\Gamma_1$ for these systems. These rates were obtained from Eq. S1.

**A**

| **System** | **$R_{1,para}$ ($s^{-1}$)** | **$R_{1,dia}$ ($s^{-1}$)** | **$\Gamma_1$ ($s^{-1}$)** |
|---|---|---|---|
| *Rrp41$^{G71tfmF}$ (Csl4$^{C122S,\ E130C\text{-}TEMPO}$)* | | | |
| Exo9 | 1.3±0.1 | 1.0±0.1 | 0.3±0.2 |
| Exo9 with RNA | 1.0±0.1 | 1.0±0.1 | 0.0±0.2 |

**B**

| **System** | **$R_{1,para}$ ($s^{-1}$)** | **$R_{1,dia}$ ($s^{-1}$)** | **$\Gamma_1$ ($s^{-1}$)** |
|---|---|---|---|
| *Rrp41$^{D113tfmF}$ (Rrp42$^{C59S,\ A106C\text{-}TEMPO}$)* | | | |
| Exo9 | 2.5±0.3 | 1.0±0.1 | 1.5±0.5 |
| Exo9 with RNA | 1.0±0.1 | 0.9±0.1 | 0.1±0.2 |
| Exo10 | 1.4±0.2 | 1.0±0.1 | 0.4±0.3 |
| Exo10 with RNA | 0.9±0.1 | 0.9±0.1 | 0.0±0.2 |

**Table S4: Rrp45-EL fit parameters.** Parameters obtained from fitting a two-state exchange model to dynamics and PRE data of Rrp42$^{C59S,\ A106C\text{-}TFA}$ in Exo9 and in Exo10.

| Parameter | Exo9 | Exo10 |
|---|---|---|
| *Global* | | |
| $k_{ex}$ ($s^{-1}$) | 5800±1400 | 35±35 |
| $p_{closed}$ | 0.05±0.01 | 0.26±0.01 |
| *$^{19}$F Rrp42$^{C59S,\ A106C\text{-}TFA}$* | | |
| $\omega_{open}$ (ppm) | -84.156 ± 0.002 | |
| $\omega_{closed}$ (ppm) | -83.795 ± 0.009 | |
| $R_1$ ($s^{-1}$) | 4.4±0.8 | 3.6±0.2 |
| $R_2$ ($s^{-1}$) | 100±16 | 234±9 |
| $R_{2,500}$ ($s^{-1}$) | 41.6±0.6 | n.d. |
| $R_{2,600}$ ($s^{-1}$) | 53.8±0.8 | n.d. |
| *$^{19}$F Rrp41$^{D113\text{-}tfmF}$ (Rrp42$^{C59S,\ A106C\text{-}TEMPO}$)* | | |
| $\Gamma_{1,open}$ ($s^{-1}$) | 0.54±0.20 | 0.00±0.00 |
| $\Gamma_{2,open}$ ($s^{-1}$) | 111±23 | 130±26 |
| $\Gamma_{1,closed}$ ($s^{-1}$) | 5.3±6.6 | 4.6±6.7 |
| $\Gamma_{2,closed}$ ( $s^{-1}$) | (16.1±0.6)*10$^3$ | (18.8±0.7)*10$^3$ |
| *Solomon-Bloembergen* | | |
| $r_{open}$ (Å) | 11.6±0.2 | |
| $S_{open}^2$ | 0.12±0.02 | |
| $\tau_{i,open}$ (ps) | 11±5 | |
| $r_{closed}$ (Å) | 6.83±0.08 | |
| $S_{closed}^2$ | 0.74±0.04 | |
| $\tau_{i,closed}$ (ns) | 24±59 | |

**Table S5: Uniprot accession codes for *Chaetomium thermophilum* proteins used in this study.**

| Protein | Accession code |
|---|---|
| Rrp45 | G0S755 |
| Rrp41 | G0SC21 |
| Rrp43 | G0S1P1 |
| Rrp46 | G0SCD1 |
| Rrp42 | G0RZG4 |
| Mtr3 | P0CT46 |
| Rrp40 | G0RZX8 |
| Rrp4 | G0S9A0 |
| Csl4 | G0SE33 |
| Rrp44 | G0SEX3 |

**Table S6: Primers and protein constructs used in this study.** (**A**) Primers for assignment mutants. (**B**) Primers for constructs other than assignment mutants. (**C**) Protein constructs used for assignments. Note, that assignments of Csl4 Ile-δ1 residues were additionally obtained from conventional assignment methods as described in materials and methods. (**D**) Internal references for plasmids. (**E**) Protein constructs and experiments conducted on them.

**A**

| Construct | forward primer | reverse primer |
|---|---|---|
| Csl4-I18L | GTCAACTGCTGGGTCCGCTGAGTAAATACCAACCGGGTC | GACCCGGTTGGTATTTACTCAGCGGACCCAGCAGTTGAC |
| Csl4-I60L | GTGAAACGTCTGAATCGCCTGACCCCGGCACCGACG | CGTCGGTGCCGGGGTCAGGCGATTCAGACGTTTCAC |
| Csl4-I95L | CGGTCGTAAACGCGAACTGCTGCCGGAAGTGGG | CCCACTTCCGGCAGCAGTTCGCGTTTACGACCG |
| Csl4-I102V | CTGCCGGAAGTGGGTAACGTCGTTCTGTGCCGTGTC | GACACGGCACAGAACGACGTTACCCACTTCCGGCAG |
| Csl4-I110L | CCGTGTCATCCGCCTGACCCCGCGTCAGG | CCTGACGCGGGGTCAGGCGGATGACACGG |
| Csl4-I140V | CTGATTCGTGTTCAAGATGTTCGCGCCACCGAAAAAGACC | GGTCTTTTTCGGTGGCGCGAACATCTTGAACACGAATCAG |
| Csl4-I159L | CGTCCGGGCGATCTGGTGCGCGCCGAAG | CTTCGGCGCGCACCAGATCGCCCGGACG |
| Csl4-I185L | CAATGAACTGGGCGTTCTGCTGGCGACCAGTGAAG | CTTCACTGGTCGCCAGCAGAACGCCCAGTTCATTG |
| Csl4-I207L | GAATACCGCGATCCGCTGACCGGCCTGACGGAAC | GTTCCGTCAGGCCGGTCAGCGGATCGCGGTATTC |
| Csl4-M1A | CTTTATTTTCAGGGCGCCGCGACGACGACGCAACCGAC | GTCGGTTGCGTCGTCGTCGCGGCGCCCTGAAAATAAAG |
| Csl4-M195A | CAGTGAAGCCGGTAATACGCTGTATCCGGTGTCATGGC | GCCATGACACCGGATACAGCGTATTACCGGCTTCACTG |
| Rrp41-I30V | GTCGCGTTCATGCCCAAGTTCGCACCCAGG | CCTGGGTGCGAACTTGGGCATGAACGCGAC |
| Rrp41-I96V | CCGAAGTCGTGGTTTCCGTTGTGATCGCAGG | CCTGCGATCACAACGGAAACCACGACTTCGG |
| Rrp41-I98V | CGTGGTTTCCATTGTGGTCGCAGGTT | CGGAACTAAAACCTGCGACCACAATGG |

| | | |
|---|---|---|
| | TTAGTTCCG | AAACCACG |
| Rrp41-I116V | CACGGCCGTAACGATAAACGCGTTATCGAAATGCAAAGC | GCTTTGCATTTCGATAACGCGTTTATCGTTACGGCCGTG |
| Rrp41-I117V | CCGTAACGATAAACGCATTGTCGAAATGCAAAGCACCG | CGGTGCTTTGCATTTCGACAATGCGTTTATCGTTACGG |
| Rrp41-I141V | GTTCCCGCATTCACAGGTTACGATCTCGCTGC | GCAGCGAGATCGTAACCTGTGAATGCGGGAAC |
| Rrp41-I143V | GCATTCACAGATTACGGTCTCGCTGCACGTCC | GGACGTGCAGCGAGACCGTAATCTGTGAATGC |
| Rrp41-I159V | CTGCTGGCTGCGCTGGTTAATGCGGCAACCCTG | CAGGGTTGCCGCATTAACCAGCGCAGCCAGCAG |
| Rrp41-I171V | GCTTGTGTTGATGCCGGTGTCCCGATGACCGATTATGTC | GACATAATCGGTCATCGGGACACCGGCATCAACACAAGC |
| Rrp41-I251V | GTGGACGGCTGTAAACAGGTTCGTGCCATCCTG | CAGGATGGCACGAACCTGTTTACAGCCGTCCAC |
| Rrp41-I254L | GTAAACAGATTCGTGCCCTGCTGGATCACGTTGTCC | GGACAACGTGATCCAGCAGGGCACGAATCTGTTTAC |
| Rrp41-I267V | GGTCGTCGCATGGTCCGTGAGGGTGCGGTTG | CAACCGCACCCTCACGGACCATGCGACGACC |
| Rrp41-M1A | CTTTATTTTCAGGGCGCCGCGCCGCTGGACACGAG | CTCGTGTCCAGCGGCGCGGCGCCCTGAAAATAAAG |
| Rrp41-M44A | GTAGCTCTTATCTGGAAGCGGGCCATACCAAAGTGATG | CATCACTTTGGTATGGCCCGCTTCCAGATAAGAGCTAC |
| Rrp41-M49A | GAAATGGGCCATACCAAAGTGGCGTGCGTTGTTACCGGTC | GACCGGTAACAACGCACGCCACTTTGGTATGGCCCATTTC |
| Rrp41-M119A | CGATAAACGCATTATCGAAGCGCAAAGCACCGTTGCC | GGCAACGGTGCTTTGCGCTTCGATAATGCGTTTATCG |
| Rrp41-M173A | GTTGATGCCGGTATCCCGGCGACCGATTATGTCGTGG | CCACGACATAATCGGTCGCCGGGATACCGGCATCAAC |
| Rrp41-M240A | GTTTCTCGCCTGGAAGGCGCGCTGGCGGTCGGTGTG | CACACCGACCGCCAGCGCGCCTTCCAGGCGAGAAAC |
| Rrp41-M266A | CCAAAAAGGTCGTCGCGCGATCCGTGAGGGTGC | GCACCCTCACGGATCGCGCGACGACCTTTTTGG |
| Rrp41-M281A | CGTCAGCCTGGATGACGCGGATGAAG | CAATCTTCATCCGCGTCATCCAGGCTG |

| | | |
|---|---|---|
| | ATTG | ACG |
| Rrp45-I76V | CGTCCGCTGGACGGTGTTTTTACCATCGCAAC | GTTGCGATGGTAAAAACACCGTCCAGCGGACG |
| Rrp45-I79V | CTGGACGGTATTTTTACCGTGGCAACGGAACTGAGTCCG | CGGACTCAGTTCCGTTGCCACGGTAAAAATACCGTCCAG |
| Rrp45-I111V | CTGCTGGAAAAAACCGTTCGTCGCAGTGGCGCTC | GAGCGCCACTGCGACGAACGGTTTTTTCCAGCAG |
| Rrp45-I133V | GGTCAGAAATGTTGGTCAGTTCGCGTTGACGTCCATGTG | CACATGGACGTCAACGCGAACTGACCAACATTTCTGACC |
| Rrp45-I152V | GACGCGGCCTGCGTGGCAGTGGTTGCAG | CTGCAACCACTGCCACGCAGGCCGCGTC |
| Rrp45-I168V | CGCAAACCGGATACCAGCGTTGAATCTGGTGTTCTGACC | GGTCAGAACACCAGATTCAACGCTGGTATCCGGTTTGCG |
| Rrp45-I175V | GAATCTGGTGTTCTGACCGTGTATACGCCGGCCGAAC | GTTCGGCCGGCGTATACACGGTCAGAACACCAGATTC |
| Rrp45-I206V | GGCGATGAAGGTGAAGTTGCTGTGCTGGACGCG | CGCGTCCAGCACAGCAACTTCACCTTCATCGCC |
| Rrp45-I225V | CGCGTCGGCTCATGCACGGTGTCGATGAACAAACATG | CATGTTTGTTCATCGACACCGTGCATGAGCCGACGCG |
| Rrp45-I233V | GATGAACAAACATGGTGAAGTTTGTCAGATTGCAAAACTGGG | CCCAGTTTTGCAATCTGACAAACTTCACCATGTTTGTTCATC |
| Rrp45-I236V | CATGGTGAAATTTGTCAGGTTGCAAAACTGGGCGGCACC | GGTGCCGCCCAGTTTTGCAACCTGACAAATTTCACCATG |
| Rrp45-I255L | GCTGCAATGCACCTCTCTGGCTCTGACGAAAG | CTTTCGTCAGAGCCAGAGAGGTGCATTGCAGC |
| Rrp45-M1A | CTTTATTTTCAGGGCGCCGCGCCGCGTGAAGTG | CACTTCACGCGGCGCGGCGCCCTGAAAATAAAG |
| Rrp45-M86A | CGGAACTGAGTCCGGCGACCAGCCCGACGTTC | GAACGTCGGGCTGGTCGCCGGACTCAGTTCCG |
| Rrp45-M140L | CGTTGACGTCCATGTGCTGTCGCACGATGGCAATCTG | CAGATTGCCATCGTGCGACAGCACATGGACGTCAACG |
| Rrp45-M227A | GGCTCATGCACGATCTCGGCGAACAAACATGGTG | CACCATGTTTGTTCGCCGAGATCGTGCATGAGCC |

**B**

| Construct | forward primer | reverse primer |
|---|---|---|
| Csl4-C122S | GTTACGATTCTGGTCTCTGGCGATACCGTGCTG | CAGCACGGTATCGCCAGAGACCAGAATCGTAAC |
| Csl4-E130C | CGTGCTGGACGCGTGCTGGCAGGGTCTGATTC | GAATCAGACCCTGCCAGCACGCGTCCAGCACG |
| Rrp41-G71amber | GAACCGGGTGCAGGCACCACGTAGGGTGGCGGTGCTG | CAGCACCGCCACCCTACGTGGTGCCTGCACCCGGTTC |
| Rrp41-Q86amber | GTGGCGGTAGTGGCGGTTAGGGTAAAGAAGCCGAAG | CTTCGGCTTCTTTACCCTAACCGCCACTACCGCCAC |
| Rrp41-D113amber | GTAAACGCCACGGCCGTAACTAGAAACGCATTATCGAAATG | CATTTCGATAATGCGTTTCTAGTTACGGCCGTGGCGTTTAC |
| Rrp42-C59S | GTTCTGCACGCGTGTCCTTCGCCGATGGCAC | GTGCCATCGGCGAAGGACACGCGTGCAGAAC |
| Rrp42-A106C | GACGAAGAAGGCTATTGCAAAGTCGGTGCGGATAAC | GTTATCCGCACCGACTTTGCAATAGCCTTCTTCGTC |
| Rrp42-Δ93-125 | GTTGCGAGCGCCGGCGTCCGCGATG | CATCGCGGACGCCGGCGCTCGCAAC |
| Rrp45-94ELGESEGESEG95 (Rrp45-L) | CGAACTGGGCGAATCTGAAGGCGAATCTGAAGGCCGTCCGACCGAAACGGAAGTTC | GAACTTCCGTTTCGGTCGGACGGCCTTCAGATTCGCCTTCAGATTCGCCCAGTTCG |
| Rrp44-PIN | GCATTTCGTTCGGGCGTTATTCTTTAATCGTTTCTTC | GCATTTCGTTCGGGCGTTATTCTTTAATCGTTTCTTC |
| Rrp44-PIN+CS | CGTGCTGGATTGCCTGCCGAAAACGGGCTAACACGACTGGCGCGTGCCGGAAG | CTTCCGGCACGCGCCAGTCGTGTTAGCCCGTTTTCGGCAGGCAATCCAGCACG |
| Rrp44-D168N | CTGTGAACGATCGTAATAACCGCGCCGTTCG | CGAACGGCGCGGTTATTACGATCGTTCACAG |
| Rrp44-D536N | GGTTGTCAGGACATTAATGATGCACTGCACAGTCG | CGACTGTGCAGTGCATCATTAATGTCCTGACAACC |

**C**

| Construct | Internal database reference | Methyl-TROSY HMQC acquired | | |
|---|---|---|---|---|
| | | monomer | Exo9 | Exo10 |
| Csl4-I60L | #1972 | yes | no | no |
| Csl4-I95L | #1970 | yes | no | no |
| Csl4-I102V | #2160 | no | yes | no |
| Csl4-I110L | #2139 | yes | yes | no |
| Csl4-I159L | #2141 | yes | yes | no |
| Csl4-I185L | #2140 | yes | no | no |
| Csl4-I207L | #2138 | yes | no | no |
| Csl4-M1A | #1969 | yes | no | no |
| Csl4-M195A | #1971 | yes | no | no |
| Rrp41-I30V | #2399 | yes | no | no |
| Rrp41-I96V | #2385 | yes | no | no |
| Rrp41-I98V | #2380 | yes | no | no |
| Rrp41-I116V | #2381 | yes | yes | no |
| Rrp41-I117V | #2387 | yes | yes | yes |
| Rrp41-I141V | #2401 | yes | yes | yes |
| Rrp41-I143V | #2402 | yes | yes | no |
| Rrp41-I159V | #2403 | yes | yes | yes |
| Rrp41-I171V | #2404 | yes | yes | no |
| Rrp41-I251V | #2400 | yes | yes | yes |
| Rrp41-I254L | #2382 | yes | no | no |
| Rrp41-I267V | #2424 | yes | yes | no |
| Rrp41-M1A | #2386 | yes | no | no |
| Rrp41-M44A | #2379 | yes | no | no |
| Rrp41-M49A | #2405 | yes | no | no |
| Rrp41-M119A | #2384 | yes | no | no |
| Rrp41-M173A | #2406 | yes | no | no |
| Rrp41-M240A | #2407 | yes | no | no |
| Rrp41-M266A | #2408 | yes | no | no |
| Rrp41-M281A | #2409 | yes | no | no |
| Rrp45-I76V | #2195 | yes | no | no |
| Rrp45-I79V | #2196 | yes | yes | yes |

| | | | | |
|---|---|---|---|---|
| Rrp45-I111V | #2197 | yes | yes | yes |
| Rrp45-I133V | #2191 | yes | yes | yes |
| Rrp45-I152V | #2192 | yes | no | no |
| Rrp45-I168V | #2193 | yes | yes | no |
| Rrp45-I175V | #2194 | yes | yes | yes |
| Rrp45-I206V | #2202 | yes | yes | no |
| Rrp45-I225V | #2203 | yes | no | no |
| Rrp45-I233V | #2198 | yes | no | no |
| Rrp45-I236V | #2204 | yes | no | no |
| Rrp45-I255L | #2205 | yes | no | no |
| Rrp45-M1A | #2199 | yes | no | no |
| Rrp45-M86A | #2200 | yes | no | no |
| Rrp45-M140L | #2253 | yes | no | no |
| Rrp45-M227A | #2201 | yes | no | no |

**D**

| **Construct** | **Internal database reference** |
|---|---|
| Csl4 | #1247 |
| Rrp4 | #1233 |
| Rrp40 | #1213 |
| Rrp41 | #1245 |
| Rrp42 | #1214 |
| Rrp44 | #1242 |
| Rrp45 | #1215 |
| Mtr3 | #1246 |
| Rrp41-Rrp45 | #1351 |
| Mtr3-Rrp42 | #1355 |
| Rrp43-Rrp46 | #1353 |
| Csl4$^{C122S, E130C}$ | #2741 |
| Rrp42$^{C59S}$ | #2172 |
| Rrp42$^{C59S, A106C}$ | #2176 |
| Rrp44$^{D168N, D536N}$ | #1671 |
| Rrp44$^{PIN}$ | #1835 |
| Rrp44$^{PIN+CS}$ | #2099 |

| | |
|---|---|
| Mtr3-Rrp42$^{\Delta 93\text{-}125}$ | #2445 |
| Rrp45-94ELGESEGESEG95 (Rrp45-L) | #2539 |
| Rrp41$^{G71amber}$ | #2716 |
| Rrp41$^{Q86amber}$ | #2661 |
| Rrp41$^{D113amber}$ | #2664 |

**E**

| **Construct** | **Type(s) of experiment** |
|---|---|
| Exo9 | Cryo-EM, X-ray crystallography |
| U-$^{15}$N/$^{13}$C Csl4 | HNCACB, HNCA, HNCOCACB, HNCO, HNCACO, H(CCCO)NH, C(CCO)NH |
| Monomeric IM-labeled Csl4 | Methyl TROSY HMQC |
| Monomeric IM-labeled Rrp41 | Methyl TROSY HMQC |
| Monomeric IM-labeled Rrp45 | Methyl TROSY HMQC |
| Exo9, IM-labeled Csl4 | Methyl TROSY HMQC |
| Exo9, IM-labeled Rrp41 | Methyl TROSY HMQC |
| Exo9, IM-labeled Rrp45 | Methyl TROSY HMQC |
| Exo10, IM-labeled Csl4 | Methyl TROSY HMQC |
| Exo10, IM-labeled Rrp41 | Methyl TROSY HMQC |
| Exo10, IM-labeled Rrp45 | Methyl TROSY HMQC |
| Exo9, 46mer RNA, IM-labeled Csl4 | Methyl TROSY HMQC |
| Exo9, 46mer RNA, IM-labeled Rrp41 | Methyl TROSY HMQC |
| Exo9, 46mer RNA, IM-labeled Rrp45 | Methyl TROSY HMQC |
| Exo10, Rrp44$^{D168N,\ D536N}$, 46mer RNA, IM-labeled Csl4 | Methyl TROSY HMQC |
| Exo10, Rrp44$^{D168N,\ D536N}$, 46mer RNA, IM-labeled Rrp41 | Methyl TROSY HMQC |
| Exo10, Rrp44$^{D168N,\ D536N}$, 46mer RNA, IM-labeled Rrp45 | Methyl TROSY HMQC |
| Exo9, Rrp42$^{C59S,\ A106C\text{-}TEMPO}$, IM-labeled Csl4 | Methyl PRE |
| Exo9, Rrp42$^{C59S,\ A106C\text{-}TEMPO}$, IM-labeled Rrp41 | Methyl PRE |
| Exo10, Rrp42$^{C59S,\ A106C\text{-}TEMPO}$, IM-labeled Rrp41 | Methyl PRE |
| Exo9, Rrp42$^{C59S,\ A106C\text{-}TEMPO}$, IM-labeled Rrp45 | Methyl PRE |
| Exo10, Rrp42$^{C59S,\ A106C\text{-}TEMPO}$, IM-labeled Rrp45 | Methyl PRE |
| Exo9, Rrp42$^{C59S,\ A106C\text{-}TEMPO}$, 46mer RNA, IM-labeled Rrp45 | Methyl PRE |
| Exo10, Rrp44$^{D168N,\ D536N}$, Rrp42$^{C59S,\ A106C\text{-}TEMPO}$, 46mer RNA, IM-labeled Rrp45 | Methyl PRE |

| | |
|---|---|
| Exo9, Rrp41$^{Q86tfmF}$ | $^{19}$F 1D, CPMG RD |
| Exo10, Rrp41$^{Q86tfmF}$ | $^{19}$F 1D, CPMG RD |
| Exo10, Rrp44$^{D168N, D536N}$, 46mer RNA, Rrp41$^{Q86tfmF}$ | $^{19}$F 1D |
| Exo9, Csl4$^{C122S, E130C-TEMPO}$, Rrp41$^{G71tfmF}$ | $^{19}$F $\Gamma_1$ PRE, $^{19}$F $\Gamma_2$ PRE |
| Exo9, Csl4$^{C122S, E130C-TEMPO}$, 46mer RNA, Rrp41$^{G71tfmF}$ | $^{19}$F $\Gamma_1$ PRE, $^{19}$F $\Gamma_2$ PRE |
| Exo9, Rrp41$^{D113tfmF}$ | $^{19}$F 1D, $^{19}$F CPMG RD |
| Exo10, Rrp41$^{D113tfmF}$ | $^{19}$F 1D, $^{19}$F CPMG RD |
| Exo10, Rrp44$^{D168N, D536N}$, 46mer RNA, Rrp41$^{D113tfmF}$ | $^{19}$F 1D |
| Exo9, Rrp42$^{C59S, A106C-TEMPO}$, Rrp41$^{D113tfmF}$ | $^{19}$F $\Gamma_1$ PRE, $^{19}$F $\Gamma_2$ PRE |
| Exo9, Rrp42$^{C59S, A106C-TEMPO}$, 46mer RNA, Rrp41$^{D113tfmF}$ | $^{19}$F $\Gamma_1$ PRE, $^{19}$F $\Gamma_2$ PRE |
| Exo10, Rrp42$^{C59S, A106C-TEMPO}$,Rrp41$^{D113tfmF}$ | $^{19}$F $\Gamma_1$ PRE, $^{19}$F $\Gamma_2$ PRE |
| Exo10, Rrp44$^{D168N, D536N}$, Rrp42$^{C59S, A106C-TEMPO}$, 46mer RNA, Rrp41$^{D113tfmF}$ | $^{19}$F $\Gamma_1$ PRE, $^{19}$F $\Gamma_2$ PRE |
| wtRrp42$^{TFA}$ | $^{19}$F 1D |
| Rrp42$^{C59S, TFA}$ | $^{19}$F 1D |
| Rrp42$^{C59S, A106C-TFA}$ | $^{19}$F 1D |
| Mtr3, Rrp42$^{C59S, A106C-TFA}$ | $^{19}$F 1D |
| Exo9, Rrp42$^{C59S, A106C-TFA}$ | $^{19}$F 1D, $^{19}$F CEST, $^{19}$F CPMG RD |
| Exo9, 46mer RNA, Rrp42$^{C59S, A106C-TFA}$ | $^{19}$F 1D, $^{19}$F CEST, $^{19}$F CPMG RD |
| Exo10, Rrp42$^{C59S, A106C-TFA}$ | $^{19}$F 1D, $^{19}$F CEST, $^{19}$F/$^{19}$F EXSY |
| Exo10, Rrp44$^{D168N, D536N}$, 46mer RNA, Rrp42$^{C59S, A106C-TFA}$ | $^{19}$F 1D, $^{19}$F CEST, $^{19}$F CPMG RD |
| Exo10, 80mer RNA, Rrp42$^{C59S, A106C-TFA}$ | activity assay |
| Exo10, Rrp44$^{PIN}$, Rrp42$^{C59S, A106C-TFA}$ | $^{19}$F 1D, $^{19}$F CPMG RD |
| Exo10, Rrp44$^{PIN+CS}$, Rrp42$^{C59S, A106C-TFA}$ | $^{19}$F 1D, $^{19}$F CPMG RD |
| Exo10, 80mer RNA | activity assay |
| Exo10, Rrp42$^{\Delta 93-125}$, 80mer RNA | activity assay |
| Exo10, Rrp45-L, 80mer RNA | activity assay |
| Exo10, Rrp45-L, Rrp42$^{\Delta 93-125}$, 80mer RNA | activity assay |

**Table S7: Sequences of RNAs used in this study.**

| RNA | Sequence | Internal database reference |
|---|---|---|
| 46mer | GGAGGAGAGGUGAGGAGAGAGGAGAGGAAGGAAGGGAAGAAAGAAG | #72 |
| 80mer | GGAAGGAGAGGAAGGAAAGGUGGGAAGAGGAAGGAGAGGAGGGAAG AAAGAAGAGGAGAGGAAGGAAGGGAAGAAAGAAG | #39 |

**Table S8: Parameters for CPMG relaxation dispersion experiments.**

| System | $\nu_{CPMG}$ | | number of | $T_{CPMG}$ (ms) |
|---|---|---|---|---|
| | min (Hz) | max (Hz) | frequencies | |
| Exo9 Rrp42$^{C59S,\ A106C\text{-}TFA}$ | 100 | 5000 | 25 | 20 |
| Exo10 PIN Rrp42$^{C59S,\ A106C\text{-}TFA}$ | 250 | 5000 | 20 | 8 |
| Exo10 PIN+CS Rrp42$^{C59S,\ A106C\text{-}TFA}$ | 250 | 5000 | 20 | 8 |
| Exo9 Rrp42$^{C59S,\ A106C\text{-}TFA}$ + RNA | 100 | 5000 | 25 | 20 |
| Exo10 Rrp42$^{C59S,\ A106C\text{-}TFA}$ + RNA | 100 | 5000 | 25 | 20 |
| Exo9 Rrp41$^{D113tfmF}$ | 500 | 5000 | 10 | 4 |
| Exo10 Rrp41$^{D113tfmF}$ | 500 | 5000 | 10 | 2 |
| Exo9 Rrp41$^{Q86tfmF}$ | 167 | 5000 | 20 | 6 |
| Exo10 Rrp41$^{Q86tfmF}$ | 167 | 5000 | 20 | 6 |

**Table S9: Elution program for HPLC runs of activity assay samples.**

| Time interval (min) | Linear changes in concentration of buffer F (%) |
|---|---|
| 0 – 3 | 0 |
| 3 – 4 | 0 – 20 |
| 4 – 11 | 20 – 50 |
| 11 – 12 | 50 – 100 |
| 12 – 14 | 100 |
| 14 – 19 | 0 |

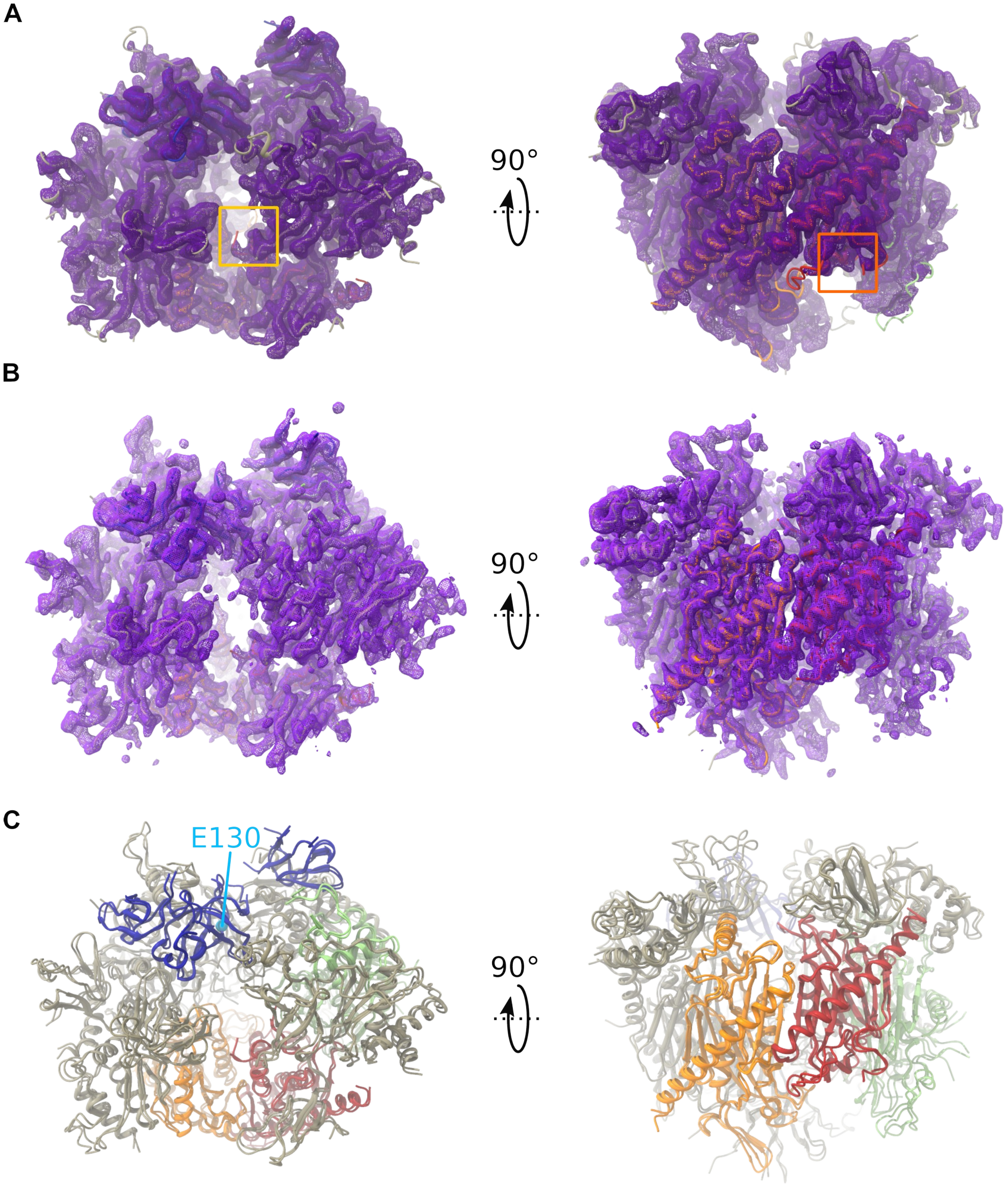
A
90°
B
90°
C
E130
90°

D

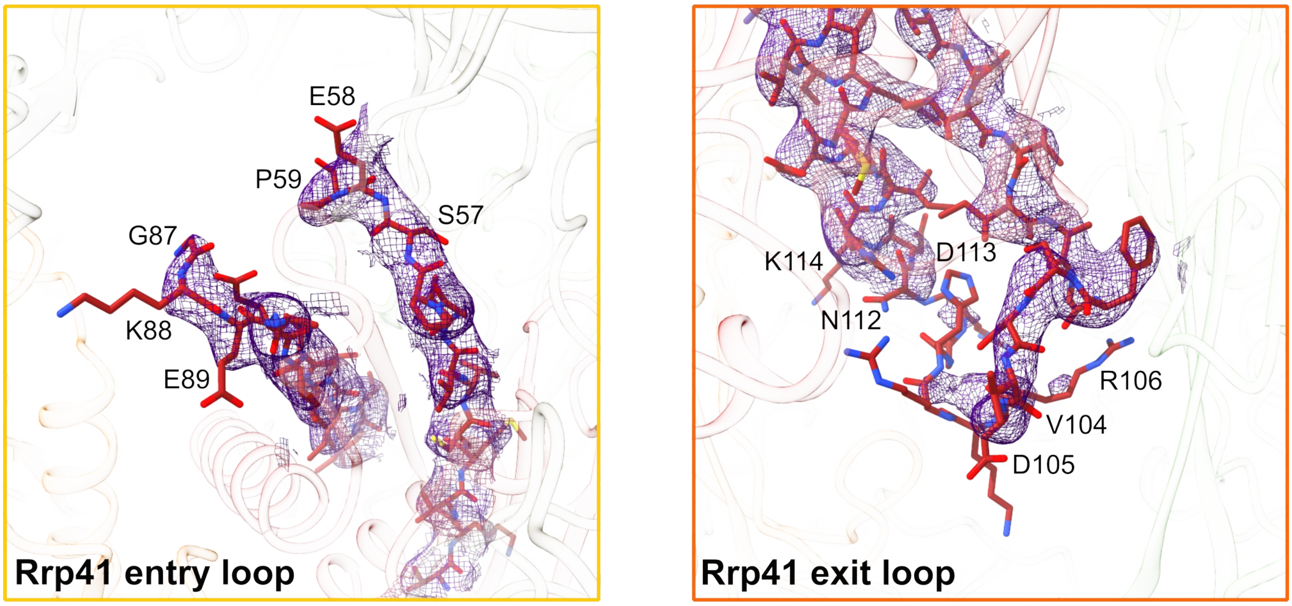

E

```
ctRrp41      51 CVVTGPSEPGPRRGTGAGTTGGGGAGGAGGGGSGGQGKEAEVV 93
cryo-EM         SSSS-----------------------------------SSS

ctRrp41      94 VSIVIAGFSSVDRKRHGRNDKRII 117
cryo-EM         SSSSS-------------HHHHH

ctRrp42      70 VKAEVEKTVSRSKEDEEVGLLVASAGDMDVDDEEGYAKVGADNRTGEASWVEITVE 125
cryo-EM         SSSSSSS-------------------------------------------SSSSS
```

(Bold in original: ctRrp41 residues GPRRGTGAGTTGGGGAGGAGGGGSGGQ and KRHGR; ctRrp42 residues VSRSKEDEEVGLLVASAGDMDVDDEEGYAKVGADNR.)

**Figure S1: Structure of the *C. thermophilum* exosome core ctExo9.** (**A**) Top view (left) and side view (right) of the cryo-EM density map at 3.2 rmsd. The close-ups in panel **D** are located inside the colored rectangles. (**B**) Top view (left) and side view (right) of the X-ray crystallography electron density map at 1 rmsd. (**C**) Overlay of the cryo-EM and X-ray crystallography structures. Csl4 is in blue, Rrp41 is in red, Rrp45 is in orange and Rrp42 is in green. Csl4 E130 is highlighted. (**D**) Cryo-EM density around the invisible section of the Rrp41 entry loop (left, yellow rectangle in panel **A**) and around the invisible section of the Rrp41 exit loop (right, orange rectangle in panel **A**), contoured at 3.1 rmsd. (**E**) Sequence and secondary structure obtained from the cryo-EM structure around the entry loop of Rrp41 (top), the exit loop of Rrp41 (center) and Rrp42-EL (bottom). S = strand, H = helix, - = loop or invisible residue. Residues in bold are invisible in the cryo-EM structure.

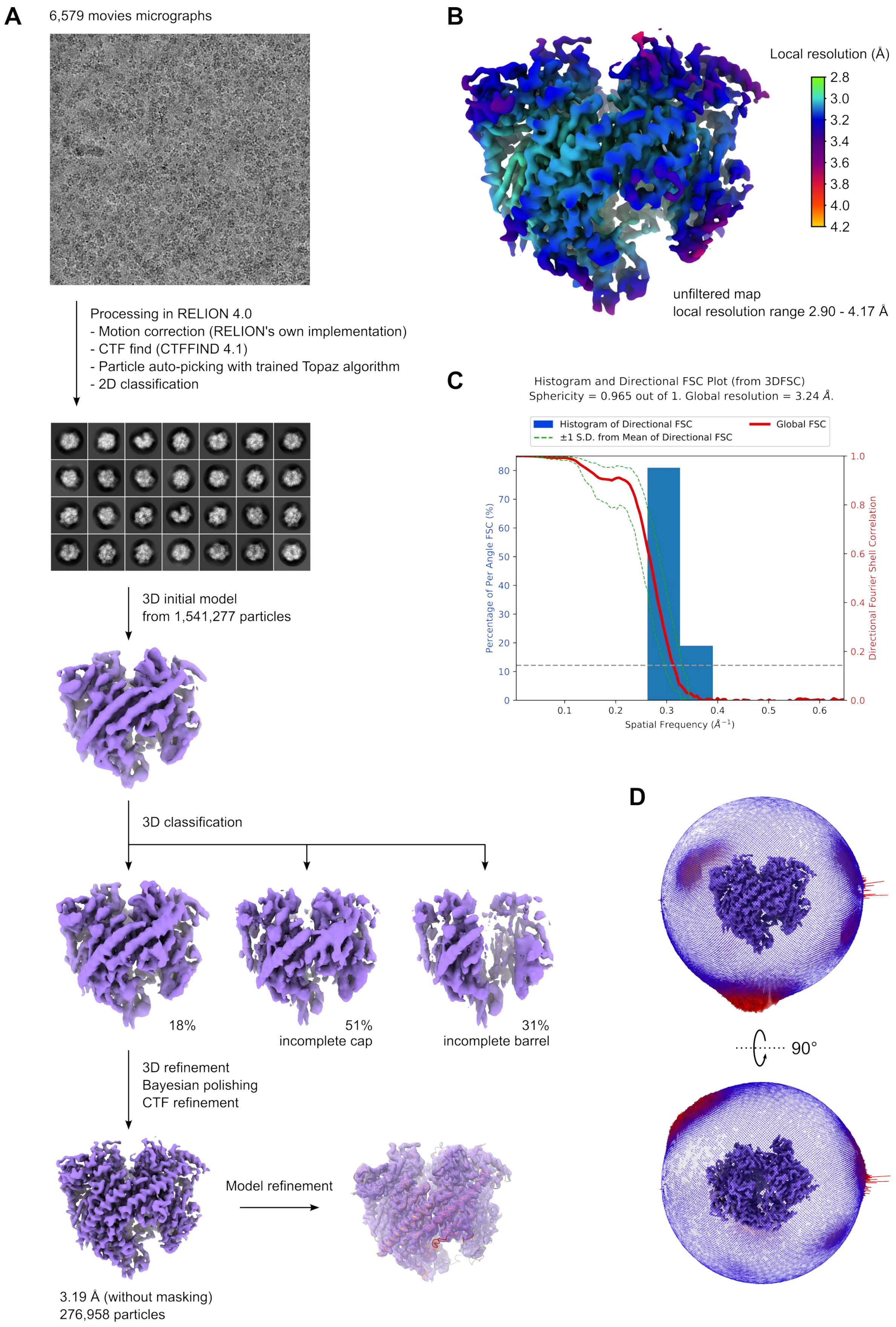
A
6,579 movies micrographs
Processing in RELION 4.0
- Motion correction (RELION's own implementation)
- CTF find (CTFFIND 4.1)
- Particle auto-picking with trained Topaz algorithm
- 2D classification
3D initial model
from 1,541,277 particles
3D classification
18%
51%
incomplete cap
31%
incomplete barrel
3D refinement
Bayesian polishing
CTF refinement
Model refinement
3.19 Å (without masking)
276,958 particles
B
Local resolution (Å)
2.8
3.0
3.2
3.4
3.6
3.8
4.0
4.2
unfiltered map
local resolution range 2.90 - 4.17 Å
C
Histogram and Directional FSC Plot (from 3DFSC)
Sphericity = 0.965 out of 1. Global resolution = 3.24 Å.
Histogram of Directional FSC
±1 S.D. from Mean of Directional FSC
Global FSC
Percentage of Per Angle FSC (%)
Directional Fourier Shell Correlation
Spatial Frequency (Å⁻¹)
0
10
20
30
40
50
60
70
80
0.1
0.2
0.3
0.4
0.5
0.6
0.0
0.2
0.4
0.6
0.8
1.0
D
90°

**Figure S2: Cryo-EM data processing.** (**A**) Scheme of the cryo-EM processing pipeline. The raw final density map of ctExo9, reconstructed from 276,958 particles, is shown at the bottom. The global resolution is 3.19 Å at FSC = 0.143. (**B**) Final Exo9 density map (as in **A**, bottom), contoured at 4 rmsd and colored according to the local resolution (calculated in RELION). (**C**) 3DFSC plot (*44*) calculated from the raw, unfiltered half maps. The sphericity of 0.965 indicates a very isotropic angular distribution. (**D**) Angular distribution of the particles used for the final reconstruction, plotted on the final map shown in panel **A**, bottom.

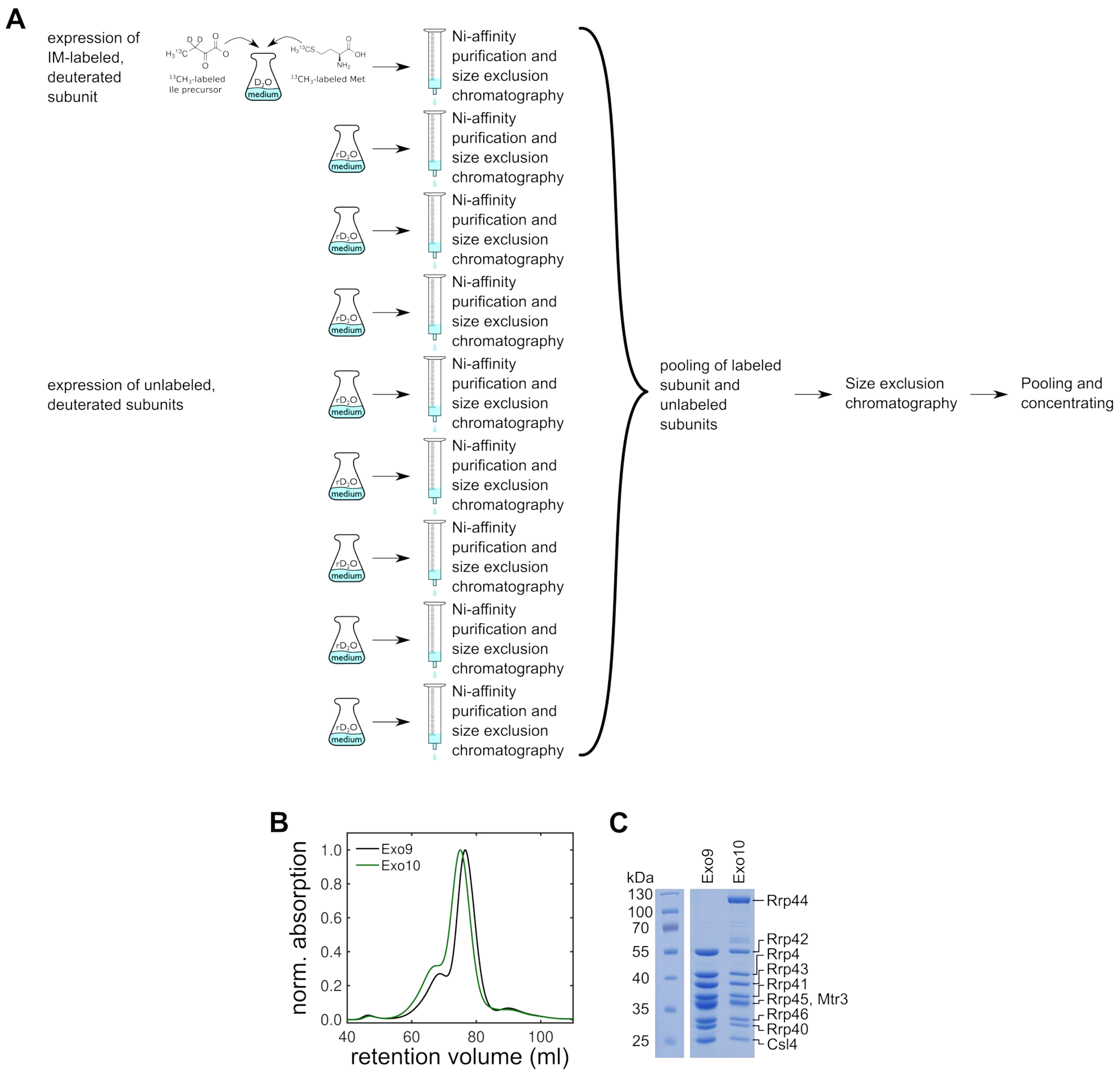

**Figure S3: Reconstitution of Exo9 and Exo10 complexes from individual subunits.** (**A**) Sample preparation to obtain modularly labeled exosome complexes (this example is for IM-labeling, reconstitutions of $^{19}$F or TEMPO spin-labeled subunits were performed analogously). The labeled subunit is expressed in $D_2O$ medium in the presence of appropriate precursors and purified. The unlabeled subunits are expressed individually (or as heterodimers) in ~95% $D_2O$ (r$D_2O$) medium and purified. The labeled subunit is pooled with the (8 or 9 other) unlabeled subunits followed by a size-exclusion purification step. Peak fractions are pooled and concentrated. (**B**) Size exclusion chromatogram of the exosome complex reconstituted *in vitro*. Exo9 is in black and Exo10 is in green. The absorption was measured at 280 nm and is normalized to the maximum absorption of each trace. (**C**) SDS-PAGE gel of the pooled peak fractions of Exo9 and Exo10.

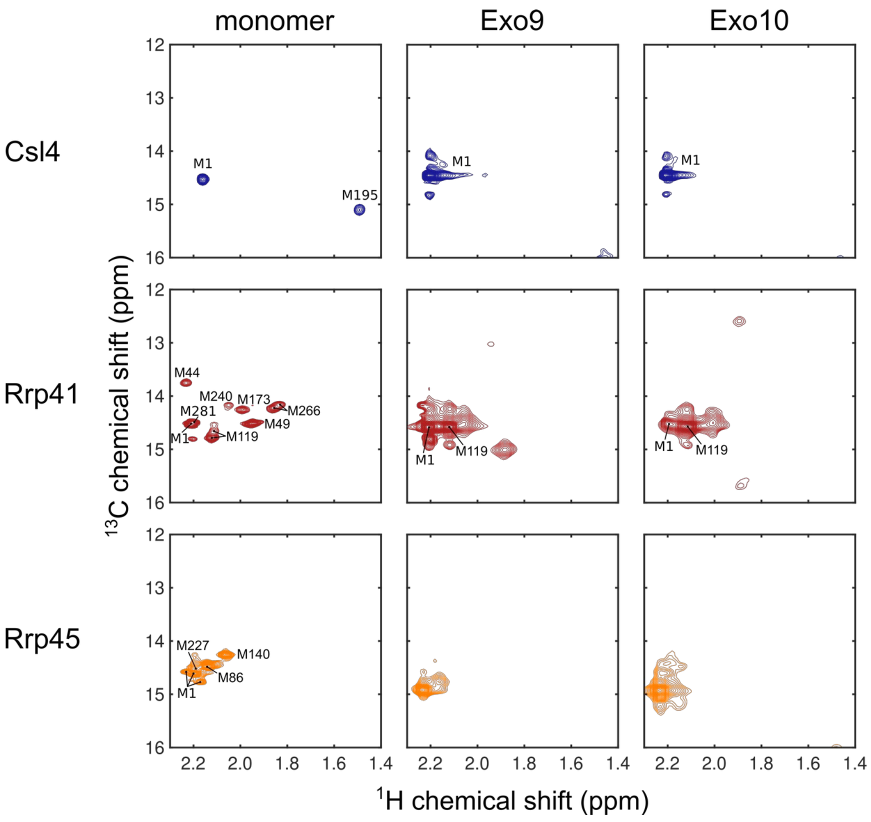

**Figure S4: Met-ε1 methyl-TROSY spectra.** Met-ε1 region of methyl-TROSY spectra for Csl4 (blue), Rrp41 (red) and Rrp45 (orange) as monomers (column 1) and when reconstituted into Exo9 (column 2) and Exo10 (column 3). Note, that the methionine resonances in the Exo9 and Exo10 complexes suffered from significant spectral overlap and were not suitable for the extraction of quantitative NMR parameters.

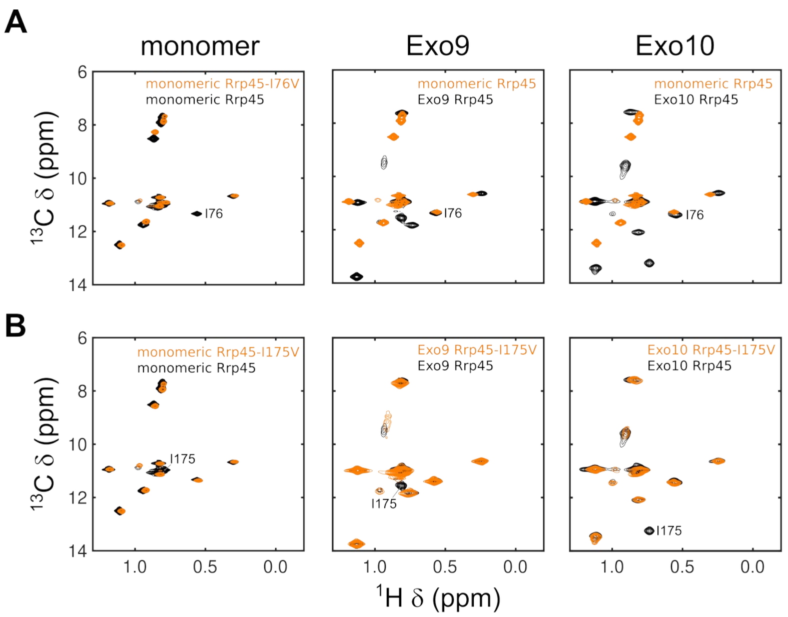

**Figure S5: Examples of assignment by point mutation.** (**A**) Assignment of a resonance (I76) that does not change chemical shift significantly between the monomeric form and the Exo9 or Exo10 complexes. Residue I76 of Rrp45 is assigned by comparing the methyl-TROSY spectrum of IM-labeled monomeric Rrp45 (left panel, black) to the spectrum of IM-labeled monomeric Rrp45-I76V (left panel, orange). Since the peak position of I76 is not strongly affected by reconstitution into Exo9 or Exo10, the assignment of monomeric Rrp45-I76 can be transferred to IM-labeled Rrp45 reconstituted into Exo9 (central panel, black) and Exo10 (right panel, black). IM-labeled monomeric Rrp45 is shown in orange in the central and right panel. (**B**) Assignment of a resonance (I175) that changes its chemical shift significantly upon formation of the Exo9 or Exo10 complexes. Residue I175 of Rrp45 is assigned by comparing the methyl-TROSY spectrum of IM-labeled monomeric Rrp45 (left panel, black) to the spectrum of IM-labeled monomeric Rrp45-I175V (left panel, orange). Since the peak position of I175 shifts substantially upon reconstitution into Exo9 (central panel, black) or Exo10 (right panel, black), the assignment in Exo9 and Exo10 is obtained by reconstituting IM-labeled Rrp45-I175V into Exo9 (central panel, orange) and Exo10 (right panel, orange).

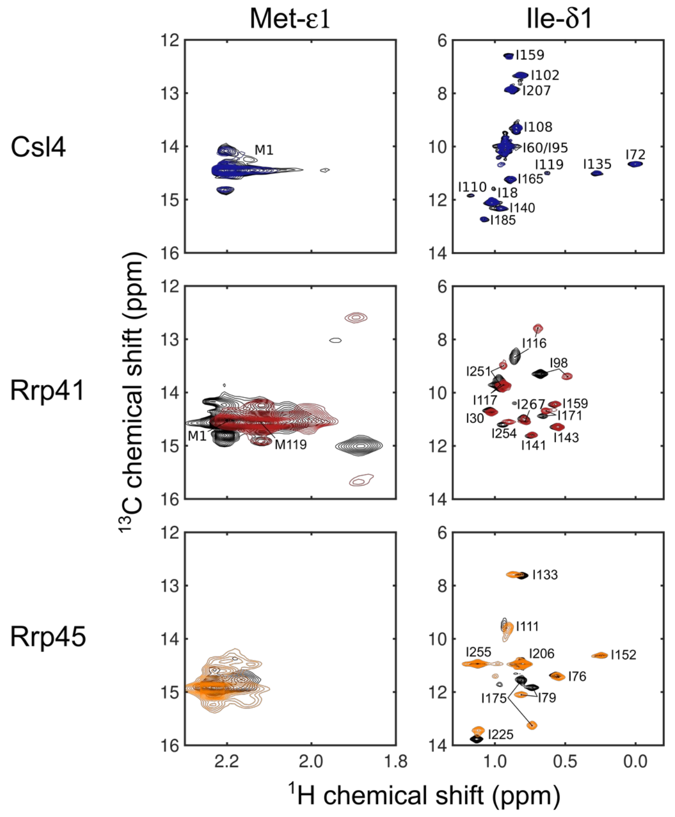

**Figure S6: Interaction between Exo9 subunits and Rrp44.** Overlay of the Met-ε1 (left) and Ile-δ1 (right) regions of methyl-TROSY spectra of IM-labeled Csl4 (blue), Rrp41 (red) and Rrp45 (orange) reconstituted into Exo9 (black) and Exo10 (in color).

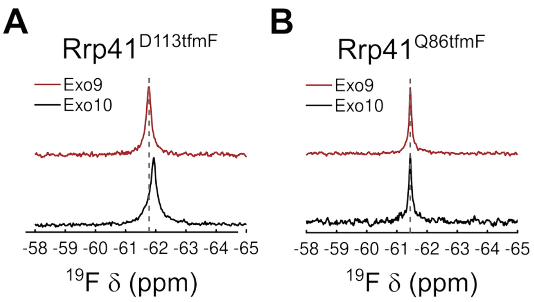

**Figure S7: Interaction of the exit and entry loop of Rrp41 with Rrp44.** (**A**) 1D $^{19}F$ spectrum of Rrp41$^{D113tfmF}$ (exit loop) and (**B**) 1D $^{19}F$ spectrum of Rrp41$^{Q86tfmF}$ (entry loop) reconstituted into Exo9 (red) and Exo10 (black). The dashed line indicates the center of the resonance for Exo9. The exit loop senses the presence of Rrp44.

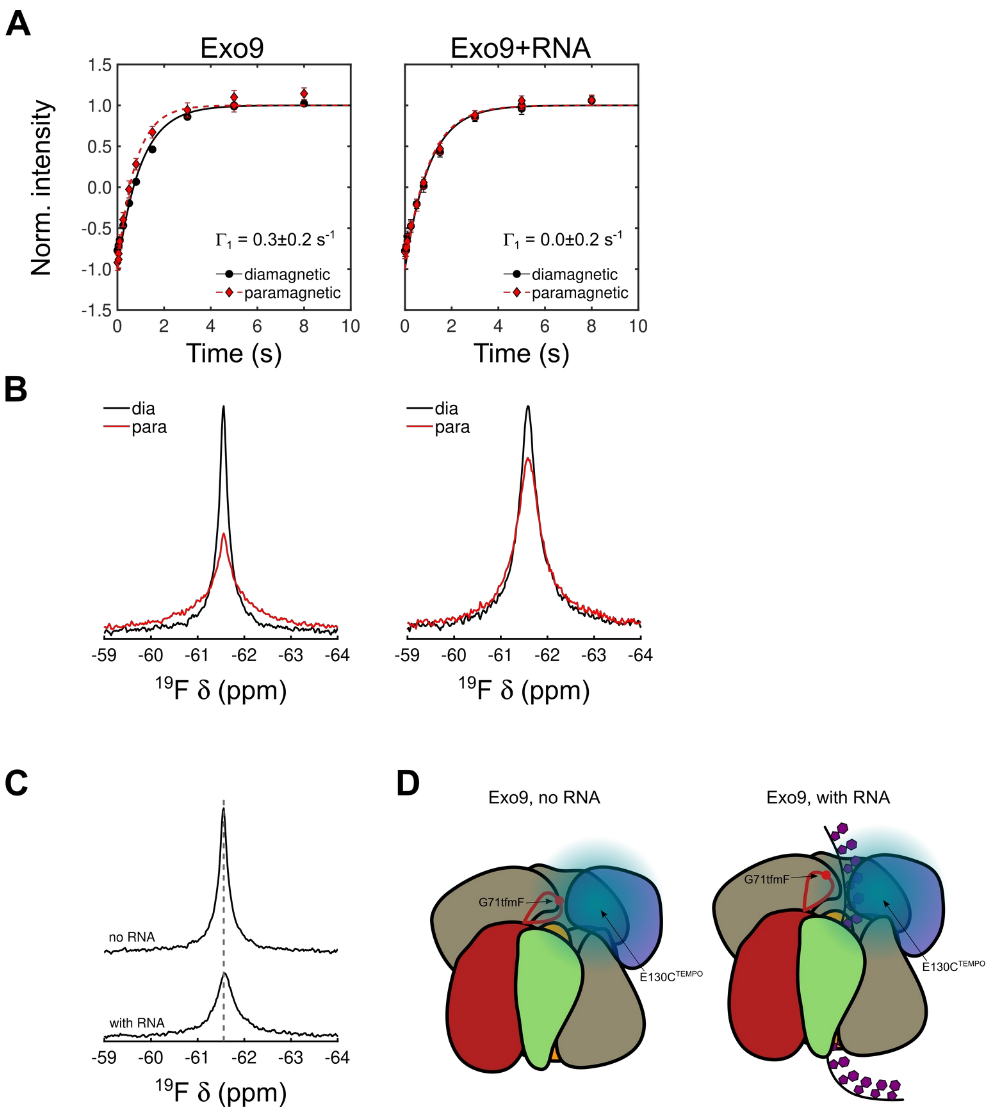

**Figure S8: Location and RNA interactions of the entry loop of Rrp41. (A)** $T_1$ inversion recovery experiments measured for Rrp41$^{G71tfmF}$ Csl4$^{C122S, E130C\text{-}TEMPO}$ in Exo9 without (left) and with (right) RNA for paramagnetic (in red) and diamagnetic (black) samples. $\Gamma_1$ PREs derived from Eq. S1 and Eq. S3 are indicated in the panels (see table S3A). (**B**) $^{19}$F 1D experiments measured for Rrp41$^{G71tfmF}$ Csl4$^{C122S, E130C\text{-}TEMPO}$ in Exo9 without (left) and with (right) RNA for paramagnetic (in red) and diamagnetic (black) samples. (**C**) Overlay of the diamagnetic 1D $^{19}$F spectra from panel **B** without (top) and with (bottom) RNA. The dotted line indicates the center of the resonance in the absence of RNA. (**D**) Schematic of the position of the entry loop of Rrp41 (red, the position of G71tfmF is shown as a red dot) with respect to Csl4 (blue, the PRE effect of the TEMPO spin-label at E130C is shown as a blue sphere). See also fig. S1C. In the absence of RNA the loop approaches Csl4 (left); in the presence of RNA the loop is displaced away from Csl4 (right). Rrp42 is shown in green and Rrp45 is depicted in orange.

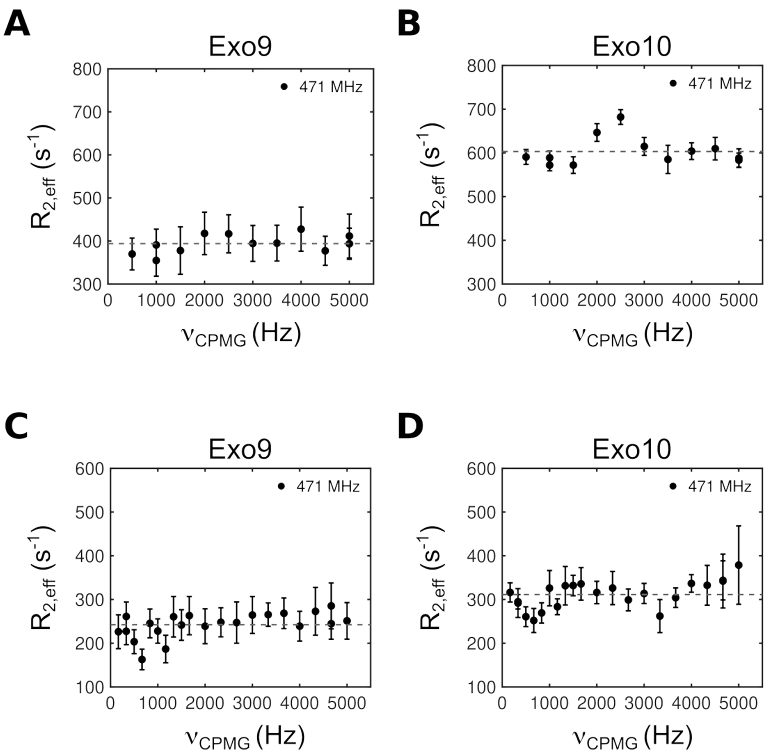

**Figure S9: Relaxation dispersion experiments of the exit and entry loop.** $^{19}$F CPMG relaxation dispersion profiles for Rrp41$^{D113tfmF}$ in (**A**) Exo9 and (**B**) Exo10 and CPMG relaxation dispersion profile for Rrp41$^{Q86tfmF}$ in (**C**) Exo9 and (**D**) Exo10.

**A**

```
ctRrp41      51 CVVTGPSEPGPRRGTGAGTTGGGGAGGAGGGGSGGQGKEAEVV 93
ssRrp41      56 AAVYGPKEMHPRHLSLP-------------------DRAVLR 75
scRrp41      51 TLVKGPKEPRLKSQMDT-------------------SKALLN 73
hsEXOSC4     50 AVVYGPHEIRGSRARAL-------------------PDRALVN 73
```

**B**

```
ctRrp41      94 VSIVIAGFSSVDRKR-HGRNDKRII 117
ssRrp41      76 VRYHMTPFSTDERKN--PAPSRREI 101
scRrp41      74 VSVNITKFSKFERSKSSHKNERRVL  98
hsEXOSC4     74 CQYSSATFSTGERKRRP-HGDRKSC  97
```

**C**

```
ctRrp45      99 TEVLLSRLLEKTIR 112
ssRrp42     106 NAIELARVVDRSLR 119
scRrp45     100 DEVLCSRIIEKSVR 113
hsEXOSC9     98 LLVKLNRLMERCLR 111
```

**Figure S10: Sequence alignments of ctRrp41 and ctRrp45.** Sequence alignment of (**A**) the entry loop of ctRrp41, (**B**) the exit loop of ctRrp41 and (**C**) a pore-facing helix of ctRrp45 with homologs from the archaeon *Sulfolobus solfataricus* (ssRrp41/ssRrp42), *S. cerevisiae* (scRrp41/scRrp45) and human (hsEXOSC4/hsEXOSC9). Amino acids that have previously been shown to coordinate RNA in *S. solfataricus* or *S. cerevisiae* (*45*, *46*) and corresponding amino acids in *C. thermophilum* and human are in bold. Italicized residues are not visible in the cryo-EM structure. Positions G71, Q86 and D113, at which a tfmF-label was introduced, are highlighted in red, and Ile residues that show CSPs upon RNA addition to the exosome are shown in cyan.

```
ctRrp42        70 VKAEVEKTVSRSKEDEEVGLLVASAGDMDVDDEEGYAKVGADNRTGEASWVEITVE 125
hsEXOSC7       62 VKAEMGTPKLEKPNEGYLEFFVDCS-------------------------ASATPE  92
scRrp42        62 IKSKVVDHHV---E---------------------------------NELLQVDVD  81
conservation      :*:::        :                                     .   :
```

**Figure S11: Conservation of Rrp42-EL.** Sequence alignment of an extended loop in ctRrp42 (Rrp42-EL) with the human (hsEXOSC7) and *S. cerevisiae* (scRrp42) homologs.

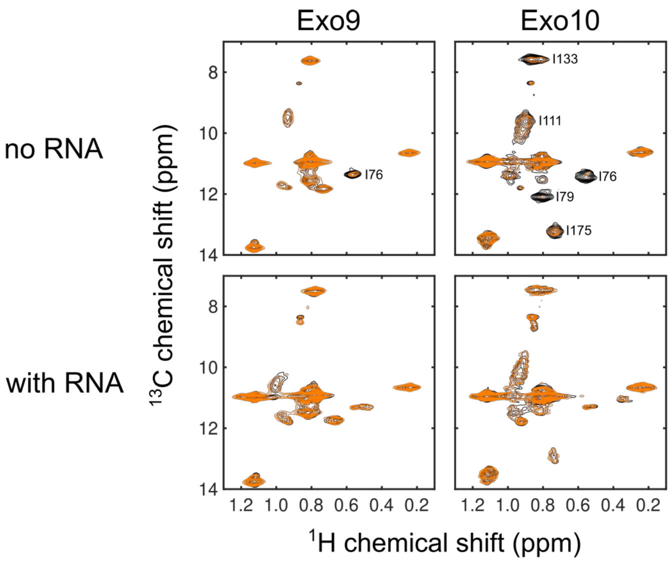

**Figure S12: Methyl PREs of Rrp42$^{C59S, A106C\text{-}TEMPO}$.** PREs for Ile-δ1 of Rrp45 in Exo9 and Exo10 with and without RNA. The spectrum of the diamagnetic sample is shown in black and the spectrum of the paramagnetic sample is depicted in orange. Resonances of residues that exhibit a substantial PRE ($I_{para}/I_{dia}$ < 0.7) are labeled.

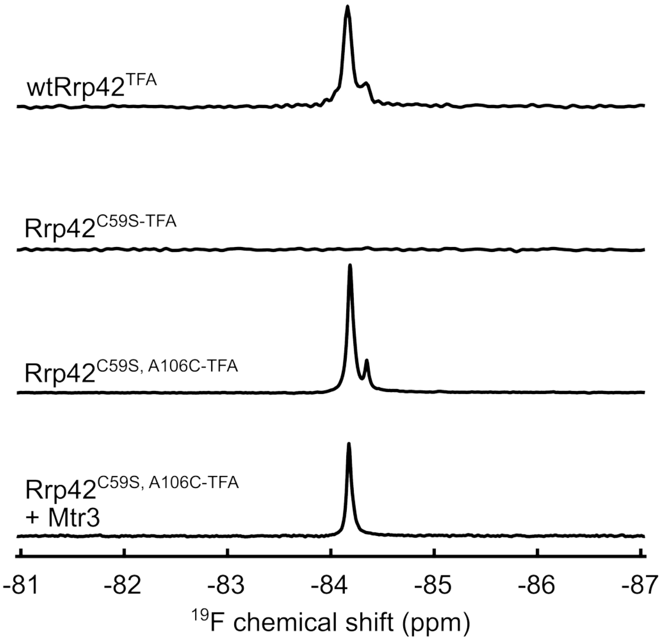

**Figure S13: TFA-labeling of Rrp42 mutants.** $^{19}$F spectra of TFA-labeled wtRrp42 (row 1), Rrp42$^{C59S}$ (row 2), Rrp42$^{C59S, A106C}$ (row 3) and Rrp42$^{C59S, A106C}$ in complex with Mtr3 (row 4).

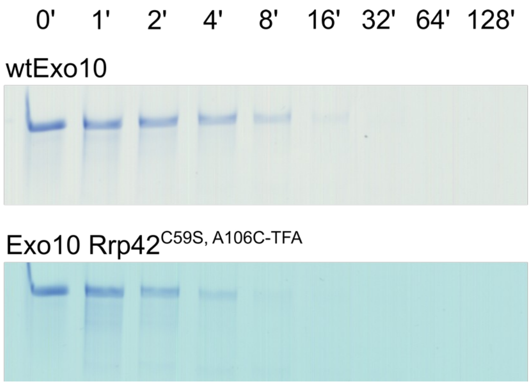

**Figure S14: Activity assay of Exo10 variants.** Activity assay of wtExo10 (top) and Exo10 Rrp42$^{C59S, A106C-TFA}$ (bottom) using an 80mer RNA (see table S7). The RNA substrate is degraded with highly similar rates in wtExo10 complex and the Exo10 complex that is $^{19}$F labeled in Rrp42-EL.

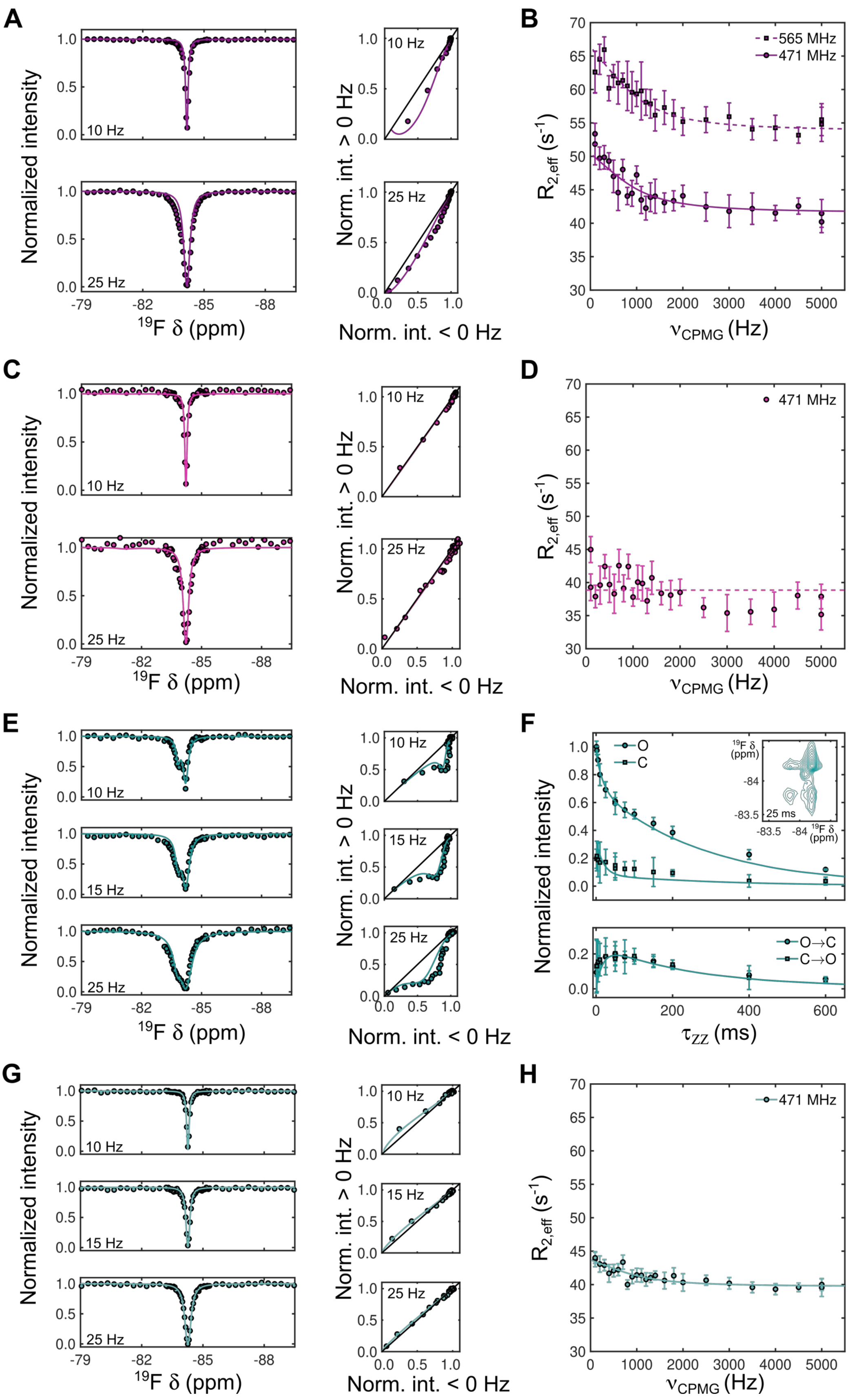
A
B
C
D
E
F
G
H
Normalized intensity
10 Hz
15 Hz
25 Hz
$^{19}$F δ (ppm)
-79
-82
-85
-88
Norm. int. > 0 Hz
Norm. int. < 0 Hz
$R_{2,eff}$ (s$^{-1}$)
$\nu_{CPMG}$ (Hz)
565 MHz
471 MHz
O
C
O→C
C→O
$\tau_{ZZ}$ (ms)
25 ms
$^{19}$F δ (ppm)

**Figure S15: Exchange dynamics of Rrp42-EL.** (**A**) and (**B**): Dynamics of Rrp42$^{C59S,\ A106C\text{-}TFA}$ in Exo9. (**A**) CEST experiments (left panels) and symmetry of the profile's dip (right panels). (**B**) CPMG RD profiles. (**C**) and (**D**): Dynamics of Rrp42$^{C59S,\ A106C\text{-}TFA}$ in Exo9 with RNA. (**C**) CEST experiments (left panels) and symmetry of the profile's dip (right panels). (**D**) CPMG RD profile. (**E**) and (**F**): Dynamics of Rrp42$^{C59S,\ A106C\text{-}TFA}$ in Exo10. (**E**) CEST experiments (left panels) and symmetry of the profile's major dip (right panels). (**F**) EXSY profiles for diagonal (open state 'O', closed state 'C') and cross ($O \rightarrow C$, $C \rightarrow O$) peaks. The inset shows the $^{19}$F/$^{19}$F correlation spectrum for $\tau_{ZZ}$ = 25 ms. (**G**) and (**H**): Dynamics of Rrp42$^{C59S,\ A106C\text{-}TFA}$ in Exo10 with RNA. (**G**) CEST experiments (left panels) and symmetry of the profile's dip (right panels). (**H**) CPMG RD profile. A two-site exchange model as described in materials and methods was globally fitted to the data in panels **A**, **B**, **E** and **F**. Fitted parameter values are shown in table S4.

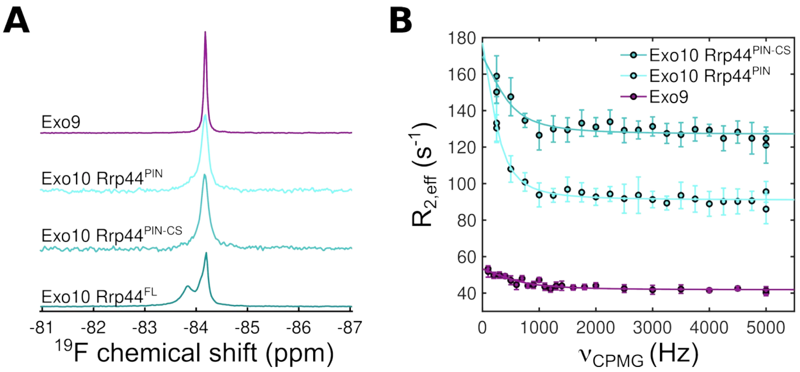

**Figure S16: Interaction of Rrp42-EL with Rrp44.** (**A**) $^{19}$F spectra of Rrp42$^{C59S,\ A106C\text{-}TFA}$ in Exo9, Exo10 Rrp44$^{PIN}$, Exo10 Rrp44$^{PIN+CS}$ and Exo10 with full-length Rrp44. (**B**) CPMG relaxation dispersion profiles for Exo10 Rrp44$^{PIN+CS}$, Exo10 Rrp44$^{PIN}$ and Exo9. Note that CPMG relaxation dispersion experiments on the Exo10 complex (with full-length Rrp44) suffered from very fast relaxation rates that prevented us from obtaining high quality fluorine NMR data.

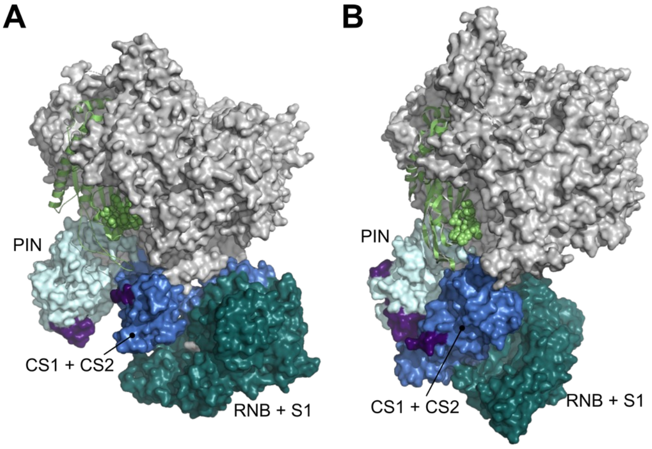

**Figure S17: Localization of Rrp42-EL with respect to Rrp44 in yeast and human exosome complexes.** Structures of (**A**) the human (PDB ID: 6H25 (*47*)) and (**B**) the *S. cerevisiae* (PDB ID: 4IFD (*48*)) exosome. Exo9 is shown in gray surface representation, except for Rrp42, which is displayed in green cartoon. Residues in the loop corresponding to ctRrp42-EL (see fig. S11) are green and shown in spherical representation. Rrp44 domains are displayed in colored surface representation: the PIN domain is in light blue, the CS1 and CS2 domains are in marine and the RNB and S1 domains are in dark teal. The structurally resolved sections of the connecting loop between the PIN and CS1 domain is highlighted in violet. The CS1 and CS2 domains come close to Rrp42, and are kept in place there due to interactions between the bottom of the Exo9 barrel and the Rrp44 RNB-S1 domains. See also fig. S24.

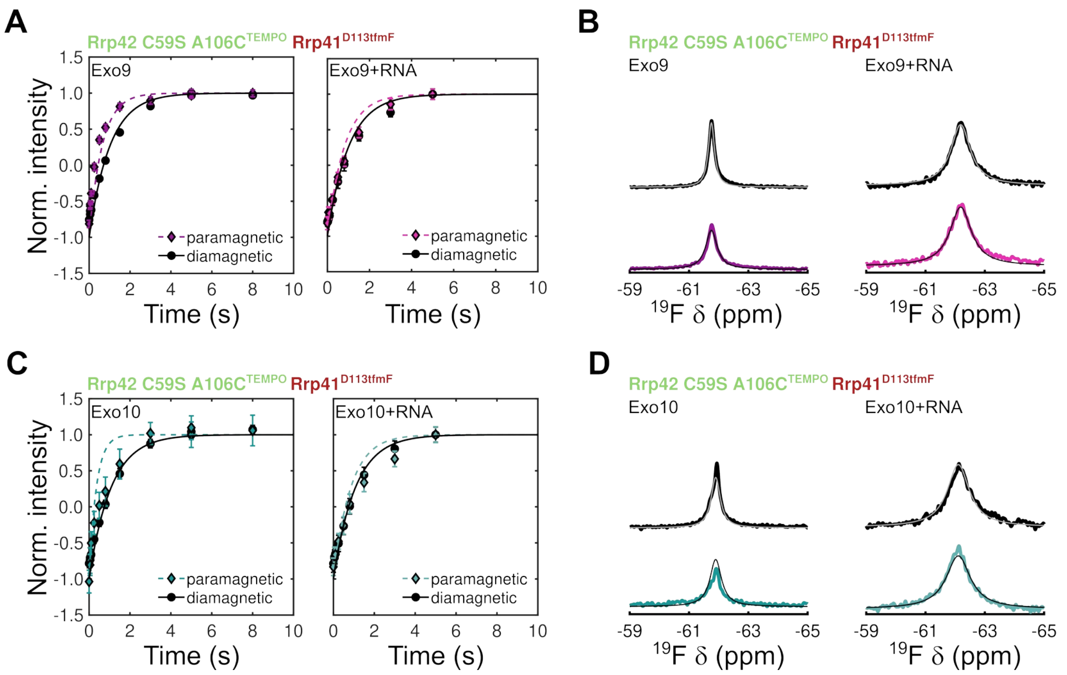

**Figure S18: Fast loop dynamics of Rrp42-EL.** (**A**) Inversion recovery experiments and (**B**) $^{19}$F 1D spectra for Rrp41$^{D113tfmF}$ in Exo9 Rrp42$^{C59S, A106C-TEMPO}$ without (left panel) and with (right panel) RNA for paramagnetic (in color) and diamagnetic (black) samples. (**C**) Inversion recovery experiments and (**D**) $^{19}$F 1D spectra for Rrp41$^{D113tfmF}$ in Exo10 Rrp42$^{C59S, A106C-TEMPO}$ without (left panel) and with (right panel) RNA for paramagnetic (in color) and diamagnetic (black) samples. A two-site exchange model as described in materials and methods was globally fitted to the data. Fitted parameter values are shown in table S4.

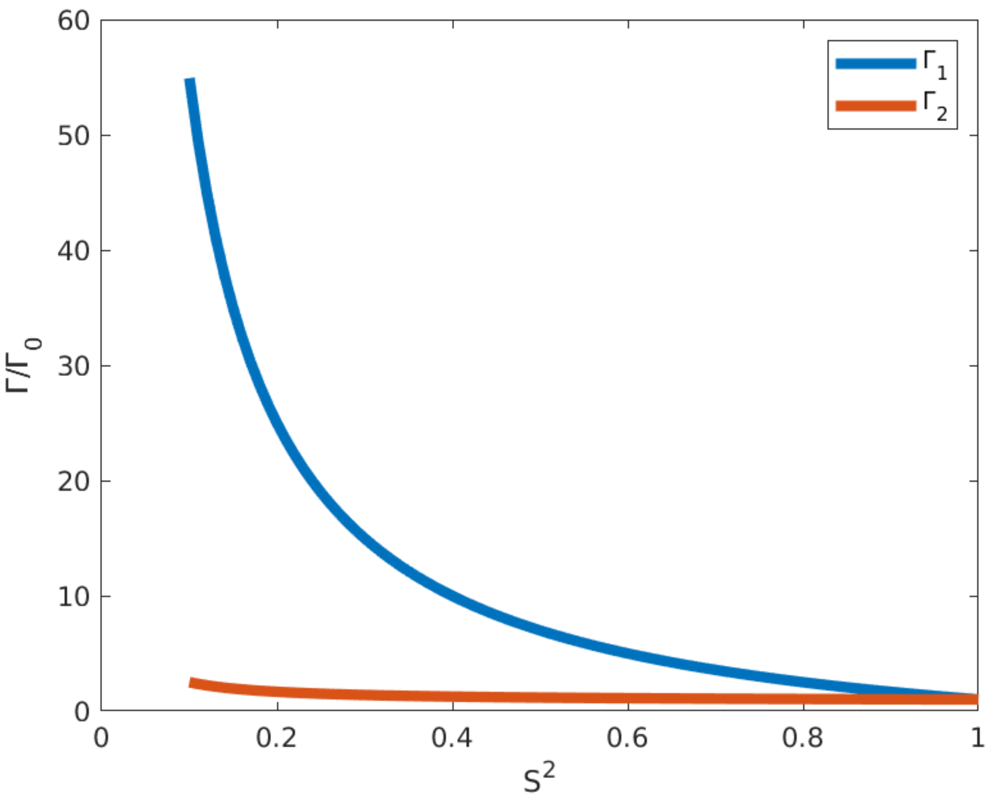

**Figure S19: Dependency of the PRE rates $\Gamma_1$ and $\Gamma_2$ on the order parameter $S^2$.** $\Gamma_{(1;\ \text{blue or}\ 2;\text{red})}$ are calculated based on equation 9 (using $\tau_i$ = 10 ns, $\tau_r$ = 100 ns and using $\omega = 2*\pi*470*10^6$ rad/s; corresponding to the fluorine resonance frequency on a 500 MHz NMR spectrometer). $\Gamma_0$ is the relaxation rate for $S^2 = 1$. The plot indicates that $\Gamma_1$ depends strongly on $S^2$, whereas $\Gamma_2$ is largely uneffected by the order parameter.

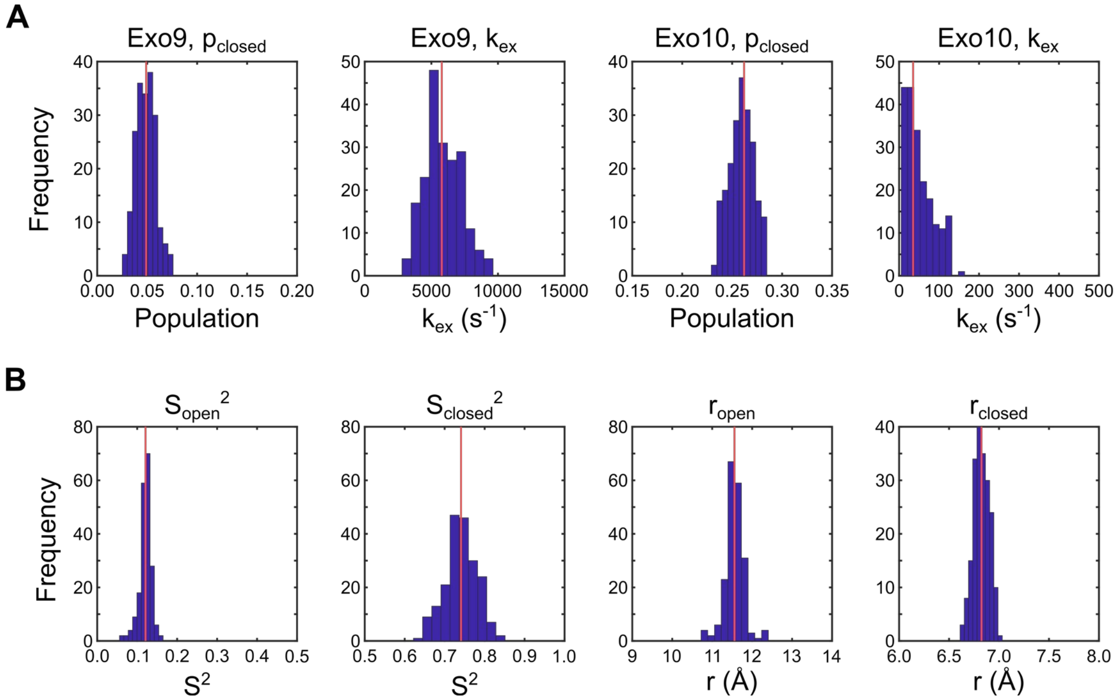

**Figure S20: Monte-Carlo simulations of dynamics data.** 200 datasets were simulated and the model was fitted to these artificial datasets to extract parameter uncertainties as described in material and methods. The histograms show the distribution of fitted values (**A**) for the population of the closed state ($p_{closed}$) and exchange rate ($k_{ex}$) in Exo9 and Exo10 and (**B**) for the order parameter ($S^2$) and distance ($r$) between the $^{19}$F label (Rrp41$^{D113tfmF}$) and the paramagnetic center of Rrp42$^{C59S,A106C\text{-}TEMPO}$ of the open and closed state of Rrp42-EL. The red line indicates the parameter value obtained from the best fit of the two-site exchange model to the experimental data.

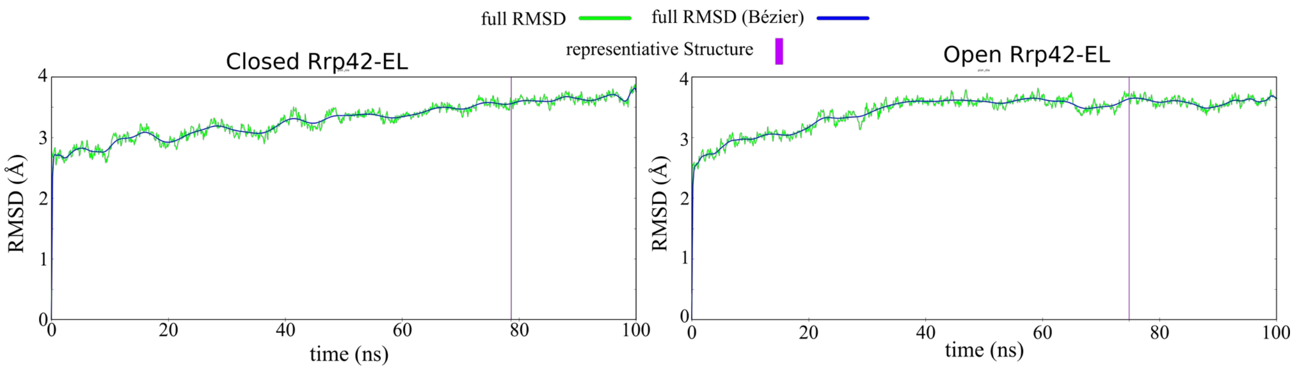

**Figure S21: C$^{\alpha}$ root mean square deviation (RMSD).** The plot of the RMSD of the C$^{\alpha}$ carbon atoms over the course of the MD simulation of Rrp42-EL closed (left) and open state (right) with respect to the MD starting structure converges towards a plateau. Therefore, both RMSD plots indicate that an energetically equilibrated structure is reached. Highlighted by the magenta line is the simulation frame that constitutes the representative structure of the equilibrated part of each simulation.

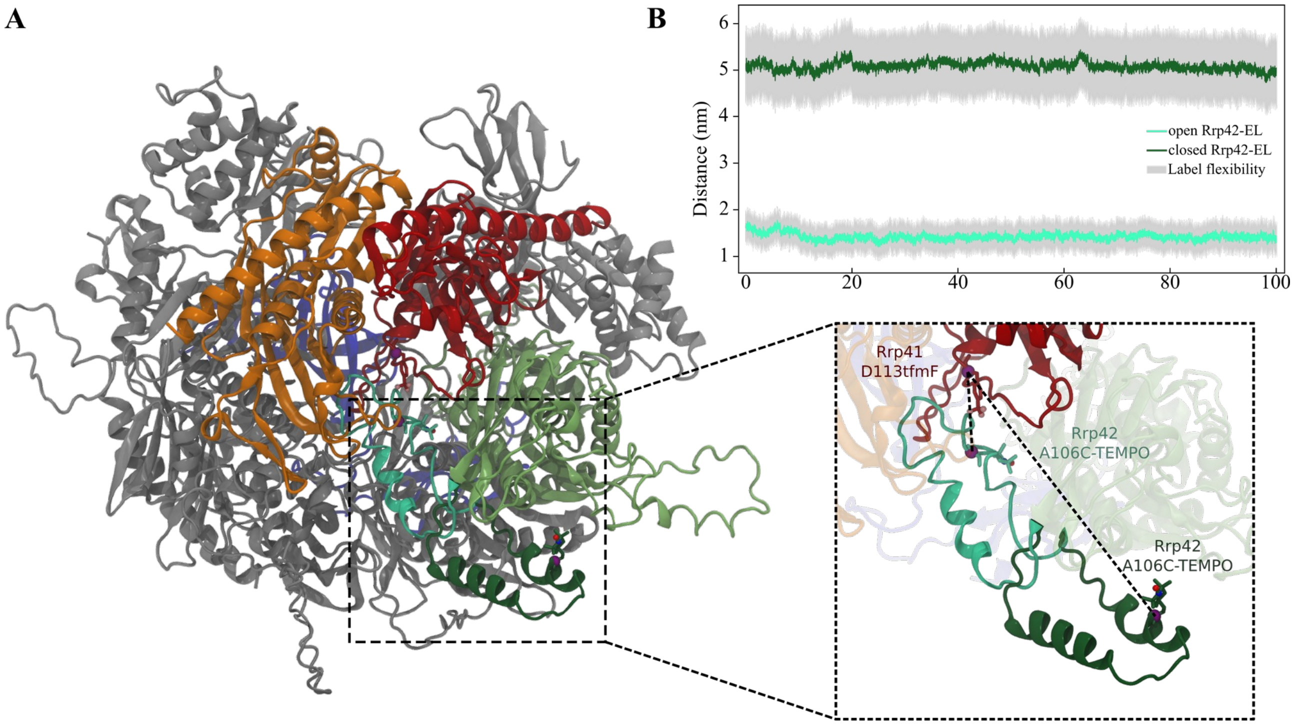

**Figure S22: Distances between the residues labeled for $^{19}$F PRE experiments.** (**A**) Location of the $^{19}$F and spin label within the modeled starting structures of Exo9 with open state (dark green) and closed state (cyan) Rrp42-EL. Rrp42 is shown in green, Rrp41 in red, Rrp45 in orange and Csl4 in blue. Zoom: Distance of the $C^{\alpha}$ carbon atoms of the labeled residues within the starting model of the MD simulation. (**B**) The MD simulations were performed without the spin labels. Therefore, the distance between the $C^{\alpha}$ carbon atoms of Rrp42-A106 and Rrp41-D113 of the MD simulation of the open (cyan) and closed (dark green) conformation of the Rrp42-EL is plotted. This distance approximates the distances between the labels measured in the NMR experiments (Fig. 4, Fig S18). The internal dynamics of the label is roughly approximated by the gray area.

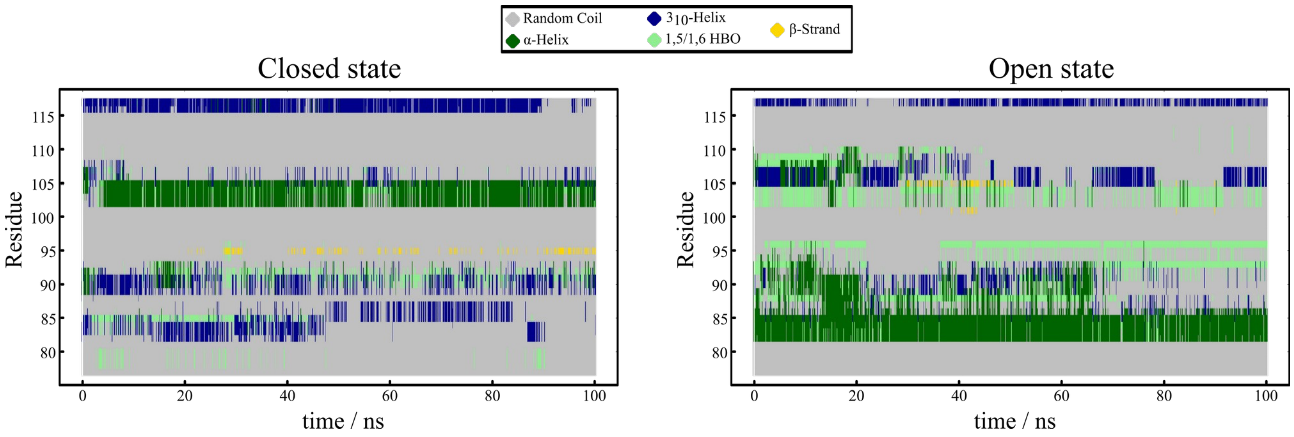

**Figure S23: Comparison of the secondary structure of Rrp42-EL within the MD simulations of the closed and open state.** Each residue of Rrp42-EL is colored with respect to its secondary structure over the time course of the MD simulation with a step size of 0.1 ns. The plot differentiates between random coil (gray), α-helix (dark green), $3_{10}$-helix (blue), β-strand (yellow) and 1,5 or 1,6 hydrogen bonds (light green). The separated first column shows the secondary structure of the starting structure obtained from the AlphaFold prediction. The MD simulations indicate that parts of the predicted α-helices are not stable within the simulations. They might fluctuate between α-helix and random coil or there might be less α-helical content than predicted.

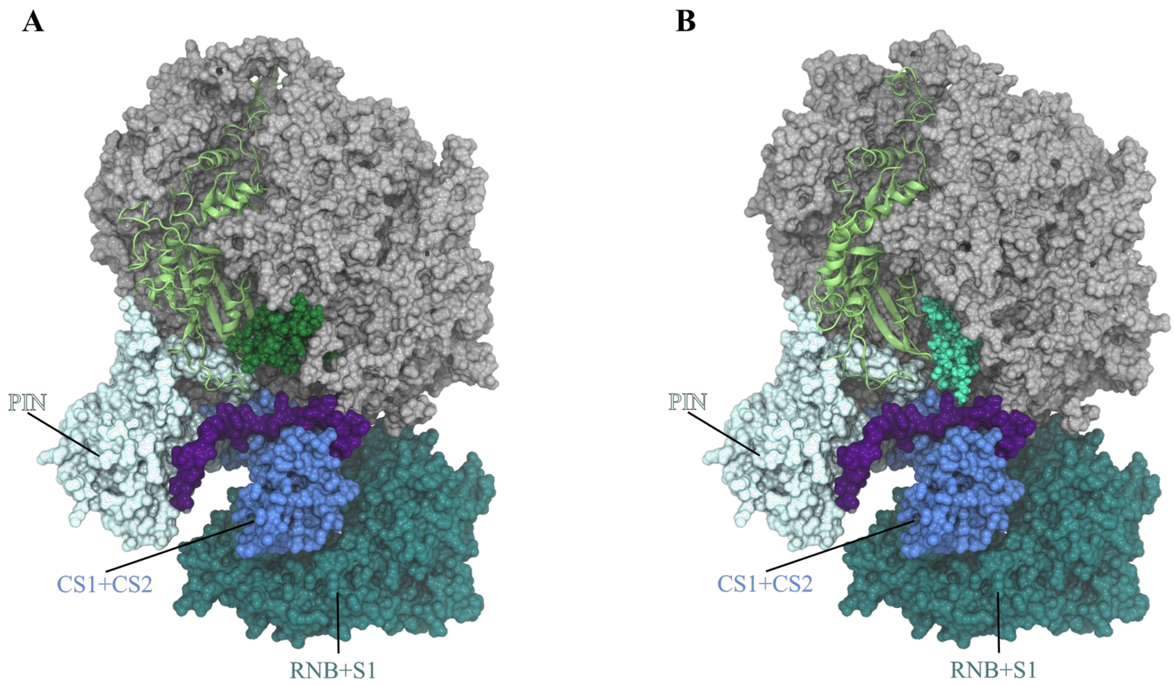

**Figure S24: Localization of Rrp42-EL with respect to Rrp44.** Superposition of the representative open (**A**) and closed (**B**) state of Rrp42-EL from ctExo9 complemented with Rrp44 from *S. cerevisiae* (chain J, PDB: 4IFD (*48)*). Exo9 is shown in surface representation in grey, Rrp42 as cartoon green, open Rrp42-EL in dark-green and the closed loop in green, each displayed as spheres. Rrp44 is shown in surface representation, the PIN domain is colored light blue, CS1 and CS2 are colored marine, the RNB and S1 domains are shown in dark teal. The connective loop between the PIN and CS1 domain is highlighted in violet. See also fig. S17.

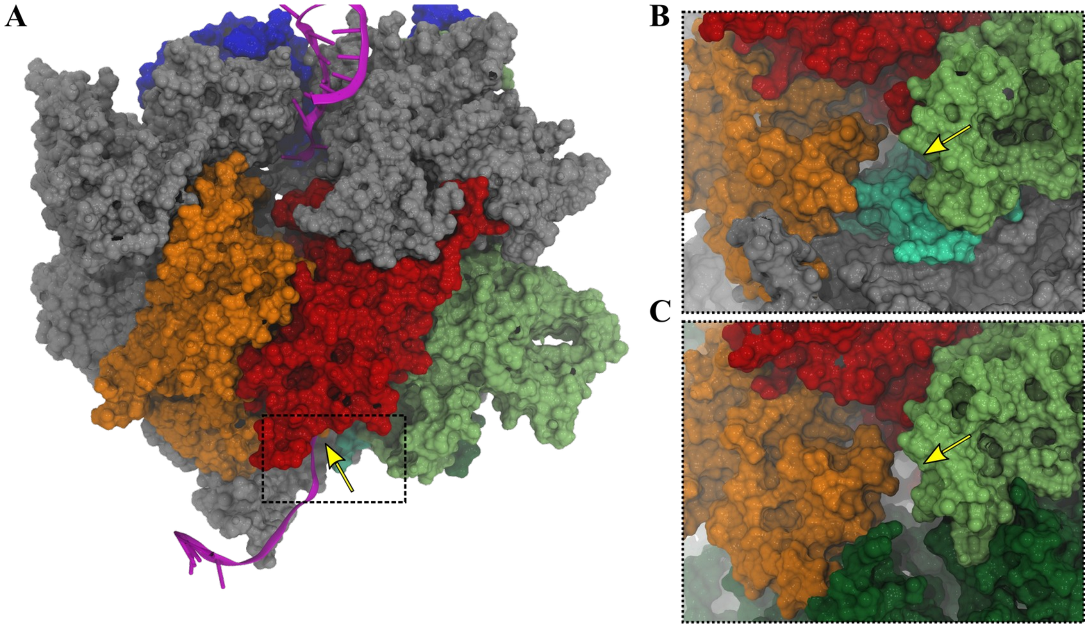

**Figure S25: The closed state of Rrp42-EL blocks the RNA channel.** (**A**) The RNA (purple) is modeled into the representative structures of the open and closed state of the Exo9 MD simulations. RNA positioning was guided by the X-Ray structure of *S. cerevisiae* Exo9 (PDB ID: 4IFD (*48*)). (**B**) The enlargement of the exit tunnel with removed RNA illustrates the blocking of Rrp42-EL in the closed state (cyan) as indicated by the arrow. (**C**) In the open state the exit tunnel is free as the Rrp42-EL (dark green) is remote from the exit site of the RNA channel. Subunit coloring: Csl4: blue, Rrp41: red, Rrp45:  orange, Rrp42: green.

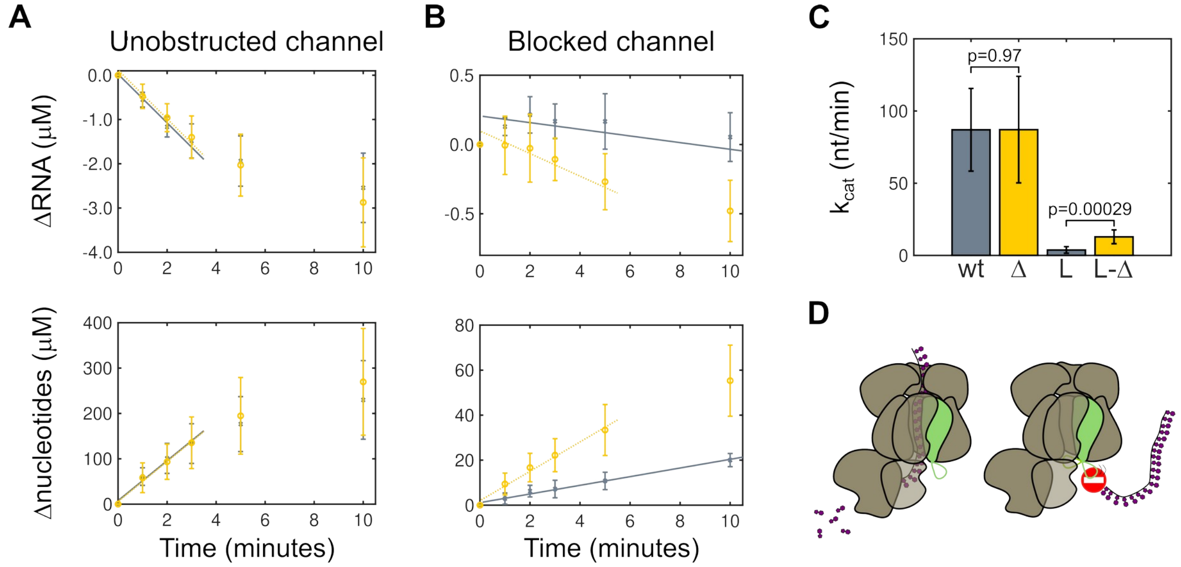

**Figure S26: Effect of Rrp42-EL on the activity of Exo10.** HPLC-based activity assays using an 80mer RNA (see table S7) (**A**) for wtExo10 (gray crosses, solid line) and Exo10 Rrp42$^{\Delta 93\text{-}125}$ (yellow circles, dashed line) and (**B**) for channel-blocked Exo10 Rrp45-L (gray crosses, solid line) and Exo10 Rrp45-L Rrp42$^{\Delta 93\text{-}125}$ (yellow circles, dashed line). The lines are global linear fits to the linear activity regime, the points that are not on the straight fitted line were considered to be outside the linear degradation regime and not used in the determination of the degradation rates. The exosome can display non-linear degradation rates due to e.g. product inhibition caused by the high concentration of nucleotides that arise, or due to the reduction in the substrate concentration during the reaction. (**C**) Catalytic activity of exosome constructs. wt = wtExo10, Δ = Exo10 Rrp42$^{\Delta 93\text{-}125}$, L = Exo10 Rrp45-L, L-Δ = Exo10 Rrp45-L Rrp42$^{\Delta 93\text{-}125}$. p-values are derived from a paired-sample *t*-test. (**D**) Rrp42-EL allows on-path RNA to access Rrp44 for degradation (left) but blocks a direct access path towards Rrp44 (right).

**A**

```
ctRrp45       79 IATELSPMTSPTFEVNRPT-ETEV 101
scRrp45       79 ISTEISPMAGSQFENGNITGEDEV 102
hsEXOSC9      78 FNLELSQMAAPAFEPGRQS-DLLV 100
conservation     :  *:* *:.  ** .. : :  *
```

**B**

```
ctRrp45       79 IATELSPMTSPTFEVNRPTETEV 101
cryo-EM          SSSSS-------------HHHHH
```

**C**

```
scRrp45       79 ISTEISPMAGSQFENGNITGEDEV 102
PDB 4IFD:A       SS----HHH-----------HHHH
```

**Figure S27:** (**A**) Sequence alignment of a channel facing loop in Rrp45 with the *S. cerevisiae* (scRrp45) and human (hsEXOSC9) homologs. The loop extension was introduced between residues N94 and R95 highlighted in bold. (**B**) Secondary structure of the loop from ctRrp45 based on the cryo-EM structure. (**C**) Secondary structure of the loop from scRrp45 obtained from PDB ID: 4IFD, chain A. S = strand, H = helix, - = unstructured.

8. D. Liebschner, P. V. Afonine, M. L. Baker, G. Bunkóczi, V. B. Chen, T. I. Croll, B. Hintze, L.-W. Hung, S. Jain, A. J. McCoy, N. W. Moriarty, R. D. Oeffner, B. K. Poon, M. G. Prisant, R. J. Read, J. S. Richardson, D. C. Richardson, M. D. Sammito, O. V. Sobolev, D. H. Stockwell, T. C. Terwilliger, A. G. Urzhumtsev, L. L. Videau, C. J. Williams, P. D. Adams,

Macromolecular structure determination using X-rays, neutrons and electrons: recent developments in Phenix. *Acta Crystallogr. D* **75**, 861–877 (2019).

16. R. L. J. Keller, *The Computer Aided Resonance Assignment Tutorial* (CANTINA Verlag, Goldau, Switzerland, 2004; cara.nmr.ch).

17. A. J. Baldwin, An exact solution for $R_{2,eff}$ in CPMG experiments in the case of two site chemical exchange. *J. Magn. Reson.* **244**, 114–124 (2014).

18. P. Vallurupalli, A. Sekhar, T. Yuwen, L. E. Kay, Probing conformational dynamics in biomolecules via chemical exchange saturation transfer: a primer. *J. Biomol. NMR* **67**, 243–

271 (2017).

39. W. G. Hoover, Canonical dynamics: Equilibrium phase-space distributions. *Phys. Rev. A* **31**,

1695–1697 (1985).
